# Human T-bet governs innate and innate-like adaptive IFN-γ immunity against mycobacteria

**DOI:** 10.1101/2020.08.31.274589

**Authors:** Rui Yang, Federico Mele, Lisa Worley, David Langlais, Jérémie Rosain, Ibithal Benhsaien, Houda Elarabi, Carys A. Croft, Jean-Marc Doisne, Peng Zhang, Marc Weisshaar, David Jarrossay, Daniela Latorre, Yichao Shen, Jing Han, Conor Gruber, Janet Markle, Fatima Al Ali, Mahbuba Rahman, Taushif Khan, Yoann Seeleuthner, Gaspard Kerner, Lucas T. Husquin, Julia L. Maclsaac, Mohamed Jeljeli, Fatima Ailal, Michael S. Kobor, Carmen Oleaga-Quintas, Manon Roynard, Mathieu Bourgey, Jamila El Baghdadi, Stéphanie Boisson-Dupuis, Anne Puel, Fréderic Batteux, Flore Rozenberg, Nico Marr, Qiang Pan-Hammarström, Dusan Bogunovic, Lluis Quintana-Murci, Thomas Carroll, Cindy S Ma, Laurent Abel, Aziz Bousfiha, James P. Di Santo, Laurie H Glimcher, Philippe Gros, Stuart G Tangye, Federica Sallusto, Jacinta Bustamante, Jean-Laurent Casanova

## Abstract

Inborn errors of human IFN-γ immunity underlie mycobacterial disease. We report a patient with mycobacterial disease due to an inherited deficiency of the transcription factor T-bet. This deficiency abolishes the expression of T-bet target genes, including *IFNG*, by altering chromatin accessibility and DNA methylation in CD4^+^ T cells. The patient has profoundly diminished counts of mycobacterial-reactive circulating NK, invariant NKT (iNKT), mucosal-associated invariant T (MAIT), and Vδ2^+^ γδ T lymphocytes, and of non-mycobacterial-reactive classic T_H_1 lymphocytes, the remainders of which also produce abnormally low amounts of IFN-γ. Other IFN-γ-producing lymphocyte subsets however develop normally, but with low levels of IFN-γ production, with exception of Vδ2^−^ γδ T lymphocytes, which produce normal amounts of IFN-γ in response to non-mycobacterial stimulation, and non-classic T_H_1 (T_H_1*) lymphocytes, which produce IFN-γ normally in response to mycobacterial antigens. Human T-bet deficiency thus underlies mycobacterial disease by preventing the development of, and IFN-γ production by, innate (NK) and innate-like adaptive lymphocytes (iNKT, MAIT, and Vδ2^+^ γδ T cells), with mycobacterial-specific, IFN-γ-producing, purely adaptive αβ T_H_1* cells unable to compensate for this deficit.

## Introduction

In the course of primary infection, life-threatening disease in otherwise healthy children, adolescents, and even adults, can result from monogenic inborn errors of immunity, which display genetic heterogeneity and physiological homogeneity (Casanova, 2015b, 2015a). Mendelian susceptibility to mycobacterial disease (MSMD) is characterized by a selective, inherited predisposition to clinical disease caused by weakly virulent mycobacteria, such as *Mycobacterium bovis* Bacille Calmette-Guérin (BCG) vaccines and environmental mycobacteria (Rosain et al., 2019). Patients are also vulnerable to *bona fide* tuberculosis. Patients with typical, “isolated” MSMD are rarely prone to other infectious agents, with the exception of *Salmonella* and occasionally other intra-macrophagic bacteria, fungi, and parasites (Bustamante et al., 2014; Rosain et al., 2019). Patients with atypical, “syndromic” MSMD often display other clinical phenotypes, infectious or otherwise. MSMD, both “isolated” and “syndromic”, displays a high level of genetic heterogeneity, with causal mutations in 15 genes, and additional allelic heterogeneity, resulting in 30 different disorders (**Table S1**). However, there is also physiological homogeneity, as all known genetic causes of MSMD affect IFN-γ-dependent immunity (Boisson-Dupuis et al., 2018; Bustamante et al., 2014; Kong et al., 2018; Martínez-Barricarte et al., 2018; Rosain et al., 2019). Mutations of *IL12B*, *IL12RB1*, *IL12RB2*, *IL23R*, *TYK2*, *ISG15*, *RORC*, *IKBKG* (NEMO), *IRF8*, and *SPPL2A* impede IFN-γ production by innate and adaptive immune cells, whereas mutations of *IFNGR1, IFNGR2*, *STAT1*, *JAK1*, and *CYBB* impair cellular responses to IFN-γ (**Fig. S1A**). The clinical penetrance and severity of MSMD depend strongly on genetic etiology and they increase with decreasing levels of IFN-γ activity (Dupuis et al., 2000). Four etiologies also result in an impairment of immunological circuits other than the IFN-γ circuit, accounting for “syndromic” MSMD in the corresponding patients (Rosain et al., 2019). Collectively, these studies revealed the crucial role of human IFN-γ in antimycobacterial immunity and its redundancy for immunity against many other pathogens.

The cellular basis of MSMD in patients with impaired responses to IFN-γ involves mononuclear phagocytes. The ability of these cells to contain the ingested mycobacteria depends on their activation by IFN-γ (Nathan et al., 1983). The cellular basis of MSMD in patients with impaired IFN-γ production is poorly understood, as most types of lymphocytes can produce IFN-γ (Wilson and Schoenborn, 2007). Some genetic etiologies of MSMD affect some lymphocyte subsets more than others. ISG15 deficiency preferentially impairs the production of IFN-γ by NK cells (Bogunovic et al., 2012; Zhang et al., 2015). IL-12Rβ2 deficiency preferentially impairs the production of IFN-γ by NK, B, γδ T, classic αβ T cells, ILC1, and ILC2 cells, whereas IL-23R deficiency preferentially impairs that by invariant NKT (iNKT) and mucosal-associated invariant T (MAIT) cells, and both IL-12Rβ1 and TYK2 deficiencies impair both the IL-12- and IL-23- dependent subsets (Boisson-Dupuis et al., 2018; Martínez-Barricarte et al., 2018). SPPL2a and IRF8 deficiencies selectively impair the production of IFN-γ by CD4^+^ CCR6^+^ T_H_1 (T_H_1*) cells (Hambleton et al., 2011; Kong et al., 2018), a T_H_1 cell subset enriched in *Mycobacterium*-specific effector cells, whereas CCR6^−^ T_H_1 cells do not respond to mycobacteria (Acosta-Rodriguez et al., 2007). RORγ/RORγT deficiency impairs the development of iNKT and MAIT cells, and also decreases the production of IFN-γ by γδ T and αβ T_H_1* cells (Okada et al., 2015). Interestingly, although the lack of both αβ T and γδ T cells in SCID patients underlies BCG disease (Casanova et al., 1995), most, if not all other deficits of antigen-specific αβ T-cell responses, whether affecting only CD4^+^ or CD8^+^ T cells, such as HLA-II or HLA-I deficiency, typically do not (**Table S2**). Moreover, selective deficiencies of NK or iNKT cells do not confer a predisposition to mycobacterial disease (Casey et al., 2012; Cottineau et al., 2017; Gineau et al., 2012; Hughes et al., 2012; Latour, 2007; Locci et al., 2009; Morgan et al., 2011; Tangye et al., 2017). The nature of the IFN-γ-producing innate, innate-like adaptive, and purely adaptive lymphocyte subsets indispensable for antimycobacterial immunity, either alone or in combination, therefore remains largely unknown. No genetic cause has yet been identified for half the MSMD patients. We therefore sought to discover a new genetic etiology of MSMD that would expand the molecular circuit controlling human IFN-γ immunity while better delineating the cellular network involved.

## Results

### Identification of an MSMD patient homozygous for an indel variant of *TBX21*

We studied a three-year-old boy (“P”) born to first-cousin Moroccan parents (**Fig. 1A**). He suffered from disseminated BCG disease (BCG-osis) following vaccination at the age of three months. He also had persistent reactive airway disease (RAD), but was otherwise healthy (Supplementary Material - Case Report). He did not suffer from any other severe infectious diseases despite documented (VirScan) infection with various viruses, including Epstein-Barr virus (EBV), human cytomegalovirus (CMV), roseola virus, adenoviruses A, B, C and D, influenza virus A, rhinovirus A, and bacteria, such as *Streptococcus pneumoniae* and *Staphylococcus aureus* (**Fig. S1B**). We hypothesized that P had an autosomal recessive (AR) defect. We performed whole-exome sequencing (WES) on P, his unaffected brother, and both parents. Genome-wide linkage (GWL) analysis revealed 32 linked regions (LOD score >1.3 and size >500 kb) under a model of complete penetrance (**Fig. S1C**). In these linked regions, there were 15 rare homozygous non-synonymous or essential splicing variants in 15 different genes (minor allele frequency, MAF < 0.003 in gnomAD v2.1 and 1000 Genomes Project, including for each major ancestry) with a combined annotation-dependent depletion (CADD) score above their mutation significance cutoffs (MSC) (Consortium, 2015; Itan et al., 2016; Kircher et al., 2014; Zhang et al., 2018) (**Fig. S1D, Table S3**). After the exclusion of genes with other predicted loss-of-function (LOF) variants with a frequency greater than 0.5% in gnomAD, 12 candidate genes remained (**Table S3**). The c.466_471delGAGATGinsAGTTTA insertion and/or deletion (indel) variant of *TBX21* (T-box protein 21, or T-box, expressed in T cells, T-bet) was the candidate variant predicted to be the most damaging (Kircher et al., 2014). Moreover, based on connectivity to *IFNG*, the central gene of the entire network of all known MSMD-causing genes (Itan et al., 2013, 2014), *TBX21* was the most plausible candidate gene (**Table S4**). T-bet is a transcription factor that governs the development or function of several IFN-γ-producing lymphocytes in mice, including T_H_1 cells (Lazarevic et al., 2013; Szabo et al., 2000, 2002). These findings suggested that homozygosity for the rare indel variant c.466_471delGAGATGinsAGTTTA of *TBX21* is MSMD-causing. We investigated this variant according to the guidelines for genetic studies of single patients (Casanova et al., 2014). Sanger sequencing confirmed that P carried the indel variant in exon 1 of *TBX21* whereas his unaffected brother was homozygous wild-type (WT/WT) and both parents were heterozygous (WT/M) (**Fig. S1E**). A closely juxtaposed 12-nucleotide (nt) region identical to this variant sequence was detected 8-nt upstream from the variant, and may have served as a template for the generation of this variant (**Fig. S2A**). The variant did not alter the exon 1-exon 2 junction of the *TBX21* mRNA in EBV-transformed B (EBV-B) cells or peripheral blood mononuclear cells (PBMCs) (**Fig. S2B - D**). The variant present in P thus resulted in the replacement of E156 and M157, two amino acids that are highly conserved across different species and among other paralogs of T-box transcription factors, with S156 and L157 (**Fig. 1B** and **Fig. S3A**).

**Figure 1.**
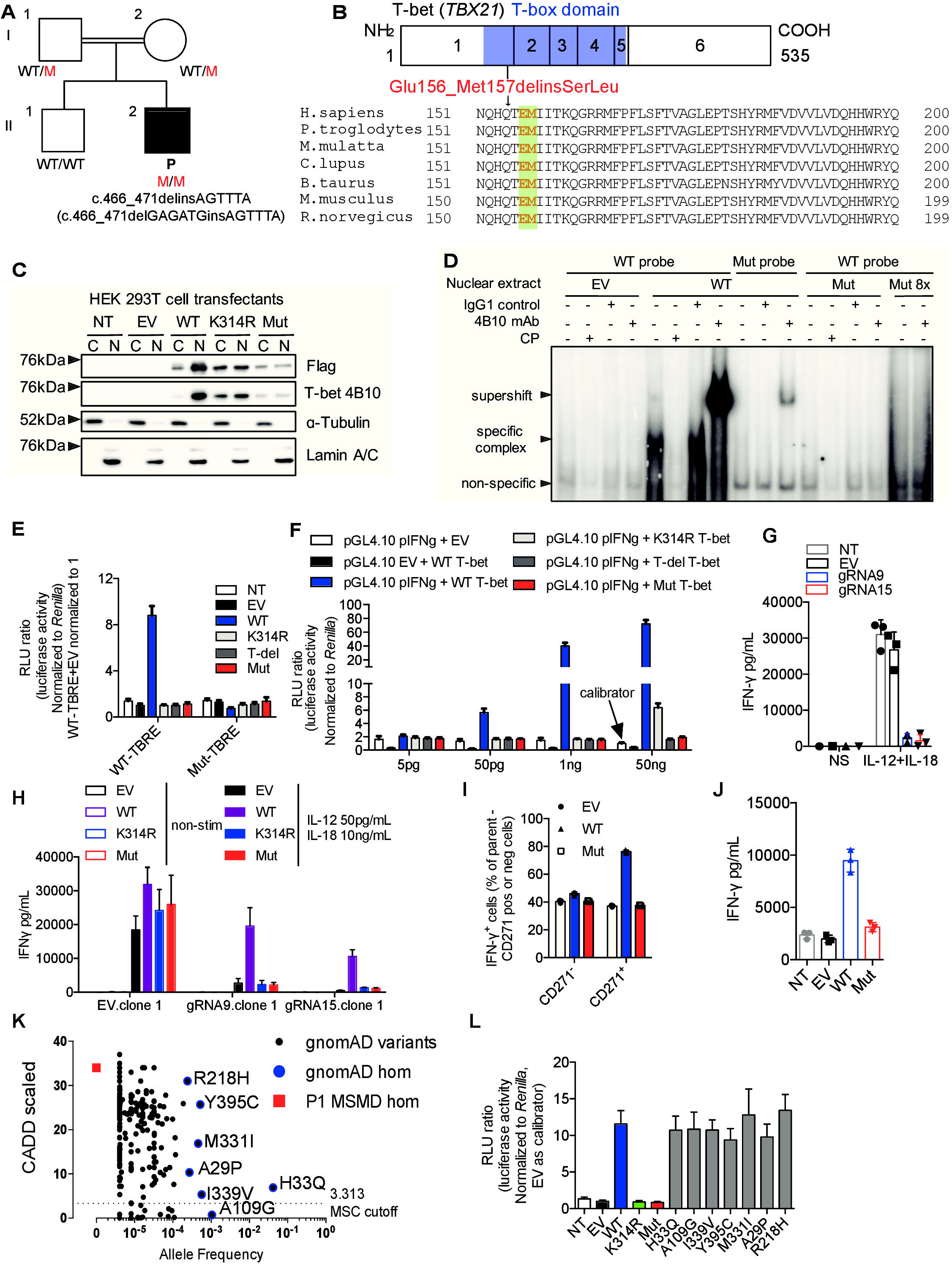

### Overexpressed mutant T-bet is LOF

We investigated the expression of the mutant allele (Mut), by overexpressing empty vector (EV), and vectors containing the WT or Mut allele, or negative controls with the T-box domain deleted (T-del) or K314R, the human ortholog of a known LOF mouse mutant, in human embryonic kidney (HEK) 293T cells (Jang et al., 2013). The production and nuclear translocation of Mut T-bet were impaired (**Fig. 1C** and **Fig. S3B**). When its DNA-binding activity to consensus T-box regulatory elements (TBRE) was assessed, nuclear proteins from WT-transfected cells bound the WT-TBRE but not the Mut-TBRE, and this specific complex was super-shifted by an anti-T-bet antibody (Ab) and inhibited by a competitor probe (CP). However, Mut T-bet did not bind WT-TBRE DNA (**Fig. 1D**). We also tested the ability of WT and Mut T-bet to induce a luciferase transgene under the control of the TBRE or human *IFNG* proximal promoter (Chen and Prywes, 1999; Janesick et al., 2012; Soutto et al., 2002; Tong et al., 2005a). WT T-bet induced high levels of luciferase activity with WT-TBRE but not with Mut-TBRE. Mut T-bet and the negative controls (T-del and K314R) did not induce luciferase activity (**Fig. 1E**). Mut T-bet was also LOF for transactivation of the *IFNG* promoter, whereas the negative control K314R was markedly hypomorphic (**Fig. 1F**). We investigated the amino-acid substitution responsible for the abolition of transcriptional activity. We tested the effects of the WT and Mut forms of T-bet and of T-bet forms with single-residue substitutions (E156S and M157L), or alanine substitutions (E156A and M157A) on T-bet protein production and transcriptional activity. The loss of E156 (E156S and E156A) abolished transcriptional activity, but the production of the T-bet protein was unaffected. By contrast, a loss of methionine residues (M157L and M157A) preserved transcriptional activity but decreased the levels of T-bet protein expression (**Fig. S3C** and **D**). T-bet is necessary for the optimal production of IFN-γ in NK, ILCs, γδ, and CD4^+^ T cells in mice and is sufficient for IFN-γ production in NK and CD4^+^ T cells in humans (Chen et al., 2007; Lazarevic et al., 2013; Powell et al., 2012; Szabo et al., 2002; Yu et al., 2006). We investigated the impact of the T-bet mutation on the induction of *IFNG* expression, by generating CRISPR/Cas9 gene-edited human NK-92 cell lines lacking *TBX21* (**Fig. S4A - D**). Upon stimulation with IL-12 + IL-18, these *TBX21* knockout (KO) NK-92 cells displayed a strong impairment of IFN-γ production (**Fig. 1G** and **Fig. S4E**). Following re-introduction of the WT or Mut *TBX21*-containing plasmid into *TBX21* KO NK-92 cells, the WT *TBX21* rescued IFN-γ production by *TBX21* KO cells, whereas the Mut *TBX21* did not (**Fig. 1H**, **Fig. S4F** and **G**). Finally, the overexpression of WT, but not of Mut T-bet increased IFN-γ levels to values above those for endogenous production in expanding naïve CD4^+^ T cells from healthy donors (**Fig. 1I** and **J**, **Fig. S4H**). Thus, Mut T-bet overexpression abolished DNA binding, and the mutant protein had no transactivation activity, failed to induce IFN-γ production in an NK cell line or primary CD4^+^ T cells, and can therefore reasonably be considered a LOF allele.

### *TBX21* variants in the general population are functional

The *TBX21* indel variant in P was not found in the gnomAD v2.1, Bravo, or Middle Eastern cohort databases, or our in-house database of more than 6,000 exomes, including > 1,000 individuals of North African origin (Karczewski et al., 2019; Scott et al., 2016; Taliun et al., 2019). The CADD score of 34 obtained for this allele is well above the MSC of 3.313 (**Fig. 1K**) (Itan et al., 2016; Kircher et al., 2014; Zhang et al., 2018). The *TBX21* gene has a low tolerance of deleterious variations, with a low gene damage index (GDI) score of 3.493 (Itan et al., 2015) and a low residual variation intolerance score (RVIS: −0.74) (Petrovski et al., 2015). Moreover, only three predicted LOF variants (variant: 17:45821662 T / TATCTTTACTTATGCTGTGG, variant: 17:45822565 G / GC, variant: 17:45822322 C/T) were found in the heterozygous state in gnomAD (Karczewski et al., 2019). As their MAFs were <5×10^−6^, homozygosity rates for any of these three variants are well below the prevalence of MSMD (about 1/50,000). In the general population covered by the gnomAD database, seven variants have been identified in the homozygous state: A109G has a low CADD score, below the MSC, whereas H33Q, I339V, Y395C, M331I, R218H, and A29P have CADD scores above the MSC (**Fig. 1K** and **Fig. S5A**). The mouse ortholog of H33Q (H32Q) has been shown to be functionally neutral (Tantisira et al., 2004). None of these seven alleles affected transactivation of the WT-TBRE promoter (**Fig. 1L** and **Fig. S5B**). Thus, all the *TBX21* variants present in the homozygous state in gnomAD are functionally neutral. In our in-house cohort of >6,000 exomes from patients with various infectious phenotypes, P is the only patient carrying a rare bi-allelic variant at the *TBX21* locus. We investigated whether any of the other 28 non-synonymous variants of *TBX21* in our in-house database could underlie infections in the heterozygous state, by testing each of them experimentally. None had any functional impact on transcriptional activity (**Fig. S5C - E**). Therefore, the data for P and his family, our in-house cohort, and the general population suggest that inherited T-bet deficiency, whether complete or partial, is exceedingly rare in the general population (< 5.8 × 10^−8^). These findings also suggest that homozygosity for the Mut LOF variant of *TBX21* is responsible for MSMD in P.

### Homozygosity for the *TBX21* mutation underlies complete T-bet deficiency

We investigated the production and function of endogenous Mut T-bet in T-saimiri virus-transformed T cells (HVS-T) and primary CD4^+^ T cells from P. Levels of *TBX21* mRNA were normal, but endogenous T-bet protein levels were low in P’s cells (**Fig. 2A** and **B**, **Fig. S6A** and **B**). Together with the observation of low levels of Mut T-bet protein on overexpression (**Fig. S3**), these findings suggest that the *TBX21* mutation decreases T-bet protein levels by a post-transcriptional mechanism. T-bet transactivates *IFNG* and *TNF* by directly binding to their regulatory promoter or enhancer (Garrett et al., 2007; Kanhere et al., 2012; Soutto et al., 2002; Szabo et al., 2000; Tong et al., 2005a). Levels of spontaneous IFN-γ and TNF-α production by P’s HVS-T cells were much lower than those for HVS-T cells from healthy donors and heterozygous relatives (**Fig. 2 C - F**). This defect of cytokine production by *TBX21* mutant HVS-T cells was rescued by WT T-bet complementation (**Fig. 2G** and **Fig. S6C**). The functional impact of the T-bet mutation was also investigated in primary CD4^+^ T cells. Upon stimulation with phorbol 12-myristate 13-acetate (PMA) and ionomycin (P/I), IFN-γ production was almost entirely abolished in P’s expanded T_H_0 cell subset and TNF-α production was impaired; the production of both these molecules was rescued by WT T-bet (**Fig. 2H and I, Fig. S6D - F**). In T_H_1 conditions, exogenous IL-12 bypassed T-bet and induced moderate IFN-γ production by P’s cells, but the levels of this cytokine were still ~60-70% lower than those in healthy controls (**Fig. 2H and I**). We investigated other T-bet-dependent transcriptional targets, by performing RNA-seq to compare T_H_0 cells from controls, P, and P’s cells complemented with WT T-bet after incubation with anti-CD3/28 Ab beads. We found that the transcription of 455 was downregulated and that of 536 genes was upregulated in P’s cells (**Fig. S7A**, **Table S5**). The complementation of P’s cells with WT T-bet reversed the differential expression of 106 of the genes downregulated and 174 of the genes upregulated in P’s cells relative to controls (**Fig. S7B** and **C**). These targets were enriched in cytokine signaling pathway genes (**Fig. 2J**). We therefore decided to focus on genes involved in immunological signaling. Only 37 such genes were upregulated, and 33 downregulated in P’s cells, but these differences in expression relative to controls were reversed by WT T-bet (**Fig. 2K and Fig. S7D - F**). Known T-bet-dependent targets, such as *IFNG*, *CCL3* and *CXCR3*, were downregulated in this patient with T-bet deficiency, whereas *IFNGR2* expression was upregulated (**Fig. 2L**) (Iwata et al., 2017; Jenner et al., 2009). A set of new T-bet target genes was also identified, including *CCL1*, *CCL13*, *CCL4*, *CSF2*, *CXCR5*, *GZMM*, *IDO1*, *IL10*, *ITGA5*, and *ITGB5* (**Fig. 2L**). Collectively, our data indicate that the patient had AR complete T-bet deficiency, which affects the expression of a set of T-bet-dependent target genes.

**Figure 2.**
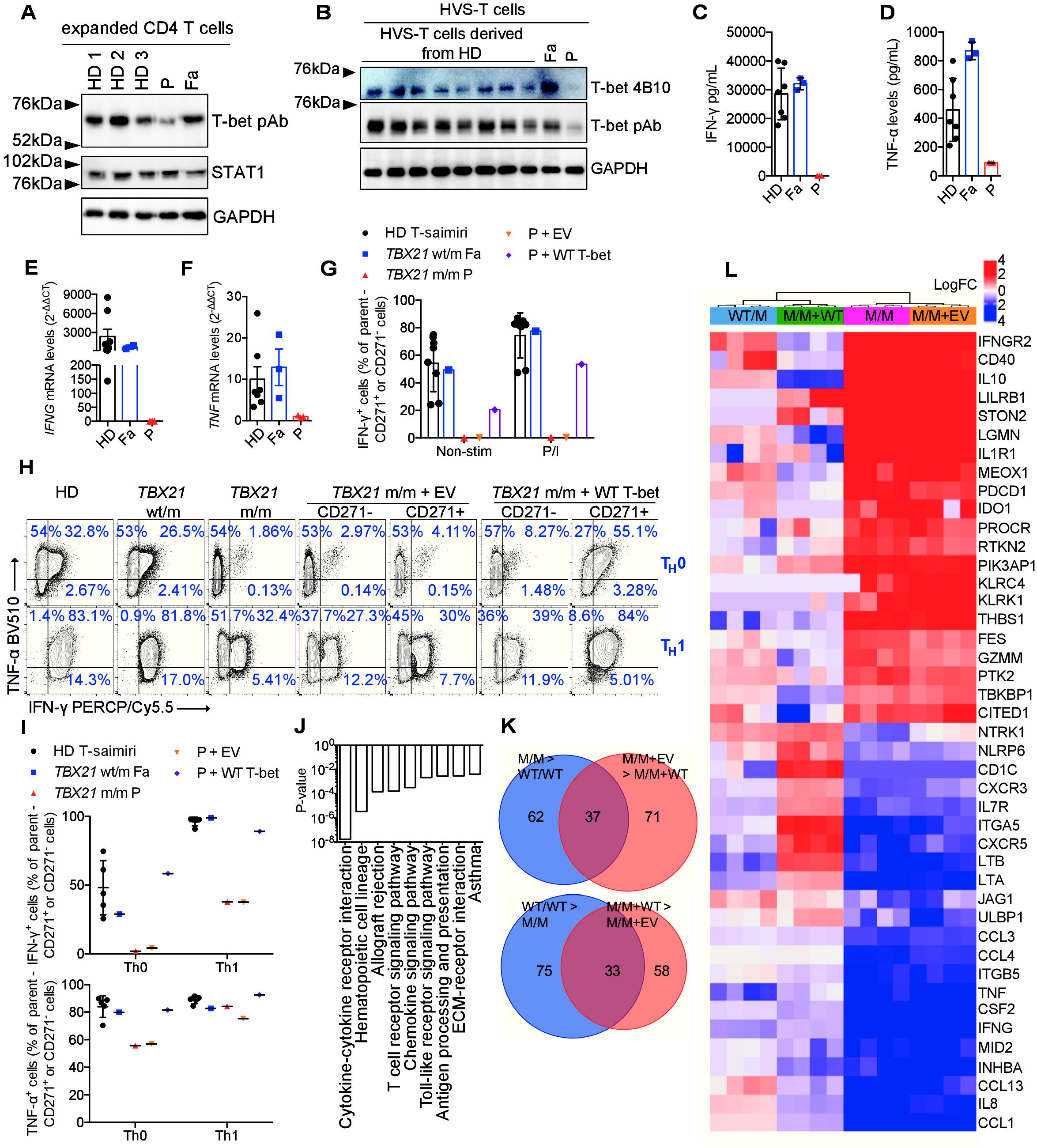

### T-bet induces permissive chromatin accessibility and CpG methylation in *IFNG*

We analyzed the molecular mechanisms by which T-bet controls transcription. Epigenetically, T-bet, is known to induce a permissive environment for transcription at the *IFNG* promoter through histone modifications and the suppression of CpG methylation (Lewis et al., 2007; Miller and Weinmann, 2010; Tong et al., 2005b). However, it remains unknown whether T-bet directly regulates chromatin accessibility in mice or humans. The regulation of CpG methylation at the genome-wide scale by T-bet has never been studied. We performed omni-ATAC-seq analysis and EPIC DNA CpG methylation array analysis with T_H_0 cells derived from P and controls (Corces et al., 2017). In P’s cells, a gain of chromatin accessibility was observed at 1,787 loci and a loss of chromatin accessibility was observed at 3,689 loci (**Fig. 3A** and **B, Fig. S8A - C**). We found that 666 and 1,649 of these loci, respectively, were subject to strict regulation by T-bet, as their gains and losses of chromatin accessibility were reversed by WT T-bet (**Fig. 3B, Table S6**). The chromatin regions opened out by T-bet were heavily occupied by bound T-bet, whereas those closed up by T-bet did not typically require T-bet binding (**Fig. 3C**) (Kanhere et al., 2012). An enrichment in T-box and ZBTB7B binding elements was observed in loci displaying an increased chromatin accessibility by T-bet, whereas an enrichment in Forkhead box elements was observed at loci at which chromatin accessibility was decreased by T-bet (**Fig. 3D - F**). The chromatin accessibility of 192 immunological genes, including *IL23R* and *IRF8*, two known MSMD-causing genes (Hambleton et al., 2011; Martínez-Barricarte et al., 2018; Salem et al., 2014), was increased by T-bet, whereas that of 75 immunological genes was decreased by T-bet (**Fig. 3G**, **Fig. S8** and **9**). Three known T-bet-dependent targets, *IFNG*, *TNF* and *CXCR3*, were among the top hits for the differentially regulated loci. The transcription start sites (TSS) of *IFNG* and *TNF*, the proximal promoter of *IFNG*, and the enhancers of *CXCR3* were inaccessible in T-bet deficient cells, and this inaccessibility was rescued by WT T-bet (**Fig. 3H - J**). The EPIC DNA CpG methylation array analysis identified 644,236 CpG sites that were differentially regulated (**Table S7**). Three CpG loci within *IFNG* were hypermethylated in conditions of T-bet deficiency, whereas their methylation was reduced to levels similar to those in controls on complementation with WT T-bet (**Fig. 3K** and **L**). *NR5A2*, *TIMD4*, *ATXN2*, *ZAK*, *SLAMF8*, *TBKBP1*, *CD247*, *HDAC4* and several other genes were also regulated in a similar manner (**Fig. 3K**). Interestingly, the methylation of six CpG loci within *ENTPD1* not previously linked to T-bet also increased in a T-bet-dependent manner (**Fig. 3K** and **L**). By contrast, *IL10* was a top target for which CpG methylation was drastically reduced in T-bet deficiency but rescued by WT T-bet (**Fig. 3K** and **M**). Taken together, these results demonstrate that T-bet orchestrates the expression of target genes by modulating both their chromatin accessibility and CpG methylation. Genome-wide omni-ATAC-seq and CpG methylation array analyses identified new epigenetic targets of T-bet (**Table S6** and **S7**). They also showed that chromatin accessibility at *IFNG* was increased by T-bet at both the TSS and promoter sites, whereas the CpG methylation of *IFNG* was dampened by T-bet at three different positions.

**Figure 3.**
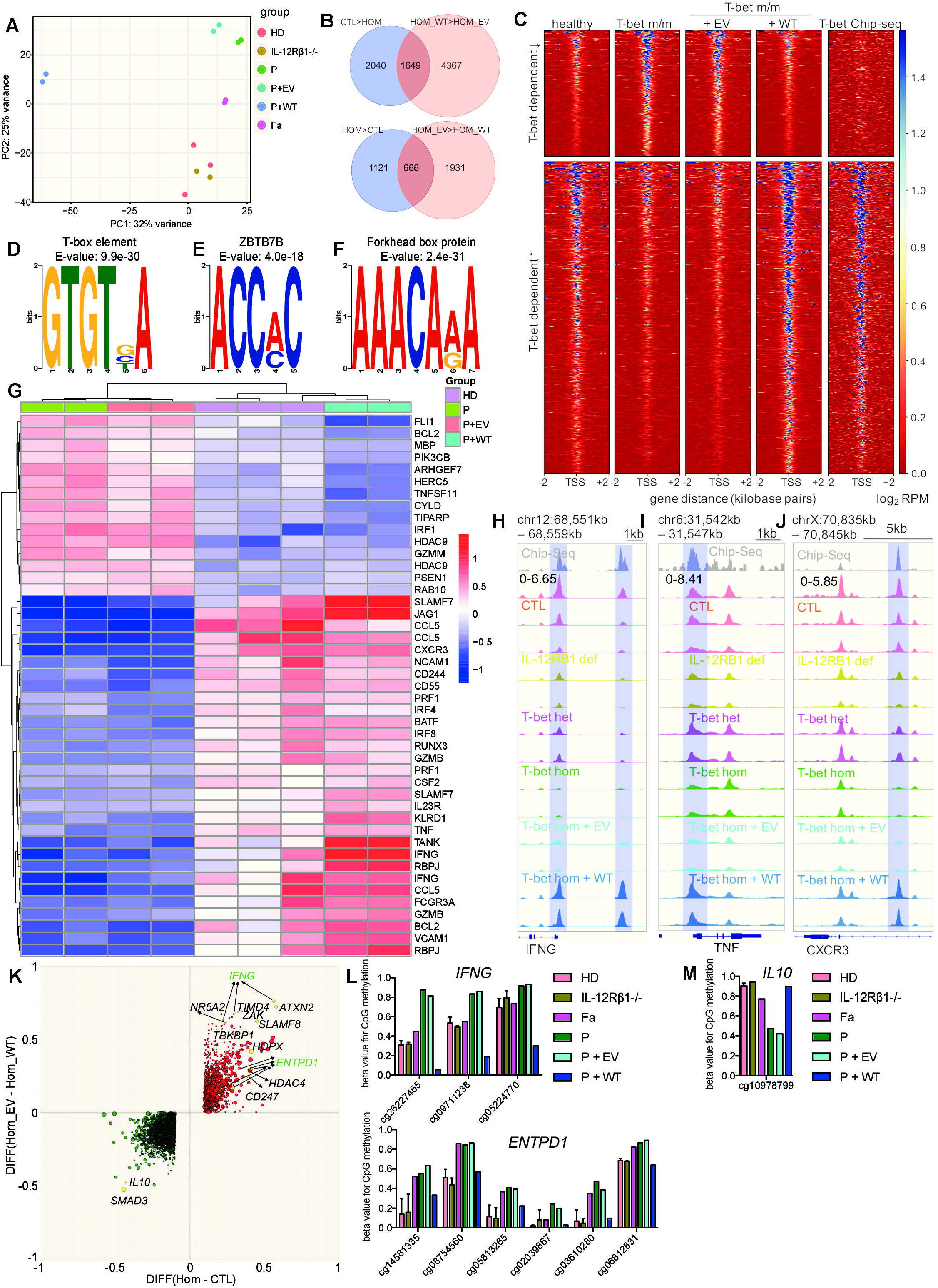

### T-bet deficiency impairs NK cell maturation

We then investigated the role of T-bet in the development of leukocyte lineages. Complete blood counts for fresh samples from P fresh samples showed that the numbers of lymphocytes, neutrophils, and monocytes were normal. We studied the PBMC subsets of P after the patient had been cured of mycobacterial disease, by mass cytometry (cytometry by time-of-flight, CyTOF) studies of 38 markers and comparisons with P’s parents, healthy donors, and patients with IL-12Rβ1 deficiency, the most common etiology of MSMD, as controls. The frequencies of plasmacytoid DCs (pDCs) and conventional DC 1 and 2 (cDC1 and cDC2) were not affected by human T-bet deficiency (**Fig. S10**). All major myeloid lineages were normal in P. We therefore focused on the development of lymphoid lineages. Total NK cells (defined as Lin^−^CD7^+^CD16^+^ or CD94^+^) were present in normal numbers in P. However, CD16^+^ and CD56^bright^ NK cells levels were ~25- and ~15-fold lower, respectively, in P than in the controls (**Fig. 4A – C**). Moreover, P had an abnormally high frequency of CD56^−^CD127^−^ NK cells **(Fig. S11)**; this NK cell subset has low levels of cytotoxicity and is rare in healthy and normal individuals (Björkström et al., 2010). The frequencies of ILC precursor (ILCP) and ILC2 in P were similar to those in healthy donors and IL-12Rβ1-deficient patients (**Fig. S12**). In stringent analyses, ILC1 and ILC3 are too rare for quantification in human peripheral blood (Lim et al., 2017). Overall, human T-bet is required for the correct development or maturation of NK cells, but not monocytes, DCs, ILC2 or ILCP.

**Figure 4.**
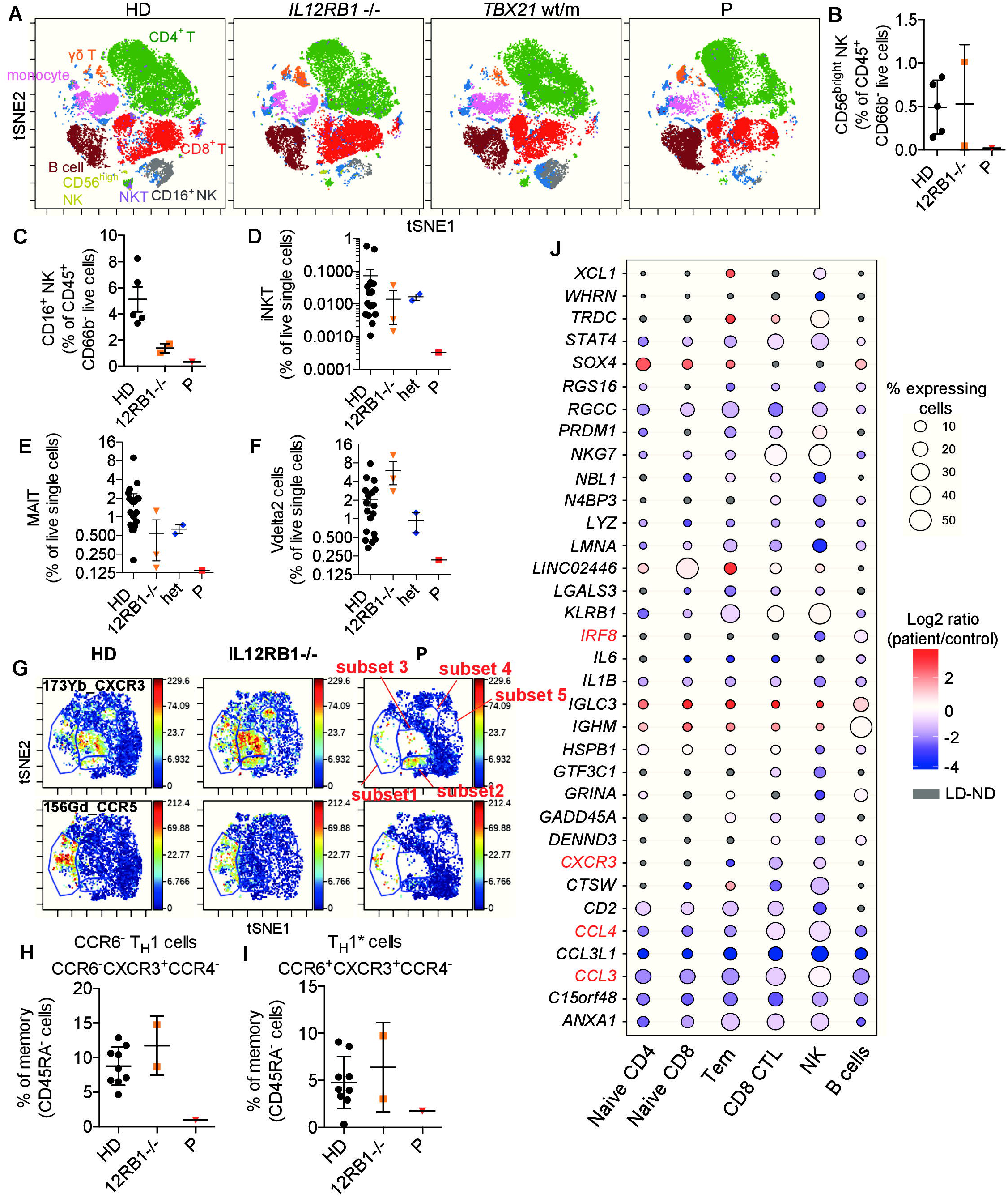

### Diminished iNKT, MAIT, and Vδ2^+^ γδ T-cell lineages in T-bet deficiency

The iNKT, MAIT, and γδ T cells are “innate-like” adaptive T-cell lineages with less T-cell receptor (TCR) diversity than conventional, “purely” adaptive αβ T cells (Chien et al., 2014; Crosby and Kronenberg, 2018; Godfrey et al., 2019). iNKT cells constitute a group of T cells with invariant TCRs combining properties from both T cells and NK cells (Crosby and Kronenberg, 2018). The iNKT cells of P were barely detectable (present at levels ~200-fold lower than in controls) **(Fig. 4D**, **Fig. S13A** and **B)**. MAIT cells express invariant Vα7.2-Jα33 TCRα restricted by a monomorphic class I-related MHC molecule, along with ligands derived from vitamin B synthesis (Kjer-Nielsen et al., 2012; Xiao and Cai, 2017). P also had a lower frequency of MAIT cells than controls (~15-fold) **(Fig. 4E** and **Fig. S13C**). Total γδ T-cell frequency in P was normal (**Fig. S13D**). However, the frequency of the Vδ2^+^ subset, a group of γδ T cells that recognize phosphoantigen (pAgs) (Gu et al., 2018; Harly et al., 2012; Vavassori et al., 2013), was low (~8-fold lower than control levels) in P, whereas the Vδ1^+^ subset of γδ T cells was normal (**Fig. 4F, Fig. S13E** and **F**). Mild abnormalities of B-lymphocyte development and antibody production unrelated to the patient’s mycobacterial disease were observed and will be reported in a separate study (Yang R, in preparation). CD4^+^ and CD8^+^ αβ T cells were the two most prevalent blood lineages of adaptive lymphocytes expressing a highly diverse αβ TCR repertoire. Antigen-driven CD8^+^ T-cell effector responses and the optimal induction of memory CD8^+^ T cells in mice are controlled by T-bet (Bettelli et al., 2004; Juedes et al., 2004; Sullivan et al., 2003). In the T-bet-deficient patient, total CD8^+^ T cells and the composition of naïve, central memory, effector memory and T_EMRA_ cells were normal (**Fig. S13G**). We further investigated CD8^+^ T cells in an unbiased manner, by automatic viSNE clustering with a panel of surface markers, including chemokine receptors (Amir el et al., 2013). We found no apparent difference between the memory CD8^+^ T cells of P and controls. However, a small subset of naïve CD8^+^ T cells (CD45RA^+^CD38^int^CXCR3^int^CCR6^−^CCR5^−^CD27^high^CD127^high^) was absent from P (**Fig. S13H**). Thus, the development of iNKT, MAIT cells, Vδ2^+^ γδ T cells, and a small subset of naïve CD8^+^ T cells is impaired in T-bet deficiency.

### Selective depletion of CCR6^−^ T_H_1 in CD4^+^ T cells in T-bet deficiency

Both P and his heterozygous parents had normal distributions of naïve and memory CD4^+^ T cells (**Fig. S14A**). We further analyzed individual CD4^+^ T-cell subsets, by viSNE clustering on antigen-experienced cells in particular (Amir el et al., 2013). Several memory CD4^+^ T-cell populations typically present in healthy donors were missing in P. Indeed, most of the CCR5^+^ cells (subset 1) and CXCR3^+^ cells (subsets 3 and 4), corresponding to T_H_1 cells in humans (Groom and Luster, 2011; Loetscher et al., 1998; Sallusto et al., 1998), were missing in P (**Fig. 4G** and **Fig. S14B**). A subset of CXCR3^high^ memory CD4^+^ T cells (CXCR3 subset 2 - CXCR3^high^CD27^low^CD127^low^CD38^int^) was, however, preserved (**Fig. 4G**). A cluster of CD127^+^CD27^+^CD25^+^CCR7^+^CD161^+^ cells (subset 5) was also missing (**Fig. 4G**). The frequency of classic CXCR3^+^CCR6^−^ T_H_1 cells was lower than that in controls (about nine-fold lower), whereas the frequency of CXCR3^+^CCR6^+^ non-classic T_H_1* cells, which are known to be mostly mycobacterium-specific, was unaffected (**Fig. 4H** and **I**). The frequencies of human T_H_2, T_H_17, and follicular helper (T_FH_) cells in peripheral blood were normal (**Fig. S14C - E**). However, the CXCR3^+^ T_FH_ cells, a group of T_H_1-biased T_FH_ cells that produce IFN-γ together with IL-21 in germinal centers (Velu et al., 2016; Zhang et al., 2019a), was diminished in P **(Fig. S14E)**. CXCR3^+^ regulatory T cells (Tregs), a group of T_H_1-skewed Tregs (Koch et al., 2009; Levine et al., 2017; Tan et al., 2016), and CCR5^+^ Tregs were also present at abnormally low levels, but the level of total Tregs was normal **(Fig. S14F** and **G)**. Thus, human T-bet deficiency selectively impairs the development of the classic CCR6^−^ T_H_1, CXCR3^+^ T_FH_, and CXCR3^+^ or CCR5^+^ Treg CD4^+^ T-cell subsets, but has no effect on the T_FH_, T_H_2, T_H_17, CCR6^+^ T_H_1*, and total Treg subsets, as shown by CyTOF and flow cytometry.

### Single-cell transcriptomic profile *in vivo* is altered by T-bet deficiency

We investigated the development and phenotype of leukocyte subsets in the patient further, by performing single-cell RNA-seq (scRNA-seq) with PBMCs from P and his father. The clustering of the various immune subsets yielded eight distinct major subsets: NK cells, pDCs, monocytes, B cells, CD8^+^ cytotoxic T lymphocytes (CTLs), CD8^+^ naïve, CD4^+^ naïve and CD4^+^ effector/memory T (T_EM_) cells (Becht et al., 2019). Consistent with the CyTOF results, normal frequencies of pDC, CTL, CD4^+^ naïve, T_EM_, and a low frequency of NK cells were obtained with scRNA-seq (**Fig. S15A**). We investigated the transcriptomic changes at single-cell level associated with T-bet deficiency, by filtering to select all genes with expression detected in > 5% of cells in at least one cluster, with at least a four-fold change in expression. We identified 34 genes as differentially regulated in T-bet-deficient cells relative to a heterozygous control. As for our RNA-seq data, some targets, including *CXCR3* in T_EM_, CD8^+^ CTL, and NK cells, *IRF8* in NK cells, and *CCL4* and *CCL3* in all cell types, before which expression was known to be dependent on T-bet, were downregulated in T-bet-deficient cells (**Fig. 4J** and **Fig. S15B**). The expression of *XCL1*, *STAT4*, *SOX4*, *LMNA* and *ANXA1* was also impaired in at least one subset of T-bet-deficient cells (**Fig. 4J** and **Fig. S15B**). In humans and mice, XCL1, STAT4, SOX4, LMNA and ANXA1 are known to be involved in T_H_1 immunity (Dorner et al., 2002, 2003, 2004; Gavins and Hickey, 2012; Kroczek and Henn, 2012; Nishikomori et al., 2002; Thieu et al., 2008; Toribio-Fernández et al., 2018; Yoshitomi et al., 2018). NKG7 is involved in the initiation of human T_H_1 commitment and its genetic locus is tightly occupied by T-bet (Jenner et al., 2009; Kanhere et al., 2012; Lund et al., 2005). The expression of *NKG7* in CD4^+^, CD8^+^ and B cells was dependent on functional T-bet, whereas *PRM1* was downregulated in CD4^+^ T and CD8^+^ CTL cells from P (**Fig. 4J**). The *IFNG* gene was weakly expressed across lymphocyte populations, as shown by scRNAseq, and its expression did not seem to be dependent on T-bet in basal conditions (data not shown). In addition, the expression of several genes not previously linked to T-bet was also altered in at least one cell subset in P (**Fig. 4J**). Thus, in addition to *IRF8*, *CXCR3*, *NKG7*, *CCL3* and *CCL4*, the expression of which was weak in at least one immune subset from this patient with human T-bet deficiency, consistent with the findings of RNA-seq and omni-ATAC-seq, T-bet is also important for the expression of a set of previously unknown target genes in immune subsets (**Fig. 4J**).

### Impaired IFN-γ production by NK, MAIT, Vδ2^+^ γδ T, and CD8^+^ T lymphocytes

Human IFN-γ is essential for antimycobacterial immunity, as all 30 known genetic etiologies of MSMD affect IFN-γ-dependent immunity. The *in vivo* development of NK, iNKT, MAIT, Vδ2^+^ γδ T, and classic T_H_1 cells was found to be impaired in P, but it remained possible that the IFN-γ production capacity of the remaining lymphocytes could compensate, thereby contributing to antimycobacterial immunity. We assessed the potential of P’s NK cells to respond to *ex vivo* stimulation with IL-12, IL-15 and IL-18. When stimulated, total NK cells from P displayed impaired degranulation, with low levels of CD107a expression, and almost no IFN-γ production (**Fig. 5A and B, Fig. S16A-C**). However, intracellular perforin and granzyme B levels in NK cells were unaffected by T-bet deficiency (**Fig. S16D** and **E**). The frequency of IFN-γ- producing total lymphocytes was also low in P (~18-fold lower than in controls), whereas the frequency of TNF-α-producing cells was only slightly lower than in the controls (~2.5-fold lower), in response to P/I stimulation *ex vivo* **(Fig. 5C – E)**. By contrast, no detectable IFN-γ-producing iNKT cells were detected in P or controls due to their very low frequency in peripheral blood. MAIT cells were present at a lower (~14-fold lower than controls) frequency *in vivo* in P, and these cells presented impaired production of IFN-γ (~9-fold lower than control levels) and TNF-α (~3 fold-fold lower) *ex vivo* **(Fig. 5F and G, Fig. S16F)**. Similarly, IFN-γ-producing Vδ2^+^ γδ T cells were barely detectable (~17-fold less frequent than in controls) whereas the frequency of TNF-α-producing Vδ2^+^ γδ T cells was only slightly low (~3-fold lower than control levels) *ex vivo* **(Fig. 5H and I, Fig. S16G)**. By contrast, we observed no difference in IFN-γ and TNF-α production by the cells of the Vδ2^−^ γδ T subset between P and controls **(Fig. 5J and K, Fig. S16H)**. CD8^+^ T cells secrete substantial amounts of IFN-γ upon microbial challenge (Wilson and Schoenborn, 2007). The frequency of CD8^+^ T cells producing IFN-γ in response to P/I was low in P (about 28-fold lower than in controls), whereas the frequency of TNF-α-producing CD8^+^ T cells was unaffected (**Fig. 5L and M, Fig. S16I**). Among the remaining circulating lymphocytes in the patient, NK, MAIT, Vδ2^+^ γδ T cells, and CD8^+^ T were equally defective for the production of IFN-γ *ex vivo* in response to IL-12, IL-15, IL-18 or P/I stimulation whereas Vδ2^−^ γδ T cells were not.

**Figure 5.**
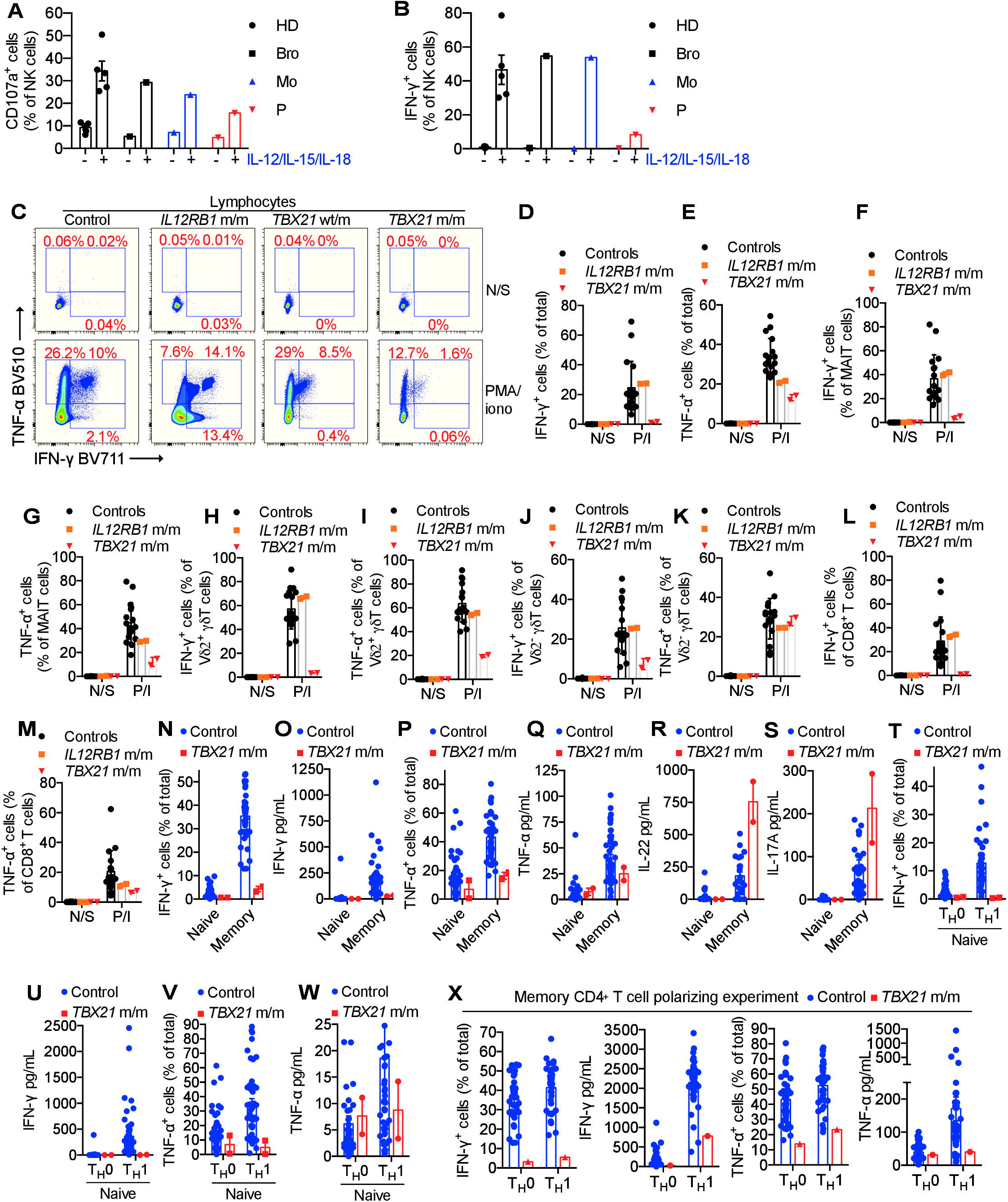

### Selective impairment of IFN-γ production by T_H_ cells in T-bet deficiency

T-bet was first discovered and has been most extensively studied in CD4^+^ T cells in the context of mouse T_H_1 cells (Szabo et al., 2000, 2002). This discover, together with that of GATA3 (Zheng and Flavell, 1997), revealed the molecular determinism of T_H_1/T_H_2 CD4^+^ T cell differentiation and paved the way for an understanding of T_H_17, iTreg, T_H_22, T_FH,_ and T_H_9 cell lineage determination (Zhu et al., 2010). We therefore investigated the impact of T-bet deficiency in primary CD4^+^ T cells. IFN-γ production by memory CD4^+^ T cells was impaired by T-bet deficiency (**Fig. 5N and O**). Another T_H_1 cytokine, TNF-α, was also produced in smaller amounts by memory CD4^+^ T cells from P than by those of most of the healthy controls (**Fig. 5P and Q)**. The memory CD4^+^ T cells of P produced larger amounts of the T_H_17 effector cytokines IL-22 and IL-17A than those from healthy donors (**Fig. 5R and S, Fig. S17A**). Unlike previous studies (Gokmen et al., 2013; Zhang et al., 2019b), we found that T-bet deficiency had no effect on IL-9 production *ex vivo* (**Fig. S17B and C**). Surprisingly, *ex vivo* T_H_2 cytokines from memory CD4^+^ T cells were not affected by human T-bet deficiency (**Fig. S17E - G**). P’s memory CD4^+^ cells produced less IL-21 *ex vivo* than the memory CD4^+^ cells of most of the controls (**Fig. S17H**). We then investigated the role of T-bet in human T_H_ cell differentiation *in vitro*. Naïve CD4^+^ T cells from P or healthy donors were allowed to differentiated in T_H_0, T_H_1, T_H_2, T_H_9, or T_H_17 conditions. The induction of IFN-γ production in naïve CD4^+^ T cells was abolished by T-bet deficiency under T_H_1 conditions (**Fig. 5T** and **U**). Similarly, T-bet-deficient naïve CD4^+^ T cells produced less TNF- α than the corresponding cells from most controls (**Fig. 5V** and **W**). Furthermore, the induction of IL-9 in various conditions *in vitro* was weaker in naïve CD4^+^ T cells from P than in the corresponding cells from most controls (**Fig. S17I** and **J**). *In vitro*-induced T_H_2 cells from P produced more IL-10, but not IL-13, than control cells (**Fig. S17K** and **L**). Even memory CD4^+^ T cells from P displayed impaired IFN-γ and TNF-α production under T_H_1 polarizing conditions (**Fig. 5X**). Thus, AR T-bet deficiency leads not only to defective IFN-γ and TNF-α production *ex vivo* and *in vitro*, but also to a moderate upregulation of the production of IL-17A and IL-22, two cytokines characteristic of T_H_17 cells (Lazarevic et al., 2011).

### Poor cellular response to BCG infection *in vitro* in T-bet deficiency

We investigated the molecular and cellular basis of BCG disease in P, by identifying the leukocyte subsets producing the largest amounts of IFN-γ in a T-bet-dependent manner during acute BCG infection *in vitro*. The infection of PBMCs with BCG induced IFN-γ production, which was further increased by stimulation with exogenous IL-12 **(Fig. 6A)**. PBMCs from P had low levels of IFN-γ production but normal levels of IL-6 and TNF-α production, and high levels of IL-5 and IL-13 production in response to BCG infection **(Fig. 6A, Fig. S18)**. Almost all the IFN-γ- producing cells had high levels of T-bet expression (**Fig. S19)**. Thus, T-bet^+^ IFN-γ^+^ double-positive cells were the major antimycobacterial cells, with a function potentially dependent on T-bet. However, T-bet^+^ IFN-γ^+^ cells were low during acute infection in P **(Fig. 6B)**. Among the T-bet^+^ IFN-γ^+^ cells of healthy donors, CD56^+^ NK, Vα7.2^+^ MAIT, Vδ2^+^ γδ T, and CD4^+^ T cells were the dominant responders in the absence of additional cytokine, while Vδ2^−^ γδ T, iNKT and CD8^+^ T cells represented the minority **(Fig. 6C and D, Fig. S20A)**. However, these subsets of T-bet^+^ IFN-γ^+^ cells were almost entirely depleted from P’s PBMCs following BCG infection **(Fig. 6D and Fig. S20B)**. We then investigated each leukocyte subset separately. Fewer than 1% of Vδ2^−^ γδ T cells, B cells, or CD4^+^ T cells became T-bet^+^ IFN-γ^+^ during BCG infection **(Fig. 6E)**. However, ~ 4-7% of NK cells, iNKT, MAIT cells, and up to ~ 15% of Vδ2^+^ γδ T cells from healthy donors, but not those from P, became T-bet^+^ IFN-γ^+^ in response to BCG infection, and the frequency of these cells was further increased by exogenous IL-12 **(Fig. 6E)**. Thus, the IFN-γ production controlled by T-bet during acute BCG infection *in vitro* takes place mostly in NK, MAIT, Vδ2^+^ γδ T and CD4^+^ T cells, but not in CD8^+^ T cells. These experimental findings *in vitro* do not exclude a contribution of other subsets *in vivo*. Thus, the NK, iNKT, MAIT, and Vδ2^+^ γδ T cells from healthy donors responded robustly to acute BCG infection *in vitro*, but these subsets were absent or functionally deficient in the patient with human T-bet deficiency.

**Figure 6.**
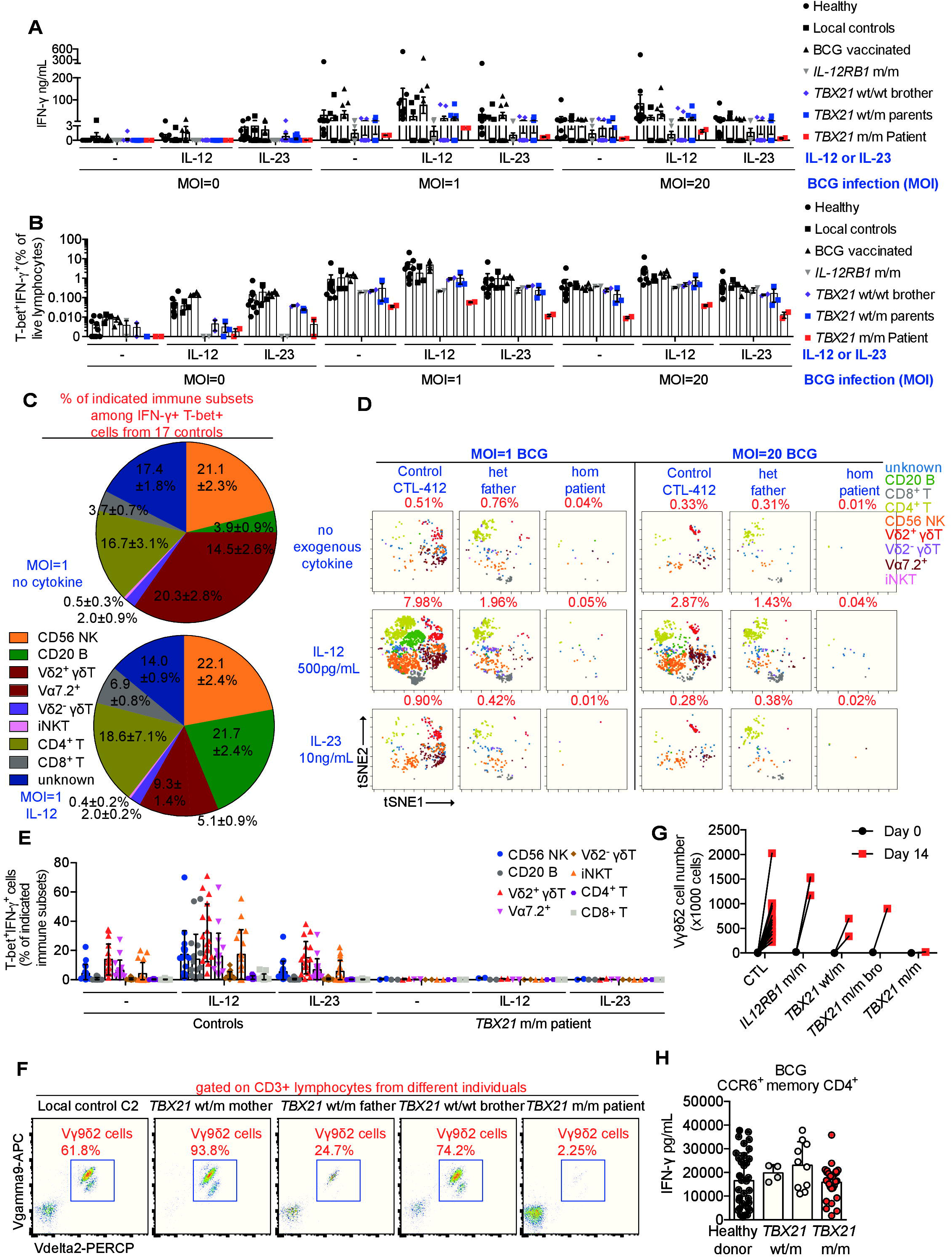

### Defective prolonged anti-BCG immunity mediated by Vδ2^+^ γδ T cells

The stimulation of PBMC *in vitro* with live BCG involves both antigens specifically recognized by mycobacterium-specific cognate αβ and γδ T cells and many other stimuli. BCG infection *in vitro* mimics acute infection *in vivo*, but may not be robust enough for investigations of the antigen-specific adaptive immune response, particularly as concerns prolonged adaptive immunity. We therefore studied Vδ2^+^ γδ T cells and CD4^+^ αβ T cells, two adaptive immune lymphocyte subsets that produced significant amounts of IFN-γ during BCG infection *in vitro* and are known to function in an antigen-specific manner. In PBMCs from healthy donors, ~15% of the Vδ2^+^ γδ T cells became T-bet^+^ IFN-γ^+^ during BCG infection **(Fig. 6E)**. Vδ2^+^ γδ T cells, a major subset of γδ T cells recognizing phosphoantigen (pAg) derived from microbial sources (Gu et al., 2018; Harly et al., 2012; Vavassori et al., 2013), are also known to play an essential role in the recall response to mycobacterial re-infection in humans and non-human primates (Chen, 2005; Shen et al., 2002). They proliferate vigorously in response to mycobacterial infection *in vivo* and can expand robustly in response to pAg-rich-lysates of mycobacterial species *in vitro* (Hoft et al., 1998; Modlin et al., 1989; Panchamoorthy et al., 1991; Parker et al., 1990; Tsukaguchi et al., 1995). We investigated whether Vδ2^+^ γδ T cells were functionally affected in the patient with T-bet deficiency, as these cells represented a small, but important proportion (~ 0.2%) of P’s peripheral lymphocytes. The populations of Vδ2^+^ T cells from all controls and relatives of P expanded vigorously following prolonged stimulation with BCG lysates. By contrast, no such expansion was observed for T-bet-deficient Vδ2^+^ T cells **(Fig. 6F** and **G, Fig. S21A** and **B)**. After two weeks of expansion, the levels of IFN-γ production by T-bet-deficient cells were lower than those for healthy control cells **(Fig. S21C)**.

### Redundant role of T-bet in IFN-γ production by BCG-specific cognate T_H_1* cells

It remains unclear whether the prolonged adaptive immunity to mycobacteria elicited by memory CD4^+^ T cells is dependent on T-bet. We addressed this issue by screening antigen-reactive T cell libraries established from CD4^+^CCR6^−^ (containing classic T_H_1 cells) and CD4^+^CCR6^+^ (containing T_H_1* *Mycobacterium*-responsive cells) memory subsets (Geiger et al., 2009). Consistent with our *in vivo* findings, the T cells in the CD4^+^CCR6^−^ and CD4^+^CCR6^+^ libraries had low levels of CXCR3 or IFN-γ **(Fig. S21D – G)**. P’s CD4^+^CCR6^+^ T-cell library responded robustly to BCG, tetanus toxoid and *C. albicans*, and his CD4^+^CCR6^−^ T-cell library responded normally to influenza virus, cytomegalovirus (CMV), and EBV **(Fig. S21H – N)**. Despite intact proliferation, IFN-γ production from T-bet-deficient T cells responding to influenza virus, EBV, tetanus toxoid, and *C. albicans* was weak **(Fig. S21O – S).**However, P’s CCR6^+^ T cells, consisting almost entirely of *Mycobacterium*-specific memory T_H_1* cells, proliferated robustly in response to BCG peptides. Moreover, their IFN-γ production was normal (Acosta-Rodriguez et al., 2007; Becattini et al., 2015), and their levels of IL-10 production were slightly higher **(Fig. 6H and Fig. S21T)**. The normal levels of IFN-γ production could not be attributed to cells with a revertant genotype, as reported in other T-cell primary immunodeficiency diseases (PIDs), because the IFN-γ^+^ BCG-specific T-cell clones still carried the *TBX21* indel variant (data not shown) (Davis et al., 2008; Revy et al., 2019). Thus, the prolonged immunity to BCG infection mediated by Vδ2^+^ γδ T and memory CD4^+^ T cells was divergently controlled by T-bet, as T-bet was required for the generation of long-term immunity due to Vδ2^+^ γδ T cells, but redundant for IFN-γ production by BCG-specific cognate T_H_1* cells.

## Discussion

We report the identification and study of a patient with MSMD due to inherited, complete T-bet deficiency. Key observations made in T-bet-deficient mice were validated in this human patient with T-bet deficiency: 1) the development of T_H_1 cells and their production of effector cytokines, including IFN-γ in particular, requires T-bet (Szabo et al., 2000, 2002); 2) the development of NK and iNKT cells is dependent on T-bet (Townsend et al., 2004) (**Table S8**); 3) the regulation of T-bet-dependent targets, including *CXCR3*, *TNF* and *IFNG*, involves both direct transactivation and epigenetic modulation (Miller and Weinmann, 2010). Accordingly, T-bet-deficient mice are highly vulnerable to mycobacteria, including *Mycobacterium tuberculosis* and *Mycobacterium avium* (Matsuyama et al., 2014; Sullivan et al., 2005), like mice deficient for other genes that govern IFN-γ immunity (Casanova, 1999). By contrast, despite the requirement of T-bet for immunity against a broad spectrum of pathogens following experimental inoculation in mice, the only apparent infectious phenotype of the T-bet-deficient patient is MSMD (**Table S9**). Our study provides compelling evidence that inherited T-bet deficiency is a genetic etiology of MSMD due to the disruption of IFN-γ immunity (Casanova et al., 2014). This experiment of nature suggests that T-bet is required for protective immunity to intramacrophagic mycobacteria but largely redundant for immunity to most intracellular pathogens, including viruses in particular. This is at odds with findings in mice, but consistent with other genetic etiologies of MSMD, all of which are inborn errors of IFN-γ immunity (Boisson-Dupuis et al., 2018; Martínez-Barricarte et al., 2018; Rosain et al., 2019) (**Table S9**). Our observation further suggests that the functions of T-bet unrelated to IFN-γ are redundant in humans. The identification of additional T-bet-deficient patients is required to draw firm conclusions. Yet, it is striking that humans genetically deprived of key immunological molecules, other than T-bet or IFN-γ, often show a much greater redundancy than the corresponding mutant mice (Casanova and Abel, 2004, 2018).

Our study also reveals unexpected immunological abnormalities not documented in T-bet-deficient mice that also contribute to the development of MSMD: 1) T-bet was required for the optimal development of two innate-like adaptive lineages of immune cells, MAIT and Vδ2^+^ γδ T cells; 2) T-bet was also required for the production of IFN-γ by the few cells from these two subsets that were able to develop. Unexpectedly, IFN-γ production by cognate purely adaptive *Mycobacterium*-specific T_H_1* CD4^+^ T cells was unaffected by T-bet deficiency. Taken together, impaired IFN-γ production by NK and iNKT cells, as in mice, and by MAIT and Vδ2^+^ γδ T cells, as shown here, accounts for MSMD in this patient with T-bet deficiency, despite normal T_H_1* development and function. By contrast, inborn errors of immunity that disrupt IFN-γ production by selective depletion of NK, iNKT, CD4^+^, or CD8^+^ αβT cells do not underlie mycobacterial disease, because of the compensation provided by other subsets **(Table S2).**Conversely, the loss of all T-cell subsets in severe combined immunodeficiency does result in predisposition to mycobacterial disease. Interestingly, a different combination of deficits accounts for MSMD in patients with RORγT deficiency, who lack iNKT and MAIT cells and whose γδ T and T_H_1* cells do not produce IFN-γ, while their NK cells are unable to compensate (Okada et al., 2015). T-bet and RORγT deficiencies are characterized by iNKT, MAIT, and γδ T-cell deficiencies, whereas an NK deficit is observed only in T-bet deficiency and a deficit of T_H_1* cells is observed only in RORγT deficiency. We found that human T-bet was essential for both innate (NK cells) and innate-like (iNKT, MAIT, and Vδ2^+^ γδ T cells) adaptive immunity to mycobacteria, but surprisingly redundant for classical, purely adaptive immunity (T_H_1*) to mycobacteria.

## Acknowledgments

We thank the patients and their families; the members of the laboratory for helpful discussions; Tatiana Kochetkov for technical assistance; Benedetta Bigio for computing support; Cecilia Lindestam Arlehamn and Alessandro Sette for providing peptide pools, and Dominick Papandrea, Yelena Nemirovskaya, Mark Woollett, and Cécile Patissier for administrative assistance. We also thank the Flow Cytometry Resource Center for technical support and the Rockefeller University Hospital for patient-oriented support. These institutions are supported in part by the National Center for Advancing Translational Sciences of the National Institutes of Health (UL1TR001866 to Rockefeller University). We also thank Steven Elledge (Brigham and Women’s Hospital, Harvard Medical School, Boston, MA, USA) for kindly providing the VirScan phage library used here for antibody profiling. This work was funded by the National Institute of Allergy and Infectious Diseases (R37AI095983 to J.L.C and U19AI118626 to F.S.), the Sackler Center for Biomedicine and Nutrition at the Center for Clinical and Translational Science, the Shapiro-Silverberg Fund for the Advancement of Translational Research at the Center for Clinical and Translational Science of the Rockefeller University (to R.Y.), the Research Grant Program of the Immune Deficiency Foundation (to R.Y.), the Integrative Biology of Emerging Infectious Diseases Laboratory of Excellence (ANR-10-LABX-62-IBEID) and the French National Research Agency under the “Investments for the future” program (ANR-10-IAHU-01), ANR-GENMSMD (ANR-16-CE17-0005-01 to J.B.), ANR-LTh-MSMD-CMCD (ANR-18-CE93-0008-01 to A.P), *Fonds de Recherche en Santé Respiratoire* (SRC2017 to J.B.), the French Foundation for Medical Research (FRM) (EQU201903007798), the SCOR Corporate Foundation for Science, ECOS Nord (C19S01-63407 to J.B.), the Swiss National Science Foundation (310030L_182728 to F.S.), the French Foundation for Medical Research (FRM) (EQU201903007798), the SCOR Corporate Foundation for Science, the St. Giles Foundation, the Rockefeller University, Howard Hughes Medical Institute, *Institut National de la Santé et de la Recherche Médicale* (INSERM), the Helmut Horten Foundation, Sidra Medicine and Paris Descartes University. The Canadian Center for Computational Genomics (C3G) is a Genomics Technology Platform (GTP) supported by the Canadian Government through Genome Canada. J. R. was supported by an INSERM *Poste d’accueil*. T.K. was supported by the Qatar National Research Fund (PPM1-1220-150017). S.G.T. was supported by a Program grant (1113904) and Principal Research Fellowship (1042925) awarded by the National Health and Medical Research Council of Australia. C.S.M. is supported by an Early/Mid-Career Development Research Fellowship (to C.S.M.) from the Ministry of Health of the NSW Government. R.Y. was supported in part by the Stony Wold-Herbert Fund and the Immune Deficiency Foundation.

