## Supplemental Figures, Table S1-4, S8, S9, Supplementary Materials for "Human T-bet governs innate and innate-like adaptive IFN-γ immunity against mycobacteria"

**This PDF file includes:**

Experimental model and subject details (including Case report)
Method details
Figs. S1 to S21
Figure legends for supplemental figures (Figs. S1 to S21)
Table S1 to S4, S8 and S9 (Table S5 – 7 were included separately as excel files)
Figure legends for supplementary tables (Tables. S1 to S9)
Reference for supplementary materials

### Materials and Methods

#### Human subjects

The patient and the relatives studied here were living in and followed up in Morocco. The study was approved by and performed in accordance with the requirements of the institutional ethics committee of Necker Hospital for Sick Children, Paris, France, and the Rockefeller University, New York, USA. Informed consent was obtained for the patient, his relatives, and healthy control volunteers enrolled in the study. This study was also approved by the Sydney Local Health District RPAH Zone Human Research Ethics Committee and Research Governance Office, Royal Prince Alfred Hospital, Camperdown, New South Wales, Australia (protocol X16-0210/LNR/16/RPAH/257). Written informed consent was obtained from participants or their guardians. Informed consent from participants in Switzerland was approved by the local ethical committee (CE3428 authorized by Comitato Etico Cantonale, www.ti.ch/CE). Experiments using samples from human subjects were conducted in the United States, France, Australia and Switzerland, in accordance with local regulations and with the approval of the IRBs of corresponding institutions. Plasma samples from unrelated healthy subjects used as controls for antibody profiling by phage immunoprecipitation-sequencing (PhIP-Seq) were collected at Sidra Medicine in accordance with a study protocol approved by the Clinical Research Ethics Board of Sidra Medicine.

#### Cell lines

HEK293T and Phoenix A retroviral packaging cells were cultured in IMDM (Gibco) supplemented with 10% fetal bovine serum (FBS). B cells from the patient or controls were immortalized in-house with Epstein Barr virus (EBV-B cells) and cultured in RPMI 1640 (Gibco) supplemented with 10% FBS. NK-92 cells were cultured in RPMI 1640 (Gibco) supplemented with 10% FBS, 1 mM sodium pyruvate, 200 U/mL recombinant proleukin IL-2 (Prometheus). *Herpesvirus saimiri*-transformed T (HVS-T) cells were generated with either *H. saimiri* strain C488 for transformation or the TERT transformation system (Wang et al., 2016). HVS-T cells were cultured in Panserin/RPMI 1640 (ratio 1:1) supplemented with 20% FBS, L-glutamine, gentamycin and 20 U/mL human rIL-2 (Roche). Isolated $CD4^+$ T cells were cultured in X-vivo 15/gentamycin/L-glutamine (Lonza) supplemented with human AB Serum (GemCell). Peripheral blood mononuclear cells (PBMCs) were cultured in RPMI 1640 supplemented with 10% FBS for stimulation experiments. All cell lines used tested negative for mycoplasma. HEK293T, Phoenix A and NK-92 cells were purchased from the ATCC.

### Method details

#### Case report

The patient (P) was born in 2015 to Moroccan first-cousin parents. He was vaccinated with BCG at the age of three months. After vaccination, he developed fever and left axillary lymphadenopathy accompanied by a cutaneous eruption. He received amoxicillin for 10 days, resulting in clinical improvement. At the age of six months, he was hospitalized for persistent fever, weight loss, cutaneous erythema, ear drainage and persistent oral thrush. Axillary adenopathy was present, with fistulation associated with hepatosplenomegaly and multiple abdominal adenopathies. The patient was treated with four antimycobacterial drugs; he displayed a good clinical response to 18 months of treatment and has been off antibiotics for 15 months. He was also found to have mild cytomegalovirus (CMV) viremia at the time (CMV PCR 3.74 log/mL, threshold < 2.4 log/mL), and was treated with ganciclovir. Serological PCR tests for CMV were negative at the age of two years, but weakly positive at the age of three years (IgM anti-CMV weakly positive, CMV PCR 3.62 log/mL, threshold < 2.4 log/mL). However, the patient showed no characteristic features of CMV infection. At the age of 11 months, he was hospitalized for four days due to airway hyperresponsiveness requiring treatment with inhaled steroids and albuterol. The patient had a high blood eosinophil count, documented since the age of since months (1,460/$mm^3$). By the age of 17 months, he had persistent upper respiratory tract inflammation, with manifestations of persistent rhinorrhea, difficult respiration, wheezing and coughing that improved on treatment with oral steroid and salbutamol. The patient has since been treated with inhaled steroids for persistent upper respiratory tract inflammation. The clinical manifestations affecting the upper respiratory tract persisted at the age of three years. No trigger was identified for these symptoms and signs, which were not triggered by exercise, activity, feeding or parasitic infections. A serum IgE allergen screen was negative for more than 250 allergens tested. The patient requires inhaled steroid therapy (fluticasone spray) every morning and evening. He also frequently visits the emergency room, and is hospitalized about once per month for exacerbations requiring oral steroid (beta-methasone) and inhaled salbutamol. The patient has been vaccinated for *Tetanus*, *Diphtheria*, and *Haemophilus* B, for all of which positive serum IgG result have been obtained. He also tested positive for anti-HSV-1 IgG, anti-EBV IgG, anti-CMV IgG, anti-mumps IgG and anti-parainfluenza IgG in clinical serology examinations. No anatomical abnormalities of the airways were noted. The patient has a brother who is healthy, with no history of severe infectious disease or airway hyperresponsiveness. The genetic cause of the patient's airway hyperresponsiveness will be studied separately.

**Phage immunoprecipitation-sequencing (PhIP-Seq)**

Patient serum and control samples were analyzed in duplicate by phage immunoprecipitation-sequencing (Hernandez et al., 2018; Xu et al., 2015). We used 10% liquid IVIg from pooled human plasma (Privigen® CSL Behring AG), human IgG-depleted serum (Supplier No HPLASERGFA5ML, Molecular Innovations, Inc.), and plasma samples from two unrelated healthy adult subjects and one unrelated healthy three-year-old boy for comparison. PhIP-Seq was carried out as previously described (Xu et al., 2015), but with the following modifications. We determined levels of total IgG in the serum or plasma samples with the Human IgG total ELISA Ready-SET-Go kit (Thermo Fisher Scientific) and incubated diluted samples containing approximately 4 μg of total IgG at 4°C overnight with $2 \times 10^{10}$ plaque-forming units (PFUs) of a modified version of the original VirScan phage library (Xu et al., 2015). Specifically, the T7 phage library used here for peptide display contained the same viral peptides as the original

VirScan phage library plus additional peptides derived from protein sequences of various microbial B-cell antigens available from the IEDB (www.iedb.org). For the computational analysis and background correction, we also sequenced the phage library before (input library sample) and after immunoprecipitation with beads alone (mock IP). We performed a number ($n$ = 46) pf technical repeats. Single-end sequencing was performed with the NextSeq500 system (Illumina), to generate approximately two million reads per sample and ~20 million reads for the input library samples. As previously described (Xu et al., 2015), reads were mapped onto the original library sequences with Bowtie 2 and read counts were adjusted according to library size. A zero-inflated generalized Poisson model was used to estimate the $p$-values, to reflect enrichment for each of the peptides. We considered peptides to be significantly enriched only if the –log10 $p$-value was at least 2.3 in all replicates. Species-specific score values were computed for each serum or plasma sample, by counting the significantly enriched peptides for a given species with less than a continuous seven-residue subsequence, the estimated size of a linear epitope, in common. We corrected for unspecific binding of peptides to the capture matrix, by also calculating species-specific background score values by counting the peptides displaying enrichment to the 90th percentile of the mock IP samples. These peptides were used for background subtraction.

**Whole-exome sequencing (WES), genotyping and genome-wide linkage (GWL) analysis**

All four members of the family tested were subjected to genomic DNA extraction followed by WES. Exome capture was performed with the SureSelect Human All Exon 50 Mb kit (Agilent Technologies). Paired-end sequencing was performed on a HiSeq 2000 (Illumina), generating 100-base reads. Sequences were aligned with the GRCh39 reference build of the human genome, with the BWA (Li and Durbin, 2009). Reads were then processed and variants were called with Genome

Analysis Toolkit, SAMtools, and Picard (Li et al., 2009; McKenna et al., 2010). The GATK Unified Genotyper was used to detect substitution and InDel calls (Depristo et al., 2011; McKenna et al., 2010). We used an annotation software system developed in-house to annotate all variants. All four members of the family were genotyped with Genome-Wide Human SNP Array 6.0 data. Genotype calling was achieved with the Affymetrix Power Tools Software Package (https://www.affymetrix.com/support/developer/powertools/changelog/index.html). Parametric multipoint linkage analysis was performed with MERLIN 1.1.2 software (Abecasis et al., 2002), assuming autosomal recessive inheritance with complete penetrance and a damaging allele frequency of 1 x $10^{-4}$. Allele frequencies were estimated for 562,685 SNP markers, with family founders and HapMap CEU trios. Markers were clustered using an $r^2$ threshold (--rsq parameter) of 0.4. The genetic variant of interest was confirmed by PCR amplification of the region (forward primer: 5'- gtgaggactacgcgctacc-3', and reverse primer: 5'-cagaagcattgtcgagccag-3') followed by Sanger sequencing.

**cDNA sequencing**

We used cDNA sequencing to characterize the consequences of the patients' variant. In brief, total RNA was extracted from EBV-B cells or PBMCs from the patient and healthy donors (Qiagen). We synthesized cDNA from mRNA, using Superscript III (Thermo Fisher Scientific). A region spanning the mutation site was amplified by PCR with the forward (5'-tagaagccaggcgtcagagc-3') and reverse R4 (5'- ctcggcattctggtaggcag-3'), R5 (5'-gcaatgaactgggtttcttgg-3'), or R6 (5'- gactcaaagttctcccggaatcc-3') primers. PCR amplicons were sequenced with the BigDye Terminator Cycle Sequencing Kit (Thermo Fisher Scientific). Alternatively, cDNA generated from patient or control PBMCs was amplified by PCR with the forward primer 5’- tctacactctttccctacacgacgctcttccgatctgtgaggactacgcgctacc-3’ and reverse primer 5’- gtgactggagttcagacgtgtgctcttccgatctgcaatgaactgggtttcttgg-3’, to amplify a region spanning the mutation in a first round of PCR. A second round of PCR was then performed on the PCR products from the first PCR as a template. We used the forward primer 5’-aatgatacggcgaccaccgagatctacactctttccctacacgac -3’ and the reverse primers 5’-caagcagaagacggcatacgagatcgtattcggtgactggagttcagacgtgtg -3’, 5’-caagcagaagacggcatacgagatctcctagagtgactggagttcagacgtgtg -3’, and 5’-caagcagaagacggcatacgagattagttgcggtgactggagttcagacgtgtg -3’, to barcode two healthy control cDNAs and the patient’s cDNA. Adaptor and barcoded PCR amplicons were mixed and purified, and used for next-generation sequencing by Nano Format 350 paired-end MiSeq sequencing (illumina). We aligned MiSeq data to the reference genome with RNA-STAR aligner (Dobin et al., 2013). The aligned sequencing data were then viewed with Integrative Genome Browser (Robinson et al., 2011).

**Reverse transcription-quantitative PCR (RT-qPCR)**

Total RNA was extracted from cells with the RNeasy Kit (Qiagen) or the Quick-RNA kit (Zymo Research). Reverse transcription was performed with the SuperScript III enzyme (Thermo Fisher Scientific). Messenger RNAs were quantified with the cDNA as a template and the *TBX21* (Probe 1: Hs00894393 and Probe 2: Hs00894391), *IFNG* (Hs99999041), *TNF* (Hs01113624), *IL5* (Hs01548712) (Thermo Fisher), *GUSB* (Thermo Fisher) probes.

**T-bet/*TBX21* overexpression**

A wild-type (WT) *TBX21* plasmid with a pCMV6 backbone was purchased from Origene (RC207902). The genetic variants of *TBX21*studied here, and the variants described in gnomAD (http://gnomad.broadinstitute.org/) or in our in-house database were introduced into pCMV6-WT *TBX21* by site-directed mutagenesis PCR with the CloneAmp HiFi PCR Premix (Takara Bio). *TBX21* WT plasmids or plasmids containing variants were used to transfect HEK293T cells in the presence of Lipofectamine 2000 reagent (Thermo Fisher Scientific) to achieve overexpression. The WT or mutant (Mut) *TBX21* was overexpressed in *in vitro*-expanded T cells, HVS-T cells, or NK-92 cells, as previously described (Martínez-Barricarte et al., 2016). In brief, the *TBX21* plasmid was subcloned into pLZRSires-ΔNGFR (Addgene). We then used pLZRS containing the WT or Mut *TBX21* to transfect Phoenix A cells, and retroviruses were produced and concentrated with Retro-X (Takara Bio). Concentrated retrovirus preparations were used for retroviral transduction.

**Immunoblotting**

Proteins were solubilized in RIPA buffer containing 10 mM Tris-Cl (pH 8.0), 1 mM EDTA, 0.5 mM EGTA, 1% Triton X100, 0.1% sodium deoxycholate, 0.1% SDS and 150 mM NaCl supplemented with protease-inhibitor cocktail (Complete mini, Roche). Alternatively, nuclear proteins from HEK293T transfectants were purified with the NE-PER Nuclear Extraction kit in accordance with the manufacturer's procedures (Thermo Fisher Scientific). Proteins were quantified with the Bradford assay. Equal amounts of lysate (5 µg to 50 µg of) were mixed with SDS sample buffer and boiled for 5min at 95°C before being subjected to electrophoresis in 12% acrylamide SDS-PAGE gels. The following antibodies were used for immunoblotting: mouse anti-human T-bet 4B10 monoclonal Ab (BioxCell), rabbit anti-human T-bet polyclonal Ab

(Proteintech), anti-Flag-M2 HRP-conjugated Ab (Sigma-Aldrich), anti-α-tubulin HRP-conjugated Ab (Proteintech), anti-lamin A/C Ab (Santa Cruz), anti-GAPDH HRP-conjugated Ab (Santa Cruz), anti-STAT1 monoclonal Ab (BD Biosciences), sheep ECL anti-mouse IgG HRP-conjugated Ab (GE Healthcare), and ECL anti-rabbit IgG HRP-conjugated Ab (GE Healthcare).

**Electrophoretic mobility shift assays (EMSA)**

HEK293T cells were transfected with empty plasmid (EV), WT or mutant *TBX21* pCMV6 plasmids. After 2 days, nuclear extracts were obtained from the cells with the NE-PER Nuclear Extraction Kit (Thermo Fisher Scientific). EMSA was performed as previously described (Liu et al., 2011). In brief, $^{32}$P was used to label consensus T-box responsive element (TBRE) duplexes of DNA oligos (WT oligo: 5'-gataatttcacacctaggtgtgaaatt-3', Mut Oligo as control: 5'-gataatttcacgtctaggtacgaaatt-3' and 5'-gataatttcgtacctagacgtgaaatt-3'). Unlabeled WT TBRE duplex was used as non-radioactive competitive probe. We mixed 10 μg of nuclear extract from each condition or 80 μg (8x) of nuclear extract from Mut *TBX21* transfectants with or without unlabeled probe, 2 μg mouse IgG1 control (BioxCell) or 2 μg mouse anti-T-bet 4B10 monoclonal Ab (BioxCell) and incubated for 30 min. $^{32}$P-labeled WT or Mut TBRE duplex was then added and the mixture was incubated for 30 min on ice in the presence of poly-dIdC (Sigma-Aldrich). Samples were subjected to polyacrylamide gel electrophoresis. The gel was dried and placed against X-ray film for 2 days, after which the radioactive signal was read on a Typhoon platform.

**Luciferase reporter assays**

HEK293T cells were transfected with the indicated expression plasmids, firefly luciferase plasmids under the control of WT or Mut *TBRE* (Janesick et al., 2012), as previously described, or with human *IFNG* promoters in a pGL4.10 backbone (-565 - +85 of *IFNG*), and with a constitutively expressed *Renilla* luciferase plasmid for normalization (pRL-SV40). Cells were transfected in the presence of Lipofectamine 2000 (Thermo Fisher Scientific) for 3 days. Luciferase levels were measured with Dual-Glo reagent, according to the manufacturer's protocol (Promega). Firefly luciferase values were normalized against *Renilla* luciferase values, and fold-induction is reported relative to controls transfect with empty plasmid.

**Generation and analysis of T-bet-deficient NK-92 cells**

T-bet-deficient NK-92 cells were generated according to a protocol described elsewhere (Sanjana et al., 2014). We used oligonucleotides for gRNA9 and gRNA15 (gRNA9_Forward: 5'-caccgtccaacaatgtgacccaggt-3', gRNA_Reverse: 5'-aaacacctgggtcacattgttggac-3'; gRNA15_Forward: 5'-caccgccgcactcaccgtccctgct-3', gRNA15_reverse: 5'-aaacagcagggacggtgagtgcggc-3'). DNA duplex was annealed from the oligonucleotides above, and was inserted into pLenti-CRISPR-V2 plasmids (Addgene). We then used pLenti-CRISPR plasmids, VSV-G envelope, and psPAX2 plasmids to transfect HEK293T cells in the presence of Lipofectamine 2000. Supernatants containing lentiviruses were harvested and concentrated with Lenti-X concentrator (Takara Bio). We resuspended$10^6$ NK-92 cells with 20x concentrated lentiviruses for gRNA9, gRNA15 or empty plasmid, in the presence of proleukin IL-2 (200 U/mL). After incubation for 2 days, we added 3.75 μg/mL puromycin to the culture for the selection of stably transduced NK-92 cells. After a further two weeks, cells were harvested for the Surveyor Nuclease Assay (Integrated DNA Tech) and a three-primer PCR-based screening system

(Harayama and Riezman, 2017) to confirm successful DNA editing. Cells were plated at a density of 0.7 cells in 200µL per well in a U-bottomed 96-well plate. Two weeks later, single-cell clones were screened with a three-primer PCR-based system, with Phire Direct PCR Master Mix (Thermo Fisher Scientific). The primers used for this screening assay were gRNA9_Forward: 5'-agcccctacccctaattcct-3', gRNA9_Reverse: 5'-ggcatctattccctgggacc-4', gRNA9_Reverse_in: 5'-cttttgaagagcaggtcctacct-3' and gRNA15_Forward: 5'-GCCTGAATATGACCCCCGTC-3', gRNA15_Reverse: 5'-caccactaccaccactaaagc-3'; gRNA15_Forward_in: 5'-acagagatgatcatcaccaagca-3'. Single-cell clones with biallelic disruptions were selected for validation by RT-qPCR and western blotting. EV-transduced single-cell clones were randomly picked as controls. Cells were then transduced with retroviruses generated from pLZRS-ires-ΔCD271 plasmids containing EV, WT, Mut or K314R *TBX21,* as described above (Martínez-Barricarte et al., 2016). Stably transduced cells were isolated with anti-CD271 antibody-coated beads (Miltenyi Biotec). Cells were stimulated with 50 pg/mL IL-12 (R&D) and 10 ng/mL IL-18 (InvivoGen), or were left unstimulated, and all cells were incubated for one day. The IFN-γ content of the supernatants was determined by ELISA. Intracellular IFN-γ production was measured with flow cytometry with intracellular staining (ICS). In brief, cells were fixed and permeabilized (BioLegend), and were subjected to ICS staining with FcBlock (Miltenyi Biotec), anti-T-bet BV421 (BioLegend) and anti-IFN-γ PE (BioLegend) before their use for ICS flow cytometry analysis.

**Cytokine determinations in *TBX21*-transduced primary CD4$^+$ T cells**

PMBCs from three healthy donors were stained with FcBlock (Miltenyi Biotec), anti-CD4-APC-Cy7, anti-CD45RA-Alexa488, anti-CCR7-Alexa647 and Live-Dead exclusion Aqua. Naïve

CD4$^+$ T cells were isolated by FACS. Cells were stimulated with anti-CD3/CD28 Ab Dynabeads (Thermo Fisher Scientific) at a 1:1 cell:bead ratio under $T_H0$ or $T_H2$ (recombinant human IL-4 12.5 ng/mL, R&D) polarizing conditions, in the presence of 100 U/mL proleukin IL-2. Fresh medium (X-vivo 15, 5% human AB serum) and cytokines were added every 3 days. After 7 days, cells were restimulated with Dynabeads at a 1:2 ratio (bead:cell). One day after restimulation, cells were transduced with retroviruses generated from pLZRS-ires-ΔCD271 plasmids containing WT or Mut *TBX21,* as previously described (Martínez-Barricarte et al., 2016). Transduced cells were isolated with anti-CD271 antibody-coated beads (Miltenyi Biotec) 14 days after stimulation. Intracellular cytokine production was determined by ICS. In brief, cells were restimulated with 25 ng/mL phorbol 12-myristate 13-acetate (PMA) and 0.5 μM ionomycin for 6 h in the presence of GolgiPlug (BD Bioscences). Cells were stained with Zombie-NIR live-dead exclusion dye (BioLegend), FcBlock (Miltenyi Biotec), and FITC-conjugated anti-CD271 antibody (BioLegend). Cells were then fixed, permeabilized and subjected to ICS staining with anti-TNF-α-BV510 (BioLegend), anti-IFN-γ-PERCP/Cy5.5 (BioLegend), anti-IL-4-PE, anti-IL-13-APC antibodies, followed by flow cytometry analysis with an Aurora cytometer (Cytek).

**HVS-T cell analysis**

We plated 200,000 HVS-T cells from healthy donors or P in a 96-well U-bottomed plate at a density of $1x10^6$ cells/mL. The plate was incubated for 2 days, and the culture supernatants and cells were harvested for ELISA or RT-qPCR for *IFNG* and *TNF*. HVS-T cells from P were transduced with retroviruses generated from pLZRS-ires-ΔCD271 plasmids containing WT *TBX21* or EV, as previously described (Martínez-Barricarte et al., 2016). After incubation for 5 days, the HVS-T cells were stimulated with 25 ng/mL PMA and 500 nM ionomycin or were left untreated; they were then incubated in the presence of GolgiPlug (BD Biosciences) for 6-8 h. Cells were stained with Zombie-NIR live-dead dye(BioLegend), FcBlock (Miltenyi Biotec), anti-CD4-BV750 (BD Biosciences), anti-CD3-BV650 (BD Biosciences), anti-CD8a-Pacific Blue (BioLegend), and anti-CD271-FITC (BioLegend) antibodies, fixed and permeabilized for ICS staining with anti-T-bet-PE (BioLegend), anti-IFN-γ-BV711 (BioLegend), anti-TNF-α-BV510 (BioLegend), anti-IL-13-APC (BioLegend), anti-IL-10-PE-Dazzle594 (BioLegend) antibodies. ICS analysis was performed on an Aurora cytometer (Cytek).

**Functional analysis of expanded T-bet-deficient CD4$^+$ T cells *in vitro***

CD4$^+$ T cells were isolated from PBMCs from five healthy donors, IL-12Rβ1-deficient (m/m) patients, the T-bet wt/m heterozygous father, and the T-bet m/m patient (P), with anti-CD4 antibody-coated beads (Miltenyi Biotec). They were then expanded with anti-CD3/CD28 antibody-coated Dynabeads (Thermo Fisher Scientific) at a 1:1 (cell:bead) ratio under $T_H0$ or $T_H1$ (2.5 ng/mL IL-12 R&D and anti-human IL-4 neutralizing Ab InVivoMab, BioXcell) conditions, in the presence of 100 U/mL proleukin IL-2 and anti-human IL-10 neutralizing Ab (Thermo Fisher Scientific). Medium (X-vivo15 5% human AB serum) and cytokine cocktails were refreshed every 3 – 4 days. After 7 days, the culture consisted of >95% CD4$^+$ T cells. Cells were restimulated with anti-CD3/CD28 Ab-coated Dynabeads every 7-8 days. P's CD4$^+$ T cells were transduced with retroviruses obtained from pLZRS-ires-ΔCD271 plasmids with or without WT *TBX21* or EV, as previously described (Martínez-Barricarte et al., 2016). Six days after transduction, cells were restimulated with 25 ng/mL PMA and 500 nM ionomycin in the presence of GolgiPlug (BD Biosciences) for 6 h. Cells were stained with Zombie NIR Live-Dead exclusion reagents (BioLegend), anti-CD271-FITC antibody and were then fixed and permeabilized. They were stained with anti-TNF-α-BV510, anti-IL-13-APC, anti-IL-4-PE, anti-IL-17A-BV785, anti-IL-10-PE-Dazzle594, anti-IL-5-BV421 and anti-IFN-γ-PerCp-Cy5.5 antibodies and subjected to flow cytometry in an Aurora cytometer (Cytek). Three weeks after their initial expansion, transduced $CD271^+$ $CD4^+$ T cells from P were isolated with anti-CD271 antibody-coated beads (Miltenyi Biotec). The cells isolated, and expanded $CD4^+$ $T_H0$ cells from healthy donors or T-bet wt/m individuals were restimulated with anti-CD3/CD28 antibody-coated beads in the absence of cytokine for 16 h or 48 h. Cytokine levels were determined in 13-Legendplex assays (BioLegend) on culture supernatants from cultures restimulated for 48 h . Total RNA was extracted from cells restimulated for 16 h for RNA-seq analysis (Qiagen). In brief, 100 ng of total RNA was used to generate RNA-Seq libraries with the Illumina TruSeq stranded mRNA LT kit (Cat# RS-122-2101). Libraries prepared with unique barcodes were pooled at equal molar ratios. The pool was denatured and sequenced on an Illumina NextSeq 500 sequencer with high-output V2 reagents and NextSeq Control Software v1.4 to generate 75 bp single reads, according to the manufacturer's protocol (Illumina, Cat# 15048776).

RNA-seq FASTQ files were obtained, and inspected by fastqc for quality control of the raw data. The sequencing reads of each FASTQ file were then mapped to the GENCODE human reference genome GRCh37.p13 (Frankish et al., 2019) with STAR aligner v2.6 (Dobin et al., 2013), and the quality of each mapped alignment in the BAM file was evaluated by RSeQC (Wang et al., 2012). Reads were quantified to generate gene-level read counts from the read alignment, with featureCounts v1.6.0 (Liao et al., 2014), based on GENCODE GRCh37.p13 gene annotation. The gene-level read counts were then normalized and log2-transformed by DESeq2 (Love et al., 2014), to obtain the gene expression value for all genes and all samples. An analysis of the differential expression of all genes and immune genes was performed to compare gene expression in the four

TBET samples (TBET_HET, TBET_HOM, TBET_HOM_EV, and TBET_HOM_WT) with that in five control samples, with each sample having two technical replicates. The 1,984 immune-related genes were identified from the immune system process gene set in the Molecular Signature Database (MSigDB) (Liberzon et al., 2015). By applying the trimmed mean of M values (TMM), normalization and gene-wise generalized linear model regression were performed in edgeR (Robinson et al., 2010). The genes displaying significant differential expression were selected on the basis of a $p$-value $\leq 0.05$ and a FDR $\leq 0.05$. The differential gene expression data were plotted as a heatmap by ComplexHeatmap (Gu et al., 2016), and genes and samples were clustered on the basis of complete linkage and Euclidean distance of gene expression values.

**Omni-ATAC-seq on CD4$^+$ T cells and analysis**

CD4$^+$ T cells derived from healthy donors, IL-12Rβ1 m/m patients, T-bet wt/m individuals and the T-bet m/m patient (P) were expanded under $T_H0$ conditions, as described above. Nine days after the retroviral transduction of CD4$^+$ T cells from P, as described above, CD271$^+$ cells were isolated with anti-CD271 antibody-coated beads (Miltenyi Biotec). Omni-ATAC-seq library preparation was performed as previously described (Corces et al., 2017). In brief, 50,000 cells from three healthy donors, two IL-12Rβ1 m/m patients, technical duplicates for a T-bet wt/m individual, the T-bet m/m P and P cells complemented with EV or WT *TBX21* were harvested, washed with 50 µl cold PBS, and centrifuged to obtain a pellet. Pellets were lysed with 50 µl cold lysis buffer consisting of 48.5 µl resuspension buffer (10 mM Tris-HCl pH 7.5, 10 mM NaCl, 3 mM $MgCl_2$ in water), 0.5 µl 10% NP-40, 0.5 µl 10% Tween-20, and 0.5 µl 1% digitonin. Lysates were incubated on ice and then washed with 0.1% Tween-20 resuspension buffer. Lysates were centrifuged and the nuclei in the pellet were subjected to Tn5 transposition with 50 µL of a mixture of 25 µL 2 x TD buffer, 16.5 µL PBS, 0.5 µL 10% Tween-20, 0.5 µL 1% digitonin, 2.5 µL Tn5 transposase, 5 µL nuclease-free water at 37 °C in a thermomixer operating at 1,000 rpm for 30 min. DNA fragments were extracted with the MinElute PCR purification kit and eluted in 30 µl EB buffer (Qiagen). We used 10 µL of eluted DNA for the amplification of DNA fragments with the NEBNext PCR master mix (New England BioLab), over six cycles, with the Ad1_forward and 15 indexed Ad2_reverse primers, as previously described (Buenrostro et al., 2013, 2015). The partially amplified library was subjected to quantitative PCR analysis with the Ad1 and Ad2 primers, Sybr Green reagents and NEBNext PCR master mix for 25 cycles. Additional PCR cycles were performed, based on 1/3 the maximal fluorescence for each sample, as previously described (Buenrostro et al., 2013, 2015). A left-sided isolation of PCR products was performed with AMPure beads (Beckman Coulter). DNA was quantified with a Qubit fluorimeter. Equal amounts of each sample were pooled and subjected to paired-end sequencing on a NextSeq high-output sequencer generating 75 bp reads.

The ATAC-seq reads were aligned with the hg19 genome from the Bsgenome.Hsapiens.UCSC.hg19 Bioconductor package (version 1.4.0) with Rsubread's align method in paired-end mode, with fragments between 1 and 5,000 base pairs long considered correctly paired (Liao et al., 2014). Normalized, fragment signal bigWigs were created with the rtracklayer package. Peak calls were made with MACS2 software in BAMPE mode (Feng et al., 2012; Zhang et al., 2008) and peak summits were used in MEME-ChIP software the identification of known and novel motifs (Ma et al., 2014). Differential ATAC-seq signals were identified with the DESeq2 package (Love et al., 2014; Ross-Innes et al.). Fragment length distribution plots and nucleosome-free/mono-nucleosome meta-TSS plots were produced with the soGGi package (de

Santiago and Carroll, 2018). Heatmaps of ChIP-seq and ATAC-seq signals were created from the normalized bigWigs signal with DeepTools2 software (Ramírez et al., 2016).

T-bet ChIP-seq data (SRR332104) and its input control (SRR332102) were retrieved as unaligned sequences from the European Nucleotide Archive (Kanhere et al., 2012). Sequences were aligned with the hg19 genome from the Bsgenome.Hsapiens.UCSC.hg19 Bioconductor package (version 1.4.0) with Rsubread's align method, with predicted fragment lengths calculated with the ChIPQC package (Carroll et al., 2014; Liao et al., 2014; de Santiago and Carroll, 2018). Normalized, fragment-extended bigWigs signals were created with the rtracklayer package.

**EPIC DNA methylation arrays and quantification of DNA methylation**

$CD4^+$ T cells derived from healthy donors, a T-bet wt/m individual and the T-bet m/m P were expanded under $T_H0$ conditions, as described above. Seventeen days after the retroviral transduction of $CD4^+$ T cells from P as described above, $CD271^+$ cells were isolated with anti-CD271 antibody-coated beads (Miltenyi Biotec). DNA from non-transduced or $CD271^+$ isolated $CD4^+$ T cells was extracted with a DNA mini kit (Qiagen). DNA was subjected to bisulfite conversion with the EZ DNA Methylation Kit (ZymoResearch). We quantified the DNA in all samples by performing EPIC arrays (Illumina), following the manufacturer’s instructions. Technical replicates were included for several samples during EPIC processin, to ensure that the measurements were reliable. All microarrays were scanned with the Illumina HiScan system. Data files were processed and converted to beta-values, corresponding to the percentage CpG methylation for a given CpG site. We wrote scripts to filter all loci, retaining those with higher levels of methylation in TBET_HOM and TBET_HOM_EV than the maximum methylation level in controls and lower levels of methylation in TBET_HOM_WT than in controls (group 1) and those with lower levels of methylation in TBET_HOM and TBET_HOM_EV than the minimum level of methylation in controls and higher levels of methylation in TBET_HOM_WT than the minimum level in controls (group 2). We then measured and plotted the difference in CpG methylation between the patient and controls, by subtracting the mean methylation level for the five controls from that for TBET_HOM and comparing the value obtained with the level of restored CpG methylation by subtracting the level of methylation in TBET_HOM_WT from the methylation in TBET_HOM_EV. On the plot, red dots represent the loci with higher levels of methylation in the patient than in controls, and green dots represent loci with lower levels of methylation in the patient than in controls. Dot size is proportional to the fold difference in methylation in patients relative to the mean value for controls. The genes of our interest are highlighted on the scatter plot.

**Immunophenotyping of leukocytes**

Metal-conjugated mAbs were obtained from Fluidigm according to the manufacturer's instructions. The following antibodies were included in the staining panel: anti-CD45-89Y, anti-CD57-113In, anti-CD11c-115In, anti-CD33-141Pr, anti-CD19-142Nd, anti-CD45RA-143Nd, anti-CD141-144Nd, anti-CD4-145Nd, anti-CD8-146Nd, anti-CD20-147Sm, anti-CD16-148Nd, anti-CD127-149Sm, anti-CD1c-150Nd, anti-CD123-151Eu, anti-CD66b-152Sm, anti-PD-1-153Eu, anti-CD86-154Sm, anti-CD27-155Gd, anti-CCR5-156Gd, anti-CD117-158Gd, anti-CD24-159Tb, anti-CD14-160Gd, anti-CD56-161Dy, anti-gdTCR-162Dy, anti-DRTH2-163Dy, anti-CLEC12A-164Dy, anti-CCR6-165Ho, anti-CD25-166Er, anti-CCR7-167Er, anti-CD3-168Er, anti-CX3CR1-169Tm, anti-CD38-170Er, anti-CD161-171Yb, anti-CD209-172Yb, anti-CXCR3-173Yb, anti-HLA-DR-174Yb, anti-CCR4-176Yb, anti-CD11b-209Bi. PBMCs from five healthy donors, two IL-12Rβ1-deficient patients, P and P's parents were included in this experiment. In total, 20,000 $DNA^+CD45^+CD66b^-$ events from individual CyTOF data files containing information for all markers except for CD45 and CD66b were deconvoluted into clusters by viSNE (Amir el et al., 2013). We also deconvoluted 5,000 memory $CD4^+$ T cells ($DNA^+CD45^+CD66b^-CD123^-CD3^+CD20^-CD56^-\gamma\delta TCR^-CD4^+CD8^-CD45RA^-$) into viSNE plots based on 21 surface markers (113ln_CD57, 115ln_CD11c, 149sm_CD127, 150Nd_CD1c, 153Eu_PD-1, 154Sm_CD86, 155Gd_CD27, 156Gd_CCR5, 158Gd_CD117, 159Tb_CD24, 163Dy_CRTH2, 165Ho_CCR6, 166Er_CD25, 167Er_CCR7, 169Tm_CX3CR1, 170Er_CD38, 171Yb_CD161, 173Yb_CXCR3, 174Yb_HLA-DR, 176Yb_CCR4, 209Bi_CD11b). Analyses were performed by manually gating populations for NK cells, naïve and memory subsets of $CD4^+$ and $CD8^+$ T cells, innate lymphoid cells (ILCs), Tregs and dendritic cell subsets. For T helper cell analysis, we included an additional four healthy donors from a separate experiment (same staining panel) in the analysis. Manual gating was used for TH analysis.

Immunophenotyping for MAIT, invariant NKT cells, Vδ2 and Vδ1 cells was performed with a flow cytometry staining panel containing FcBlock (Miltenyi Biotec), Zombie-NIR live-dead exclusion dye (BioLegend), anti-CD3-Alexa532 (Thermo Fisher Scientific), anti-γδTCR-FITC (Thermo Fisher Scientific), anti-Vδ2-APC-Fire750 (BioLegend), anti-CD56-BV605 (BioLegend), anti-CD4-BV750 (BD Biosciences), anti-CD8a-BV510 (BioLegend), anti-Vα7.2-BV711 (BioLegend), anti-Vα24-Jα18-PE-Cy7 (BioLegend), anti-Vδ1-Vioblue (Miltenyi Biotec), anti-CD161-PE (BioLegend) and anti-Vβ11-APC (Miltenyi Biotec) antibodies. Cells were analyzed with an Aurora cytometer (Cytek). Immunophenotyping for innate lymphoid cells and NK cells was confirmed separately with a conventional flow cytometry. In brief, biotinylated anti-human anti-CD1a (HI149), anti-CD3 (OKT3), anti-CD5 (UCHT2), anti-CD14 (61D3), anti-CD19

(HIB19), anti-CD34 (4H11), anti-CD123 (6H6), anti-CD203c (FR3-16A11), anti-CD303 (AC144), anti-TCRαβ (IP26), anti-TCRγδ (B1) and anti-FcεRIα (AER-37) antibodies were used in combination with streptavidin BUV661; conjugated anti-CD62L-FITC (Dreg-56), anti-CD94-PerCP-Vio700 (REA113), anti-TCRVα24Jα18-PE (6B11), anti-CD26-PE-CF594 (M-A261), anti-CD127-PE-Cy7 (eBioRDR5), anti-CRTH2-Alexa Fluor 647 (BM16), anti-CD161-Alexa Fluor 700 (HP-3G10), anti-EOMES-APC-eFluor 780 (WD1928), anti-NKp46-BV421 (9E2/NKp46), anti-CD45RA-BV570 (HI100), anti-CD117-BV605 (104D2), anti-RORγt-BV650 (Q21-559), anti-CD7-BV711(M-T701), anti-CD16-BUV496 (3G8), anti-CD25-BUV563 (2A3), anti-CD56-BUV737 (NCAM16.2) and anti-CD45-BUV805 (HI30); anti- T-bet-BV786 (O4-46), anti-GATA3-BUV395 (L50-823) antibodies were purchased from BioLegend, Thermo Fisher Scientific, BD Biosciences, or Miltenyi Biotec. Fc receptors were blocked with IgG from human serum (Sigma-Aldrich). Surface membrane staining was performed in Brilliant Stain Buffer (BD Biosciences). Transcription factors were stained with the Foxp3 staining buffer set (Thermo Fisher Scientific) according to the manufacturer's instructions. The fixable viability dye eFluor 506 (Thermo Fisher Scientific) was used to exclude dead cells. Samples were acquired on a Symphony A5 cytometer (BD Biosciences) with FACSDiva 8 software and were analyzed with FlowJo 10 (BD Biosciences).

Immunophenotyping was confirmed separately for the T- and B-cell subsets by conventional flow cytometry. In brief, PBMCs from healthy controls and P were labeled with mAbs against CD3, CD4, CD8, CD45RA, CCR7, CD127, CD25, CXCR5, CXCR3, CCR6, PD-1, and CD56. The $CD4^+$ and $CD8^+$ T-cell subsets were separated into naïve ($T_{Naive}$: $CCR7^+CD45RA^+$), central memory ($T_{CM}$: $CCR7^+CD45RA^-$), effector memory ($T_{EM}$: $CCR7^-CD45RA^-$), and terminally differentiated effector memory cells ($T_{EMRA}$: $CCR7^-CD45RA^+$) after initial gating on $CD8^-CD4^+$ or $CD8^+CD4^-$ cells. Proportions of regulatory T cells ($CD4^+CD127^{lo}CD25^{hi}$), total memory ($CD4^+CD45RA^-$), and circulating Tfh (cTfh; $CD4^+CD45RA^-CXCR5^+$) cells, and subsets of total memory and cTfh cells defined on the basis of CXCR3 and CCR6 expression (Th1: $CCR6^-CXCR3^+$, Th17: $CCR6^+CXCR3^-$, Th1*/Th1-17: $CCR6^+CXCR3^+$, Th2: $CCR6^-CXCR3^-$) were also determined.

**Single-cell (sc) RNA-seq and analysis**

We performed scRNA-seq on PBMCs obtained from the *TBX21* wt/m heterozygous father and the *TBX21* m/m patient. PBMCs were quickly thawed at 37°C and gently resuspended by serial additions of DMEM + 10% heat-inactivated (HI) FBS, according to the recommendations from the 10x Genomics "Sample preparation Fresh Frozen Human PBMC" protocol Rev C. Cells were centrifuged at 400 x *g* for 5 min, washed with DMEM + 10% HI-FBS, and cells were then counted and viability was assessed with the LIVE/DEAD™ Viability kit (Thermo Fisher Scientific) according to the manufacturer's guidelines. Following centrifugation at 400 x *g* for 5 min, cells were resuspended at a concentration of 1000 cells/µl in PBS + 0.04% BSA and loaded onto a 10 x Genomics Chromium chip for single-cell capture. Reverse transcription and library preparation were performed with Chromium Single Cell 3′ Reagent Kits (v2) in accordance with the manufacturer's guidelines, and library quality was assessed with a Bioanalyzer DNA chip. Each library was sequenced on two lanes of an Illumina HiSeq 4000 sequencer in a 28 bp/98 bp paired-end configuration.

The sequence-read quality of individual sequencing lanes was assessed with BVAtools (https://bitbucket.org/mugqic/bvatools). Cell Ranger v3.0.1 was used to aggregate the sequences from two lanes/samples, to map reads to the hg38 human reference genome assembly, for filtering, and counting barcodes and UMIs; a list of UMI counts was thus obtained for each gene and each cell. Cells outside the [5%,95%] interval for these metrics or with >10% mitochondrial genes were excluded. The DoubletFinder package was used to identify and filter out cell doublets (McGinnis et al., 2019). After the removal of dead cells and doublets, 10,409 cells for the T-bet wt/m heterozygous father and 11,741 cells for the T-bet m/m P were analyzed together with the Seurat v3 R package (Stuart et al., 2019), and cell clustering was performed by the Uniform Manifold Approximation and Projection dimension reduction method (Becht et al., 2019), using the most variable genes, but excluding mitochondrial and ribosomal protein genes. This analysis identified eight distinct cell clusters; the marker genes for each cluster were identified with Seurat and used to determine cell types. Differential expression was analyzed by comparing UMI gene counts between the patient and the control for each cluster, with the MAST approach (Finak et al., 2015). Genes with a $p$-value < $10^{-25}$ for differential expression were analyzed in more detail. For each selected gene, cells in which the expression of the gene was detectable were counted for the index case and the control in each cluster, to reduce the dropout bias inherent to droplet scRNA approaches. The number of cells expressing the gene was normalized according to the number of cells in each cluster and the fold-difference in expression was calculated by comparing the index case and the control. For the figures, cluster annotations and the levels of expression of genes of interest were projected onto the UMAP clustering, but with the cells of the patient and the control shown separately.

**Stimulation of NK cells *ex vivo***

PBMCs from five healthy donors including a travel control, the T-bet wt/m mother, the T-bet wt/wt brother and the T-bet m/m P were left untreated or stimulated with IL-12 (Miltenyi

Biotec), IL-15 (Miltenyi Biotec) and IL-18 (MBL/CliniSciences) for 18 h. Cells were treated with GolgiPlug and GolgiStop (BD Biosciences), and stained with anti-CD107a-PE antibody (Thermo Fisher Scientific) 4 h before harvest. Cells were harvested and stained with eFluor506 live-dead reagents (Thermo Fisher) and then with surface Abs, including anti-CD1a-FITC (BioLegend), anti-CD3-FITC (Thermo Fisher Scientific), anti-CD5-FITC (Thermo Fisher Scientific), anti-CD14-FITC (Miltenyi Biotec), anti-CD19-FITC (BioLegend), anti-CD34-FITC (BioLegend), anti-CD123-FITC (BioLegend), anti-CD203c (BioLegend), anti-CD303-FITC (BioLegend), anti-FcεRIα-FITC (BioLegend), anti-TCRαβ-FITC (Thermo Fisher Scientific), anti-TCRγδ-FITC (BioLegend), anti-CD94-PerCP-Vio700 (Miltenyi Biotec), anti-CD127-PE-Cy7 (Thermo Fisher Scientific), anti-NKp46-APC (BD Biosciences), anti-CD7-BV711 (BD Biosciences), anti-CD56-BV786 (BD Biosciences), anti-CD16-BUV737 (BD Biosciences), and anti-CD45-BUV805 (BD Biosciences) antibodies. All the markers targeted by FITC-conjugated Abs were considered to be Lineage-specific markers (Lin). Cells were fixed and permeabilized (Thermo Fisher Scientific) and subjected to intracellular staining with anti-granzyme B-PE-CF594 (BD Biosciences), anti-EOMES-APC-eFluor780 (Thermo Fisher Scientific), anti-perforin-BV421 (BD Biosciences), anti-TNF-α-BV605 (BioLegend) and anti-IFN-γ-BUV395 (BD Biosciences) antibodies. ICS and surface staining were analyzed with an LSRFortessa flow cytometer (BD Biosciences).

**CD4$^+$ T-cell isolation and functional characterization *ex vivo* and T$_H$ differentiation *in vitro***

PBMCs from healthy donors and P were incubated with mAbs against CD4, CD45RA, CCR7, CXCR5, CD127 and CD25. Naïve and memory CD4$^+$ T cells were isolated by first excluding Tregs (CD25$^{hi}$CD127$^{lo}$) and then sorting CD4$^+$CD45RA$^+$CCR7$^+$CXCR5$^-$ and CD4$^+$CD45RA$^-$CCR7$^\pm$ cells, respectively. Isolated naïve and memory CD4$^+$ T cells were then cultured in 96-well round-bottomed well plates (30-40 x$10^3$ cells/well) with T-cell activation and expansion (TAE) beads (anti-CD2/CD3/CD28; Miltenyi Biotech) alone ($T_H0$) or under $T_H1$ (50 ng/mL IL-12), $T_H2$ (100 U/mL, IL-4), $T_H9$ (2.5 ng/mL TGF-β, 100 U/mL IL-4), or $T_H17$ (2.5 ng/mL TGF-β, 50 ng/mL IL-1-β, 50 ng/mL IL-6, 50 ng/mL IL-21, 50 ng/mL IL-23, 50 ng/mL PGE2) polarizing conditions. After 5 days, supernatants were harvested and the production of IL-4, IL-5, IL-9, IL-10, IL-13, IL-17A, IL-17F, IFN-γ and TNF-β was assessed with cytometric bead arrays (Becton Dickinson); IL-22 secretion was measured by ELISA (eBioscience). For cytokine expression, activated $CD4^+$ T cells were restimulated with PMA (100 ng/mL)/ionomycin (750 ng/mL) for 6 h, with brefeldin A (10 μg/mL) added after two hours. Cells were then fixed and the intracellular expression of IL-4, IL-9, IL-13, IL-10, IL-17A, IL-17F, IL-22, IL-21 and IFN-γ was determined.

**Stimulation of PBMCs with live *M. bovis*-BCG or PMA/ionomycin**

Ten healthy donors and four BCG-vaccinated healthy donors were recruited at the Rockefeller University (New York, USA). Three local controls, one IL-12Rβ1 m/m patient and the T-bet m/m P and his members of his family were recruited at Necker Hospital (Paris, France). PBMCs from these individuals were isolated and cryopreserved under the same conditions. We plated 200,000 cells per well in 96-well U-bottomed plates at a density of 1 x $10^6$ cells/ml. PBMCs from the T-bet m/m P, his T-bet wt/m mother, T-bet wt/wt brother, and the IL-12Rβ1-deficient patient were analyzed in technical duplicates. For each sample, cells were plated in the presence or absence of live *M. bovis*-BCG at a MOI=1 or 20, and in the presence and absence of recombinant IL-12 (500 pg/mL, R&D) or recombinant IL-23 (10 ng/mL, R&D). Golgiplug (BD Biosciences) was added to each well after 40 h of stimulation. Cells were also stimulated with 25 ng/mL PMA and 500 nM ionomycin. Eight hours later, supernatants were harvested for cytokine determinations in a 13-plex Legendplex assay (BioLegend), and cells were collected by centrifugation for FACS staining. Cells were stained with the Zombie NIR Viability kit (BioLegend) and were then surface-stained with Abs including FcBlock (Miltenyi Biotec), anti-CD3-Alexa532 (Thermo Fisher Scientific), anti-γδTCR-FITC (Thermo Fisher Scientific), anti-Vδ2 TCR-APCFire750 (BioLegend), anti-CD56-BV605 (BioLegend), anti-CD4-BV750 (BD Biosciences), anti-CD8a-Pacific Blue (BioLegend), anti-Vα7.2 TCR-APC (BioLegend), anti-Vα24-Jα18 (iNKT)-PE/Cy7 (BioLegend), and anti-CD20-BV785 (BioLegend) antibodies. Cells were then fixed and permeabilized for staining for intracellular proteins with antibodies including FcBlock (Miltenyi Biotec), anti-IFN-γ-BV711 (BioLegend), anti-TNF-α-BV510 (BioLegend), anti-T-bet-PE (BioLegend), and anti-IL-10-PE/Dazzle594 (BioLegend) antibodies. Samples were then analyzed on an Aurora cytometer (Cytek). Flow data were analyzed with Cytobank. T-bet$^{+}$ IFN-γ$^{+}$ double-positive cells were gated manually from the live single-lymphocyte population. These T-bet$^{+}$ IFN-γ$^{+}$ double-positive cells were clustered with the viSNE algorithm, with CD3-Alexa532, γδTCR-FITC, Vδ2 TCR-APCFire750, CD56-BV605, CD4-BV750, CD8a-Pacific Blue, Vα7.2 TCR-APC, Vα24-Jα18 (iNKT)-PE/Cy7, CD20-BV785, TNF-α-BV510 and IL-10-PE/Dazzle594 (Amir el et al., 2013). Automated viSNE clusters were plotted with tSNE1 and tSNE2. Clusters were then overlaid on manually gated immune cell subsets for illustration.

**Vδ2 γδT cell stimulation**

Lysates of *M. bovis*-BCG was prepared and provided by Dr. Carl F. Nathan. PBMCs from healthy donors, local control donors, BCG-vaccinated healthy donors, IL-12Rβ1 m/m patients, the T-bet m/m P and his relatives were resuspended at a density of $1x10^{6}$ cells/mL. We plated 200 μL cells per well for each condition, in a 96-well U-bottomed plate. We added BCG lysate to each well at a final concentration of 5 μg/mL. Recombinant IL-2 (Roche) was added to a final concentration of 10 U/mL. Fresh medium and IL-2 were added to the culture every three to four days. Eleven days after stimulation, the cells were fully confluent in the 96-well plate. Cells were then transferred to a 48-well plate and each well was topped up to500 μL per well with medium. After 14 days of stimulation, the cells were stained with FcBlock (Miltenyi Biotec), Zombie-NIR live/dead (BioLegend), anti-CD3-Alexa532 (Thermo Fisher Scientific), anti-γδTCR-FITC (Thermo Fisher Scientific), anti-Vδ2 TCR-APCFire750 (BioLegend), anti-CD56-BV605 (BioLegend), anti-CD4-BV750 (BD Biosciences), anti-CD8a-BV510 (BioLegend), anti-β11TCR-APC (Miltenyi Biotec), anti-CD161-PE (BioLegend), anti- Vα7.2 TCR-BV711 (BioLegend), anti-Vα24-Jα18 (iNKT)-PE/Cy7 (BioLegend), anti-Vδ1-Vioblue (Miltenyi Biotec), anti-CD16-BV650 (BioLegend), anti-CD20-BV785 (BioLegend), anti-αβTCR-Alexa700 (BioLegend) antibodies and analyzed by flow cytometry on an Aurora cytometer (Cytek).

**Isolation and screening of CD4$^+$ T cell lines**

The T-cell clone experiment was performed as previously described (Geiger et al., 2009). In brief, CCR6$^+$ and CCR6$^-$ CD4$^+$ memory T cells were sorted from CD14$^-$ PBMCs with a FACS Aria (BD Biosciences), excluding CD45RA$^+$CCR7$^+$ naïve cells, and CD25$^+$, CD19$^+$, and CD8$^+$ cells. Sorted T cells (500 cells/well) were polyclonally stimulated with 1 μg/mL PHA (Remel) in the presence of irradiated (45 Gy) allogeneic feeder cells ($10^4$ per well) and IL-2 (500 IU/mL) in RPMI complete medium. After expansion for at least 20 days, the T-cell lines were analyzed by surface or intracellular staining with the following monoclonal antibodies: anti-CCR6-PE (BD Biosciences; clone: 11A9); anti-CXCR3-APC (BD Biosciences; clone: 1C6); anti-IFN-γ-FITC or anti-IFN-γ-PE (BD Biosciences, clone: B27); anti-IL-17A-eFluor660 FITC (eBiosciences, clone: eBio64DEC17); anti-IL-4-PE (BD Biosciences, clone: MP4-25D2); anti-IL-22-PerCP-eFluor-710 (Thermo Fisher Scientific, clone: 22URTI). For identification of the antigen-responding T-cell clones, the T-cell lines were also screened by culturing thoroughly washed cells ($2.5\times10^5$ cells/well) with autologous irradiated B cells ($2.5\times10^4$ cells/well), with or without a three-hour pulse of the following antigens: *M. tuberculosis* peptide pool (0.5 µg/mL/peptide, comprising 207 15-mer peptides), BCG peptide pool (0.5 µg/mL/peptide, comprising 211 15-mer peptides), HCMV peptide pool (0.5 µg/mL/peptide, comprising 76 15-20-mer peptides), EBV peptide pool (0.5 µg/mL/peptide, comprising 46 15-20-mer peptides), Influenza virus HA peptide pool (2 µg/mL/peptide, comprising 351 15-mer peptides), *Candida albicans* peptide pool (0.5 µg mL/peptide, comprising 252 15-mer peptides) or tetanus toxoid peptide pool (1 µg/mL/peptide, comprising 125 15-20-mer peptides). These peptide pools were kindly provided by Cecilia Lindestam Arlehamn and Alessandro Sette at the La Jolla Institute for Immunology (Lindestam Arlehamn et al., 2016). Proliferation was assessed on day 4, after incubation for 16 h with 1 µCi/mL [$^3$H]-thymidine (GE Healthcare). The concentrations of cytokines in the culture supernatants produced by the antigen-responding T-cell clones were measured after 48 h of stimulation with cytometric bead arrays (eBioscience).

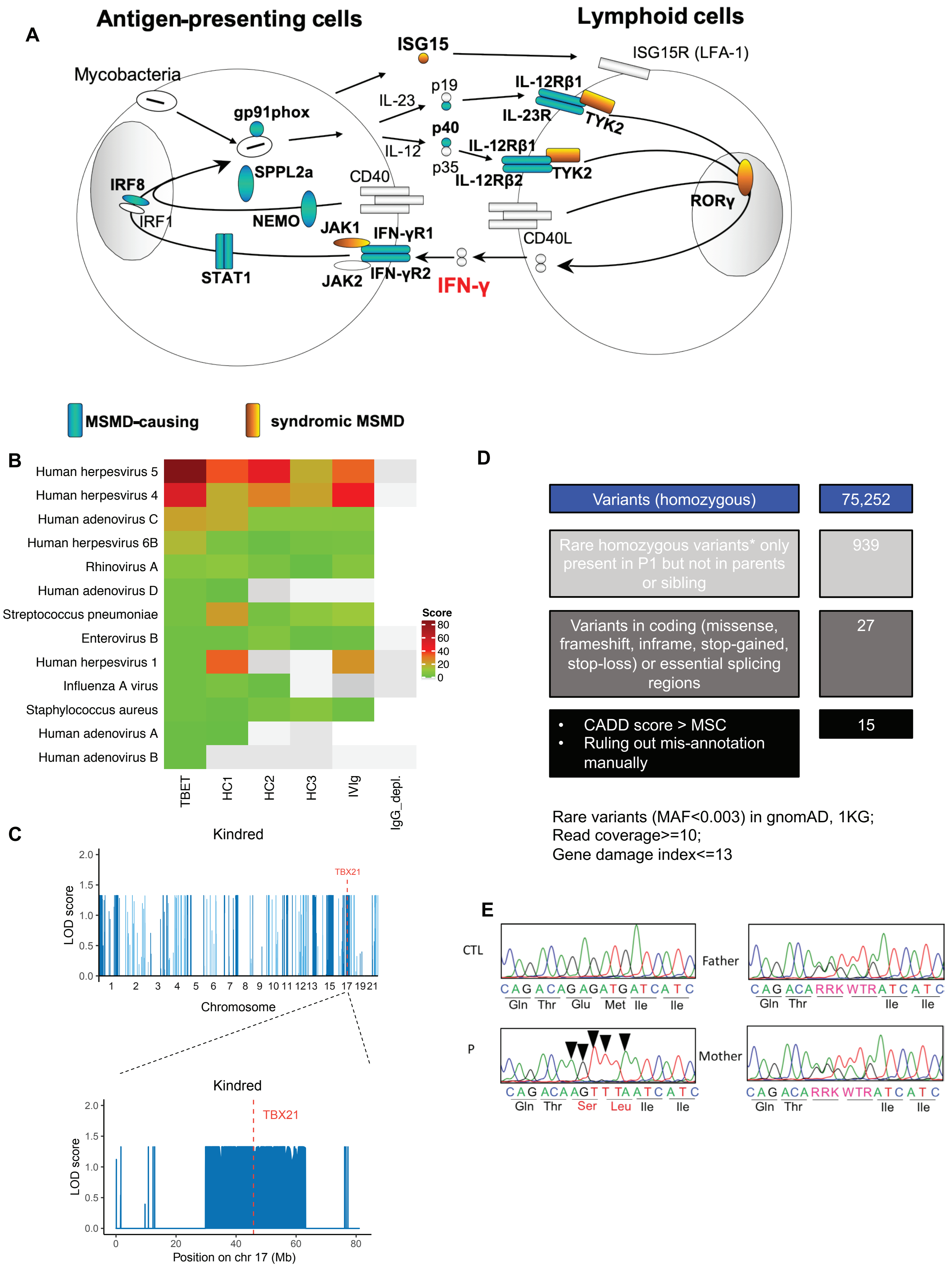
A
Antigen-presenting cells
Lymphoid cells
Mycobacteria
gp91phox
SPPL2a
IRF8
IRF1
NEMO
STAT1
CD40
JAK1
JAK2
IFN-γR1
IFN-γR2
IFN-γ
ISG15
ISG15R (LFA-1)
IL-23
IL-12
p19
p40
p35
IL-12Rβ1
IL-23R
TYK2
IL-12Rβ1
IL-12Rβ2
TYK2
CD40L
RORγ
MSMD-causing
syndromic MSMD
B
Human herpesvirus 5
Human herpesvirus 4
Human adenovirus C
Human herpesvirus 6B
Rhinovirus A
Human adenovirus D
Streptococcus pneumoniae
Enterovirus B
Human herpesvirus 1
Influenza A virus
Staphylococcus aureus
Human adenovirus A
Human adenovirus B
TBET
HC1
HC2
HC3
IVIg
IgG_depl.
Score
80
60
40
20
0
C
Kindred
LOD score
TBX21
Chromosome
1 2 3 4 5 6 7 8 9 10 11 12 13 15 17 19 21
Kindred
TBX21
LOD score
Position on chr 17 (Mb)
0 20 40 60 80
D
Variants (homozygous)
75,252
Rare homozygous variants* only present in P1 but not in parents or sibling
939
Variants in coding (missense, frameshift, inframe, stop-gained, stop-loss) or essential splicing regions
27
CADD score > MSC
Ruling out mis-annotation manually
15
Rare variants (MAF<0.003) in gnomAD, 1KG;
Read coverage>=10;
Gene damage index<=13
E
CTL
CAG ACAG AGATGATCATC
Gln Thr Glu Met Ile Ile
Father
CAG ACA RRK WTR ATC ATC
Gln Thr Ile Ile
P
CAG ACAAGTT TAATC ATC
Gln Thr Ser Leu Ile Ile
Mother
CAG ACA RRK WTR ATC ATC
Gln Thr Ile Ile

**Figure S1**

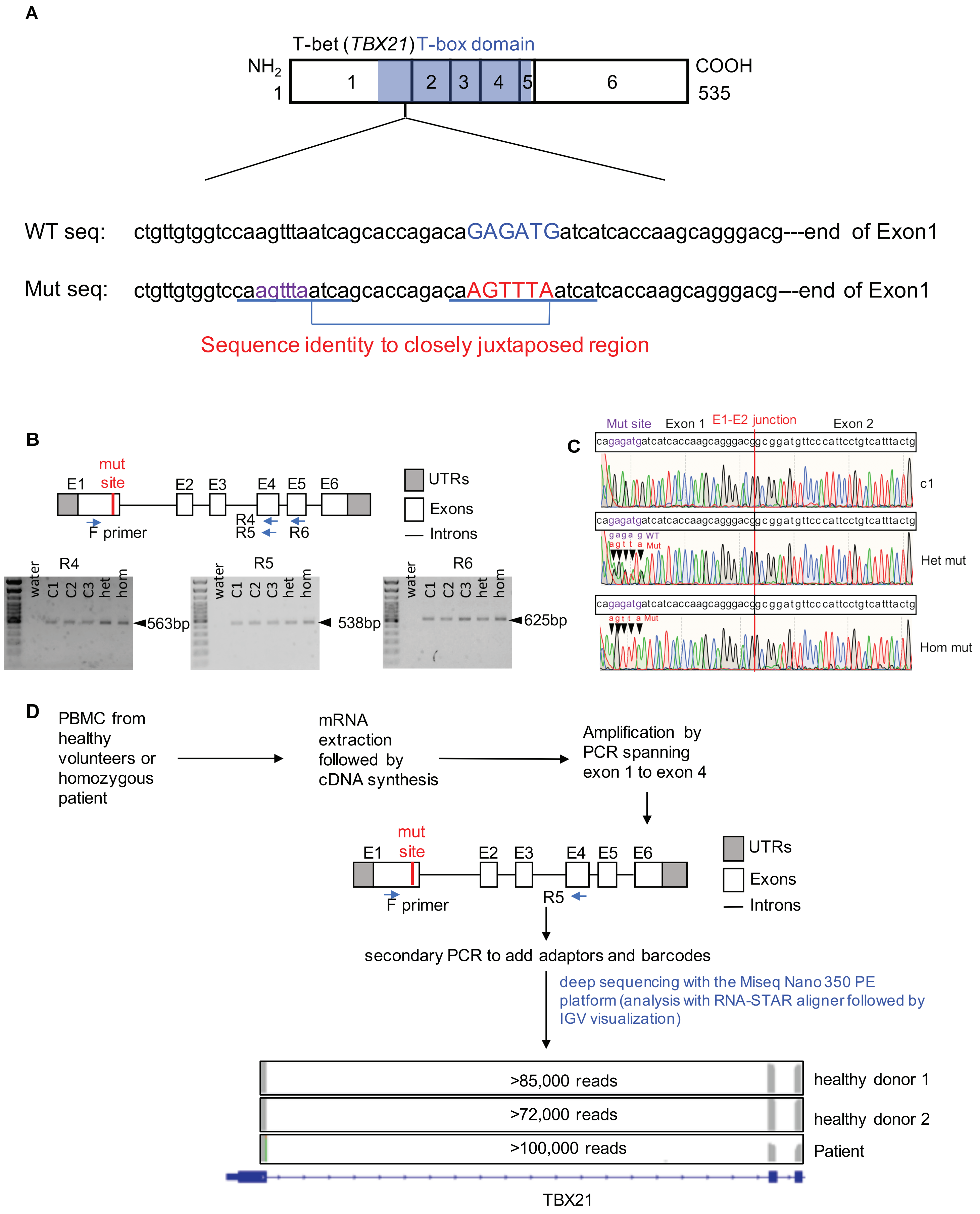
A
T-bet (TBX21) T-box domain
NH2
1
1
2
3
4
5
6
COOH
535
WT seq: ctgttgtggtccaagtttaatcagcaccagacaGAGATGatcatcaccaagcagggacg---end of Exon1
Mut seq: ctgttgtggtccaagtttaatcagcaccagacaAGTTTAatcatcaccaagcagggacg---end of Exon1
Sequence identity to closely juxtaposed region
B
mut site
E1
E2
E3
E4
E5
E6
F primer
R4
R5
R6
UTRs
Exons
Introns
R4
water
C1
C2
C3
het
hom
563bp
R5
538bp
R6
625bp
C
Mut site
Exon 1
E1-E2 junction
Exon 2
cagagatgatcatcaccaagcagggacggcggatgttcccattcctgtcatttactg
c1
gaga g WT
agtt a Mut
Het mut
Hom mut
D
PBMC from healthy volunteers or homozygous patient
mRNA extraction followed by cDNA synthesis
Amplification by PCR spanning exon 1 to exon 4
secondary PCR to add adaptors and barcodes
deep sequencing with the Miseq Nano 350 PE platform (analysis with RNA-STAR aligner followed by IGV visualization)
>85,000 reads
healthy donor 1
>72,000 reads
healthy donor 2
>100,000 reads
Patient
TBX21

**Figure S2**

**A**

| T-box family paralogs | Amino acids |
|---|---|
| sp\|O43435\|TBX1_HUMAN | GT**EM**IVTK |
| sp\|Q13207\|TBX2_HUMAN | GT**EM**VITK |
| sp\|O15119\|TBX3_HUMAN | GT**EM**VITK |
| sp\|P57082\|TBX4_HUMAN | GT**EM**IITK |
| sp\|Q99593\|TBX5_HUMAN | GT**EM**IITK |
| sp\|O95947\|TBX6_HUMAN | GT**EM**IITK |
| sp\|O75333\|TBX10_HUMAN | GT**EM**IVTK |
| sp\|Q96SF7\|TBX15_HUMAN | GT**EM**IITK |
| sp\|O95935\|TBX18_HUMAN | GT**EM**IITK |
| sp\|O60806\|TBX19_HUMAN | TN**EM**IVTK |
| sp\|Q9UMR3\|TBX20_HUMAN | GT**EM**IITK |
| sp\|Q9Y458\|TBX22_HUMAN | GT**EM**IITK |
| sp\|O95936\|EOMES_HUMAN | QT**EM**IITK |
| sp\|Q8IWI9\|MGAP_HUMAN | ST**EM**ILTK |
| sp\|O15178\|TBXT_HUMAN | TN**EM**IVTK |
| sp\|Q16650\|TBR1_HUMAN | QT**EM**IITK |
| sp\|Q9UL17\|TBX21_HUMAN | QT**EM**IITK |
| | .**:::** |

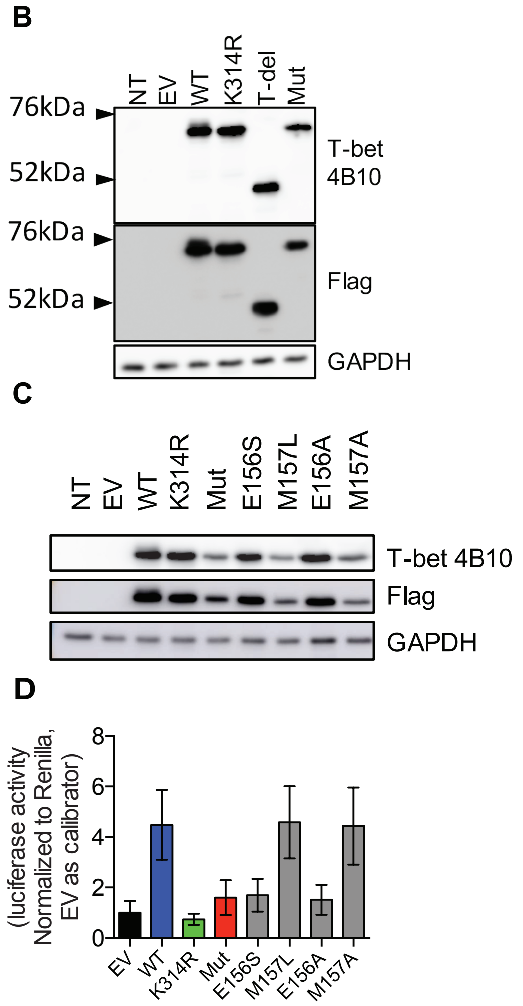


**Figure S3**

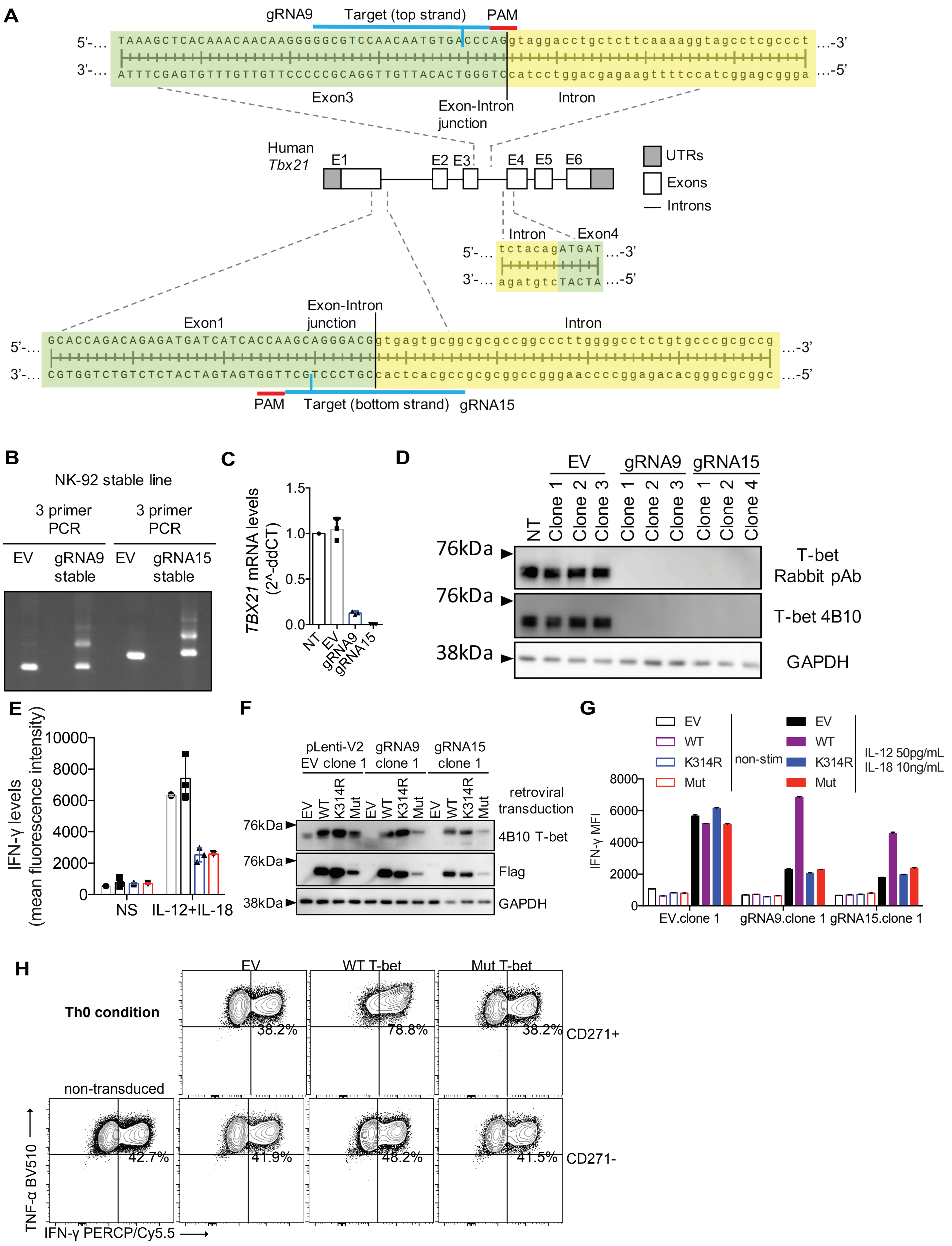
A
gRNA9
Target (top strand)
PAM
5'-… TAAAGCTCACAAACAACAAGGGGGCGTCCAACAATGTGACCCAGgtaggacctgctcttcaaaaggtagcctcgccct …-3'
3'-… ATTTCGAGTGTTTGTTGTTCCCCCGCAGGTTGTTACACTGGGTCcatcctggacgagaagtttccatcggagcggga …-5'
Exon3
Exon-Intron junction
Intron
Human Tbx21
E1
E2 E3
E4 E5 E6
UTRs
Exons
Introns
Intron
Exon4
5'-… tctacagATGAT …-3'
3'-… agatgtcTACTA …-5'
Exon1
Exon-Intron junction
Intron
5'-… GCACCAGACAGAGATGATCATCACCAAGCAGGGACGgtgagtgcggcgcgccggcccttggggcctctgtgcccgcgccg …-3'
3'-… CGTGGTCTGTCTCTACTAGTAGTGGTTCGTCCCTGCcactcacgccgcgcggccgggaaccccggagacacgggcgcggc …-5'
PAM
Target (bottom strand)
gRNA15
B
NK-92 stable line
3 primer PCR
3 primer PCR
EV
gRNA9 stable
EV
gRNA15 stable
C
TBX21 mRNA levels (2^-ddCT)
1.5
1.0
0.5
0.0
NT
EV
gRNA9
gRNA15
D
EV
gRNA9
gRNA15
NT
Clone 1
Clone 2
Clone 3
Clone 1
Clone 2
Clone 3
Clone 1
Clone 2
Clone 4
76kDa
76kDa
38kDa
T-bet Rabbit pAb
T-bet 4B10
GAPDH
E
IFN-γ levels (mean fluorescence intensity)
10000
8000
6000
4000
2000
0
NS
IL-12+IL-18
F
pLenti-V2 EV clone 1
gRNA9 clone 1
gRNA15 clone 1
EV
WT
K314R
Mut
EV
WT
K314R
Mut
EV
WT
K314R
Mut
retroviral transduction
76kDa
76kDa
38kDa
4B10 T-bet
Flag
GAPDH
G
EV
WT
K314R
Mut
non-stim
EV
WT
K314R
Mut
IL-12 50pg/mL
IL-18 10ng/mL
IFN-γ MFI
8000
6000
4000
2000
0
EV.clone 1
gRNA9.clone 1
gRNA15.clone 1
H
EV
WT T-bet
Mut T-bet
Th0 condition
38.2%
78.8%
38.2%
CD271+
non-transduced
42.7%
41.9%
48.2%
41.5%
CD271-
TNF-α BV510
IFN-γ PERCP/Cy5.5

**Figure S4**

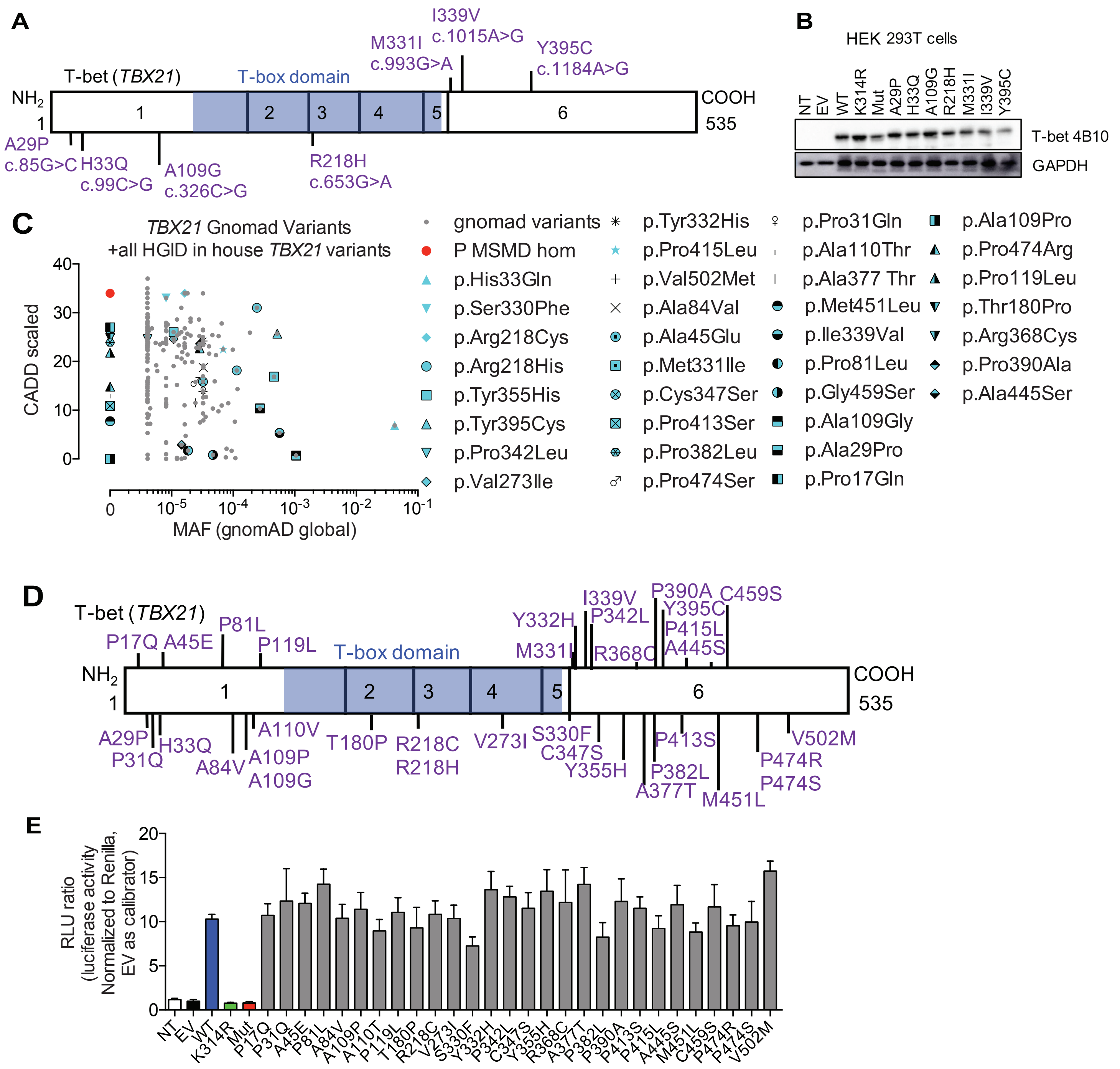
A
T-bet (TBX21)
T-box domain
NH2
1
COOH
535
1 2 3 4 5 6
A29P c.85G>C
H33Q c.99C>G
A109G c.326C>G
R218H c.653G>A
M331I c.993G>A
I339V c.1015A>G
Y395C c.1184A>G
B
HEK 293T cells
NT EV WT K314R Mut A29P H33Q A109G R218H M331I I339V Y395C
T-bet 4B10
GAPDH
C
TBX21 Gnomad Variants
+all HGID in house TBX21 variants
CADD scaled
MAF (gnomAD global)
gnomad variants
P MSMD hom
p.His33Gln
p.Ser330Phe
p.Arg218Cys
p.Arg218His
p.Tyr355His
p.Tyr395Cys
p.Pro342Leu
p.Val273Ile
p.Tyr332His
p.Pro415Leu
p.Val502Met
p.Ala84Val
p.Ala45Glu
p.Met331Ile
p.Cys347Ser
p.Pro413Ser
p.Pro382Leu
p.Pro474Ser
p.Pro31Gln
p.Ala110Thr
p.Ala377 Thr
p.Met451Leu
p.Ile339Val
p.Pro81Leu
p.Gly459Ser
p.Ala109Gly
p.Ala29Pro
p.Pro17Gln
p.Ala109Pro
p.Pro474Arg
p.Pro119Leu
p.Thr180Pro
p.Arg368Cys
p.Pro390Ala
p.Ala445Ser
D
T-bet (TBX21)
T-box domain
P17Q A45E P81L P119L
M331I Y332H I339V P342L R368C P390A Y395C P415L A445S C459S
A29P P31Q H33Q A84V A110V A109P A109G T180P R218C R218H V273I S330F C347S Y355H P382L A377T P413S M451L P474R P474S V502M
E
RLU ratio
(luciferase activity
Normalized to Renilla,
EV as calibrator)
NT EV WT K314R Mut P17Q P31Q A45E P81L A84V A109P A110T P119L T180P R218C V273I S330F Y332H P342L C347S Y355H R368C A377T P382L P390A P413S P415L A445S M451L C459S P474R P474S V502M

**Figure S5**

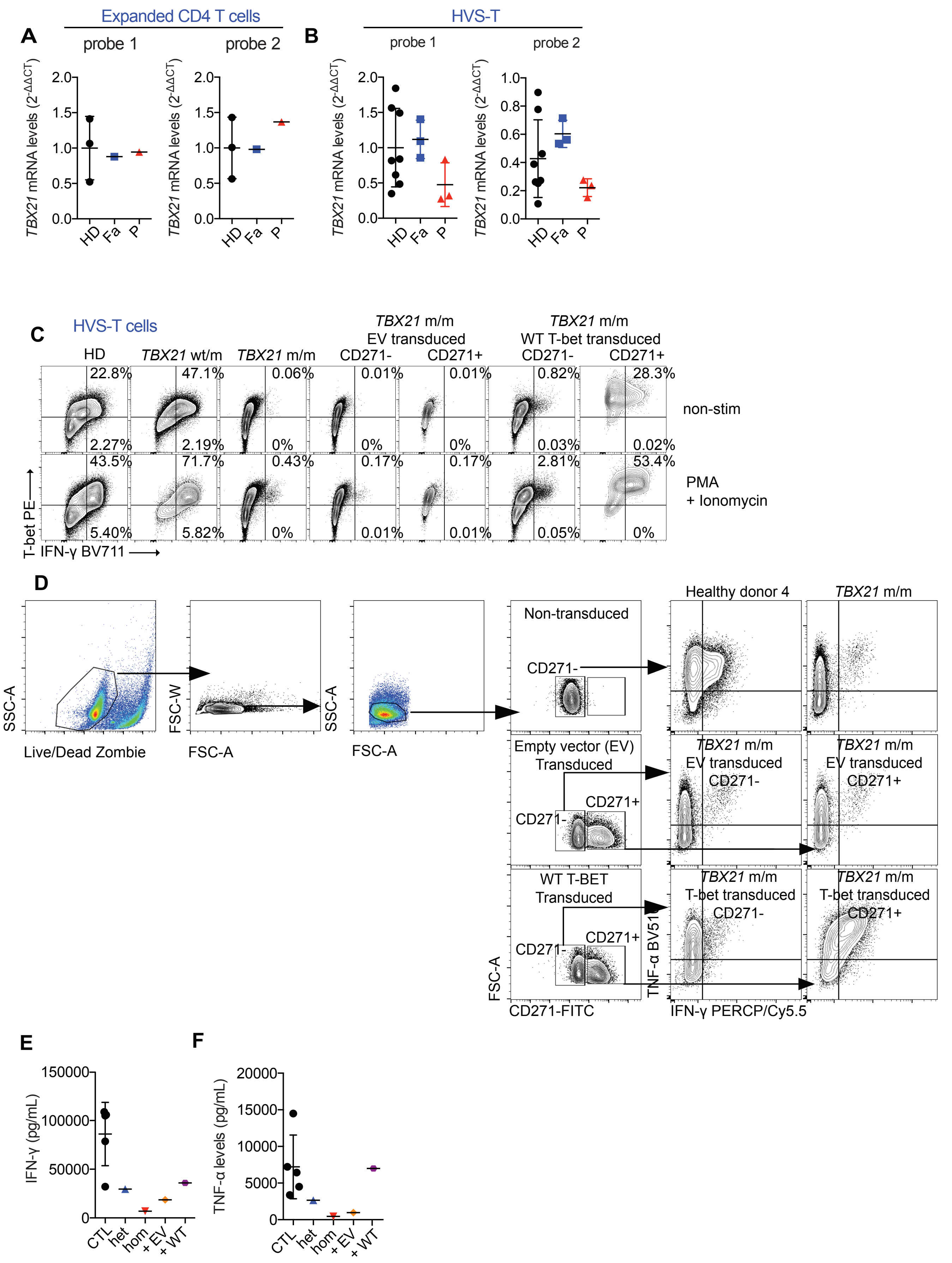
A
Expanded CD4 T cells
probe 1
probe 2
B
HVS-T
probe 1
probe 2
TBX21 mRNA levels (2-ΔΔCT)
HD
Fa
P
C
HVS-T cells
HD
TBX21 wt/m
TBX21 m/m
TBX21 m/m
EV transduced
CD271-
CD271+
TBX21 m/m
WT T-bet transduced
CD271-
CD271+
non-stim
PMA
+ Ionomycin
T-bet PE
IFN-γ BV711
D
SSC-A
Live/Dead Zombie
FSC-W
FSC-A
SSC-A
FSC-A
Non-transduced
CD271-
Empty vector (EV)
Transduced
CD271-
CD271+
WT T-BET
Transduced
CD271-
CD271+
FSC-A
CD271-FITC
Healthy donor 4
TBX21 m/m
TBX21 m/m
EV transduced
CD271-
TBX21 m/m
EV transduced
CD271+
TBX21 m/m
T-bet transduced
CD271-
TBX21 m/m
T-bet transduced
CD271+
TNF-α BV510
IFN-γ PERCP/Cy5.5
E
IFN-γ (pg/mL)
CTL
het
hom
+ EV
+ WT
F
TNF-α levels (pg/mL)
CTL
het
hom
+ EV
+ WT

**Figure S6**

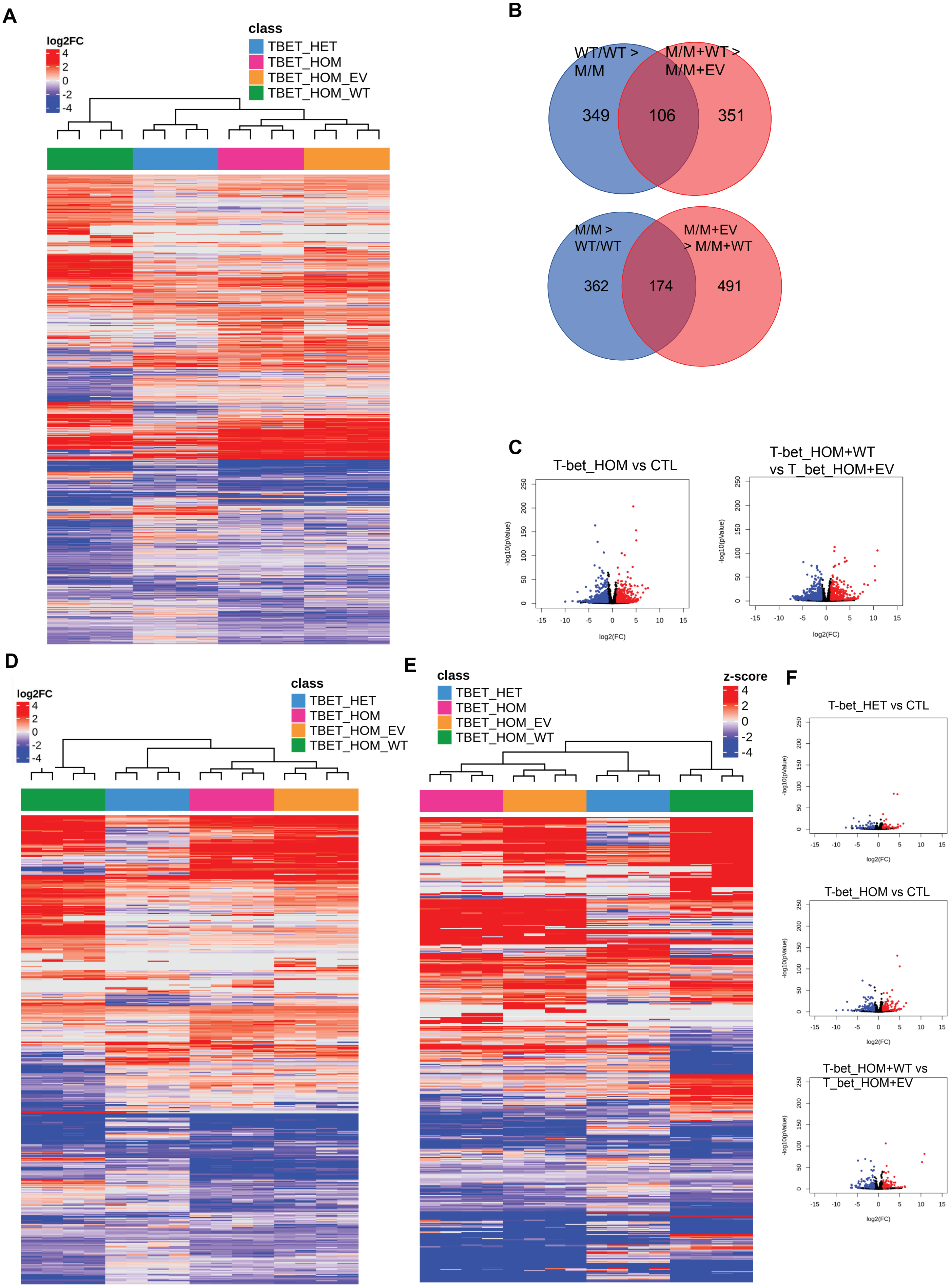
A
log2FC
4
2
0
-2
-4
class
TBET_HET
TBET_HOM
TBET_HOM_EV
TBET_HOM_WT
B
WT/WT > M/M
M/M+WT > M/M+EV
349
106
351
M/M > WT/WT
M/M+EV > M/M+WT
362
174
491
C
T-bet_HOM vs CTL
T-bet_HOM+WT vs T_bet_HOM+EV
-log10(pValue)
log2(FC)
D
log2FC
class
TBET_HET
TBET_HOM
TBET_HOM_EV
TBET_HOM_WT
E
class
TBET_HET
TBET_HOM
TBET_HOM_EV
TBET_HOM_WT
z-score
F
T-bet_HET vs CTL
T-bet_HOM vs CTL
T-bet_HOM+WT vs T_bet_HOM+EV

**Figure S7**

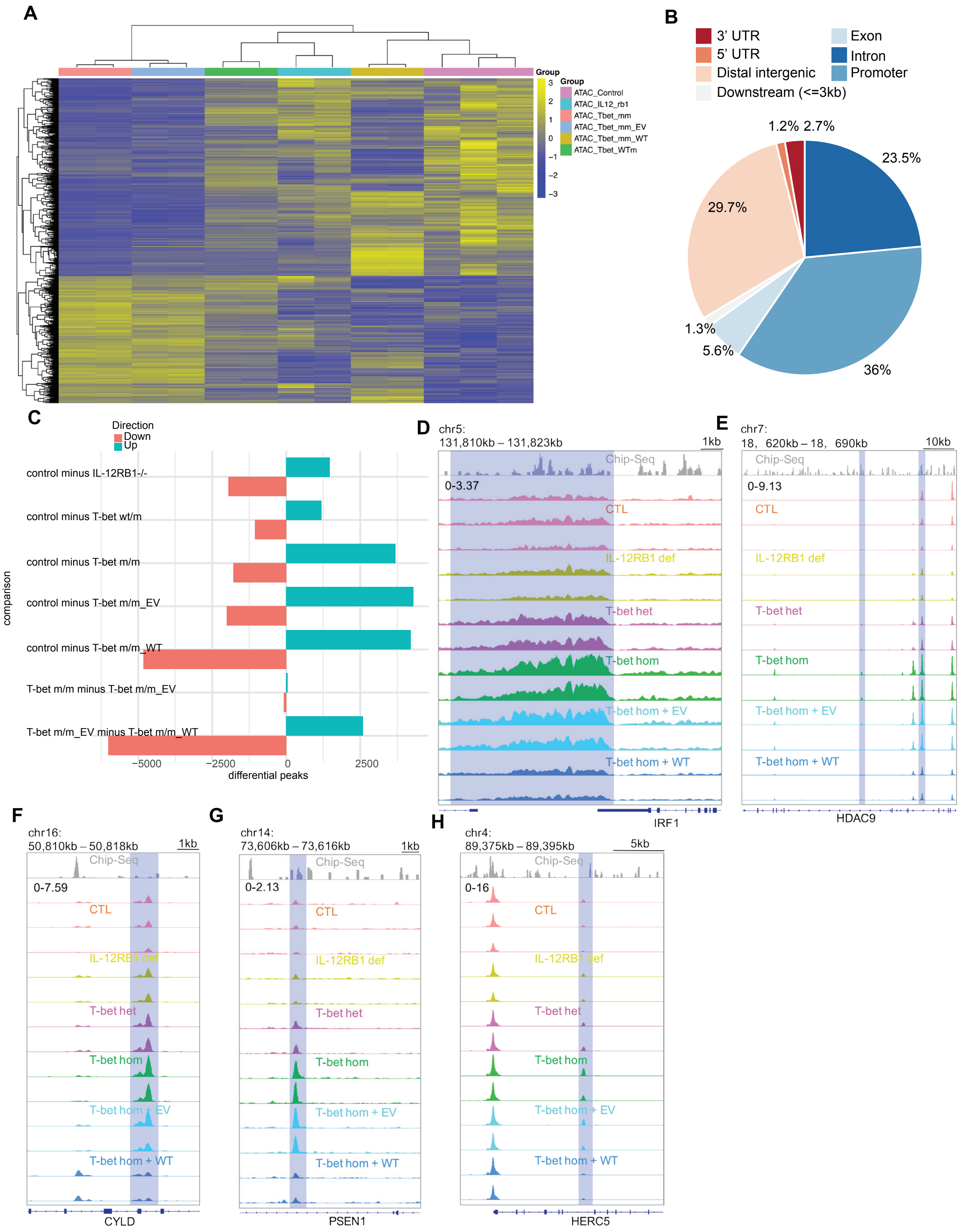
A
Group
ATAC_Control
ATAC_IL12_rb1
ATAC_Tbet_mm
ATAC_Tbet_mm_EV
ATAC_Tbet_mm_WT
ATAC_Tbet_WTm
B
3' UTR
5' UTR
Distal intergenic
Downstream (<=3kb)
Exon
Intron
Promoter
1.2% 2.7%
23.5%
29.7%
1.3%
5.6%
36%
C
Direction
Down
Up
control minus IL-12RB1-/-
control minus T-bet wt/m
control minus T-bet m/m
control minus T-bet m/m_EV
control minus T-bet m/m_WT
T-bet m/m minus T-bet m/m_EV
T-bet m/m_EV minus T-bet m/m_WT
comparison
-5000 -2500 0 2500
differential peaks
D
chr5:
131,810kb – 131,823kb
1kb
Chip-Seq
0-3.37
CTL
IL-12RB1 def
T-bet het
T-bet hom
T-bet hom + EV
T-bet hom + WT
IRF1
E
chr7:
18, 620kb – 18, 690kb
10kb
Chip-Seq
0-9.13
CTL
IL-12RB1 def
T-bet het
T-bet hom
T-bet hom + EV
T-bet hom + WT
HDAC9
F
chr16:
50,810kb – 50,818kb
1kb
Chip-Seq
0-7.59
CTL
IL-12RB1 def
T-bet het
T-bet hom
T-bet hom + EV
T-bet hom + WT
CYLD
G
chr14:
73,606kb – 73,616kb
1kb
Chip-Seq
0-2.13
CTL
IL-12RB1 def
T-bet het
T-bet hom
T-bet hom + EV
T-bet hom + WT
PSEN1
H
chr4:
89,375kb – 89,395kb
5kb
Chip-Seq
0-16
CTL
IL-12RB1 def
T-bet het
T-bet hom
T-bet hom + EV
T-bet hom + WT
HERC5

**Figure S8**

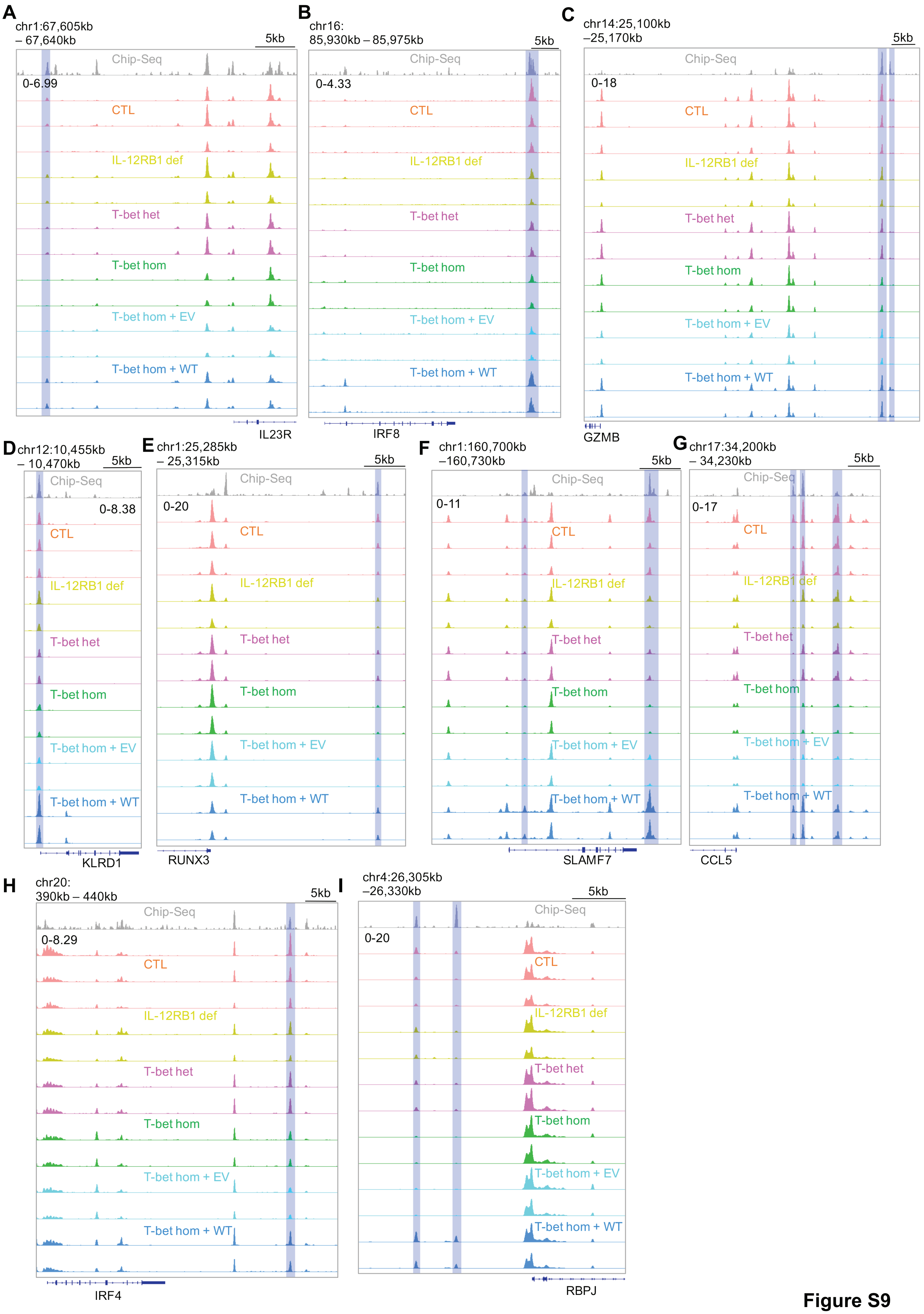

A
chr1:67,605kb – 67,640kb
5kb
Chip-Seq
0-6.99
CTL
IL-12RB1 def
T-bet het
T-bet hom
T-bet hom + EV
T-bet hom + WT
IL23R
B
chr16: 85,930kb – 85,975kb
5kb
Chip-Seq
0-4.33
CTL
IL-12RB1 def
T-bet het
T-bet hom
T-bet hom + EV
T-bet hom + WT
IRF8
C
chr14:25,100kb –25,170kb
5kb
Chip-Seq
0-18
CTL
IL-12RB1 def
T-bet het
T-bet hom
T-bet hom + EV
T-bet hom + WT
GZMB
D
chr12:10,455kb – 10,470kb
5kb
Chip-Seq
0-8.38
CTL
IL-12RB1 def
T-bet het
T-bet hom
T-bet hom + EV
T-bet hom + WT
KLRD1
E
chr1:25,285kb – 25,315kb
5kb
Chip-Seq
0-20
CTL
IL-12RB1 def
T-bet het
T-bet hom
T-bet hom + EV
T-bet hom + WT
RUNX3
F
chr1:160,700kb –160,730kb
5kb
Chip-Seq
0-11
CTL
IL-12RB1 def
T-bet het
T-bet hom
T-bet hom + EV
T-bet hom + WT
SLAMF7
G
chr17:34,200kb – 34,230kb
5kb
Chip-Seq
0-17
CTL
IL-12RB1 def
T-bet het
T-bet hom
T-bet hom + EV
T-bet hom + WT
CCL5
H
chr20: 390kb – 440kb
5kb
Chip-Seq
0-8.29
CTL
IL-12RB1 def
T-bet het
T-bet hom
T-bet hom + EV
T-bet hom + WT
IRF4
I
chr4:26,305kb –26,330kb
5kb
Chip-Seq
0-20
CTL
IL-12RB1 def
T-bet het
T-bet hom
T-bet hom + EV
T-bet hom + WT
RBPJ


**Figure S9**

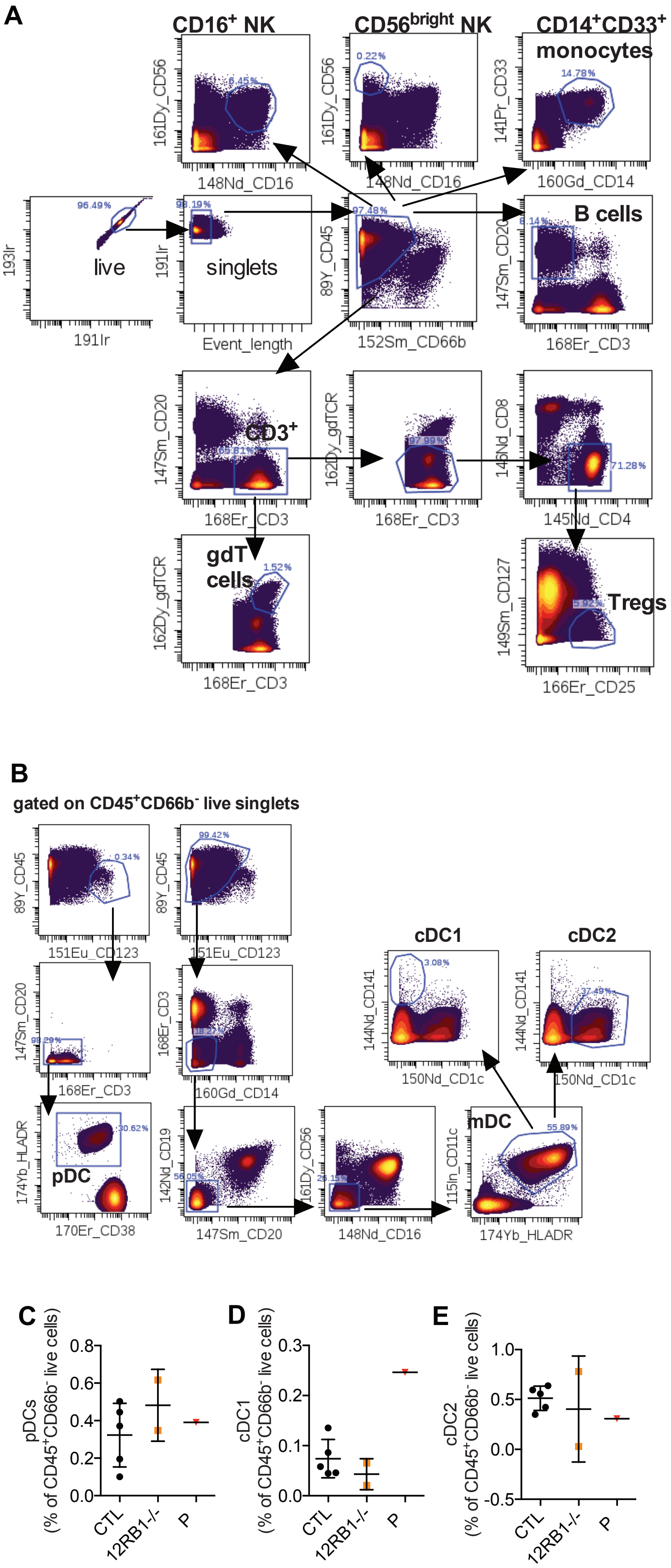
A
CD16+ NK
CD56bright NK
CD14+CD33+ monocytes
B cells
live
singlets
CD3+
gdT cells
Tregs
193Ir
191Ir
Event_length
161Dy_CD56
148Nd_CD16
141Pr_CD33
160Gd_CD14
89Y_CD45
152Sm_CD66b
147Sm_CD20
168Er_CD3
162Dy_gdTCR
145Nd_CD8
145Nd_CD4
149Sm_CD127
166Er_CD25
96.49%
98.19%
97.48%
6.45%
0.22%
14.78%
8.14%
65.81%
97.99%
71.28%
1.52%
5.92%
B
gated on CD45+CD66b- live singlets
cDC1
cDC2
mDC
pDC
89Y_CD45
151Eu_CD123
147Sm_CD20
168Er_CD3
160Gd_CD14
144Nd_CD141
150Nd_CD1c
174Yb_HLADR
170Er_CD38
142Nd_CD19
161Dy_CD56
148Nd_CD16
115In_CD11c
0.34%
99.42%
98.29%
3.08%
37.49%
30.62%
56.05%
55.89%
C
D
E
pDCs (% of CD45+CD66b- live cells)
cDC1 (% of CD45+CD66b- live cells)
cDC2 (% of CD45+CD66b- live cells)
0.8
0.6
0.4
0.2
0.0
0.3
0.1
1.0
0.5
-0.5
CTL
12RB1-/-
P

Figure S10

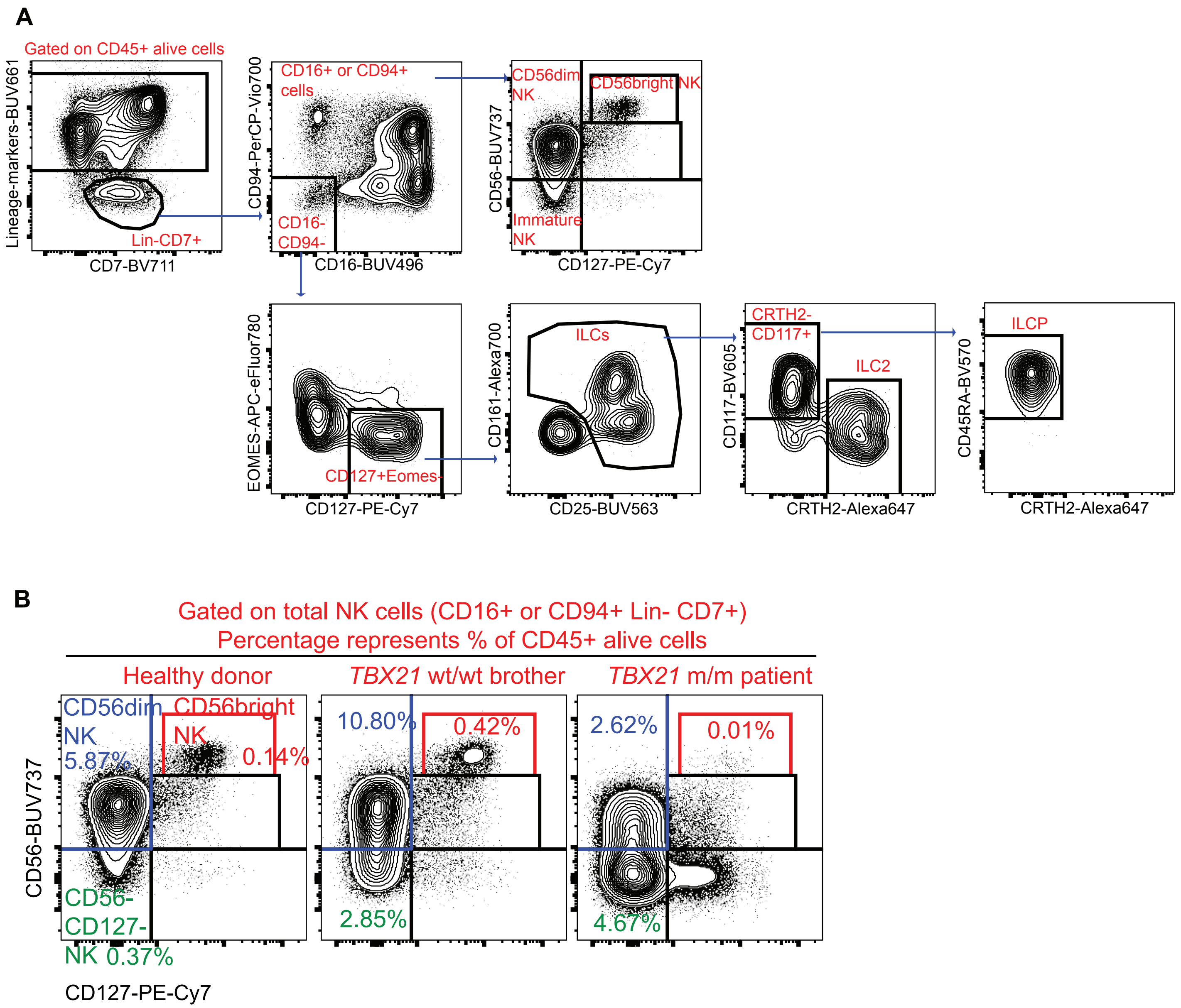
A
Gated on CD45+ alive cells
Lineage-markers-BUV661
CD7-BV711
Lin-CD7+
CD16+ or CD94+ cells
CD94-PerCP-Vio700
CD16-BUV496
CD16- CD94-
CD56dim NK
CD56bright NK
Immature NK
CD56-BUV737
CD127-PE-Cy7
EOMES-APC-eFluor780
CD127+Eomes-
ILCs
CD161-Alexa700
CD25-BUV563
CRTH2- CD117+
ILC2
CD117-BV605
CRTH2-Alexa647
ILCP
CD45RA-BV570
CRTH2-Alexa647
B
Gated on total NK cells (CD16+ or CD94+ Lin- CD7+)
Percentage represents % of CD45+ alive cells
Healthy donor
TBX21 wt/wt brother
TBX21 m/m patient
CD56dim NK 5.87%
CD56bright NK 0.14%
CD56- CD127- NK 0.37%
10.80%
0.42%
2.85%
2.62%
0.01%
4.67%
CD56-BUV737
CD127-PE-Cy7

**Figure S11**

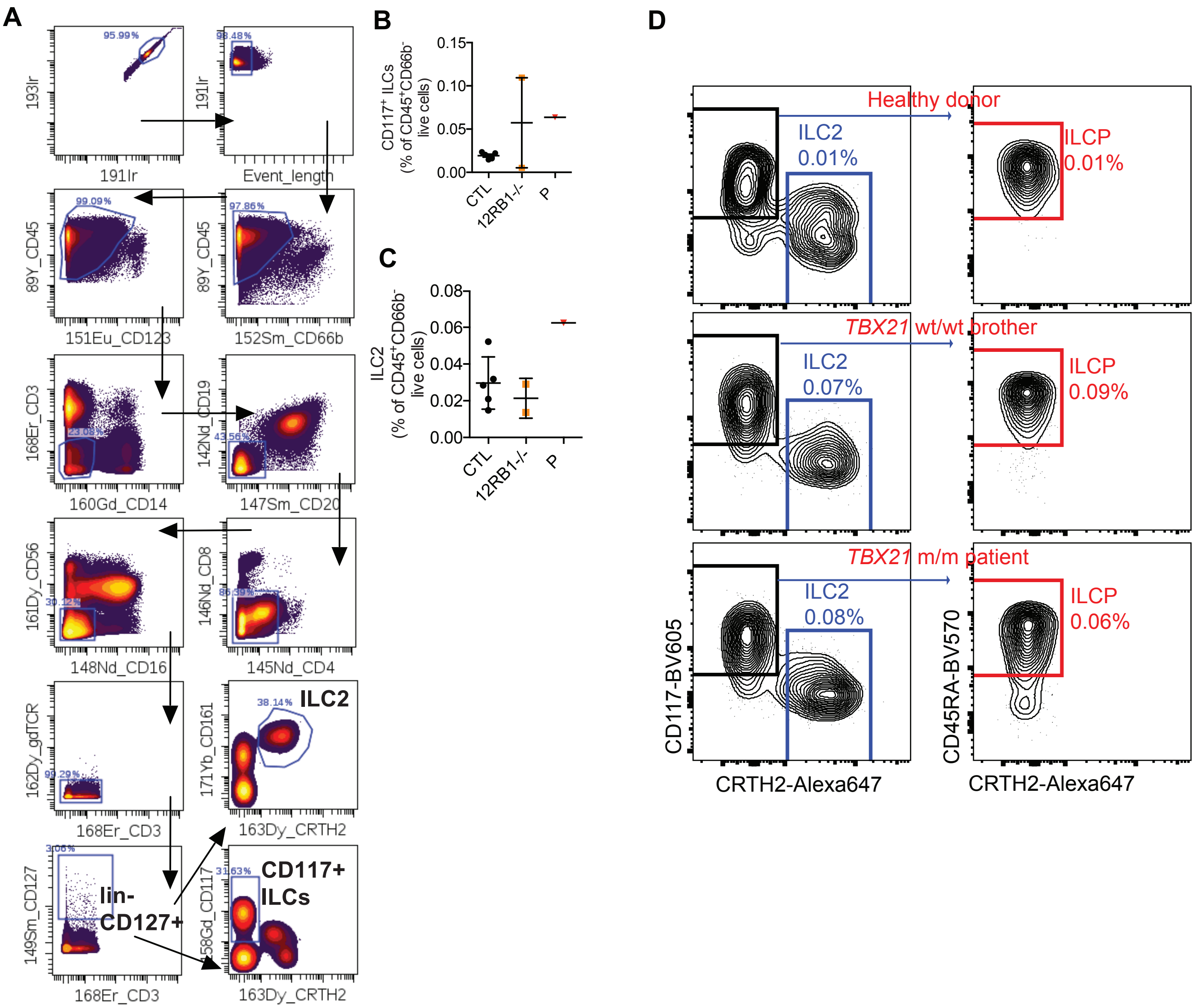

A
B
C
D
95.99%
193Ir
191Ir
98.48%
Event_length
99.09%
89Y_CD45
151Eu_CD123
97.86%
152Sm_CD66b
168Er_CD3
23.08%
160Gd_CD14
142Nd_CD19
43.56%
147Sm_CD20
161Dy_CD56
30.12%
148Nd_CD16
146Nd_CD8
86.39%
145Nd_CD4
162Dy_gdTCR
99.29%
168Er_CD3
171Yb_CD161
38.14%
ILC2
163Dy_CRTH2
149Sm_CD127
3.06%
lin-
CD127+
158Gd_CD117
31.63%
CD117+
ILCs
CD117+ ILCs
(% of CD45+CD66b- live cells)
0.15
0.10
0.05
0.00
CTL
12RB1-/-
P
ILC2
(% of CD45+CD66b- live cells)
0.08
0.06
0.04
0.02
0.00
Healthy donor
ILC2
0.01%
ILCP
0.01%
TBX21 wt/wt brother
ILC2
0.07%
ILCP
0.09%
TBX21 m/m patient
ILC2
0.08%
ILCP
0.06%
CD117-BV605
CRTH2-Alexa647
CD45RA-BV570
CRTH2-Alexa647


Figure S12

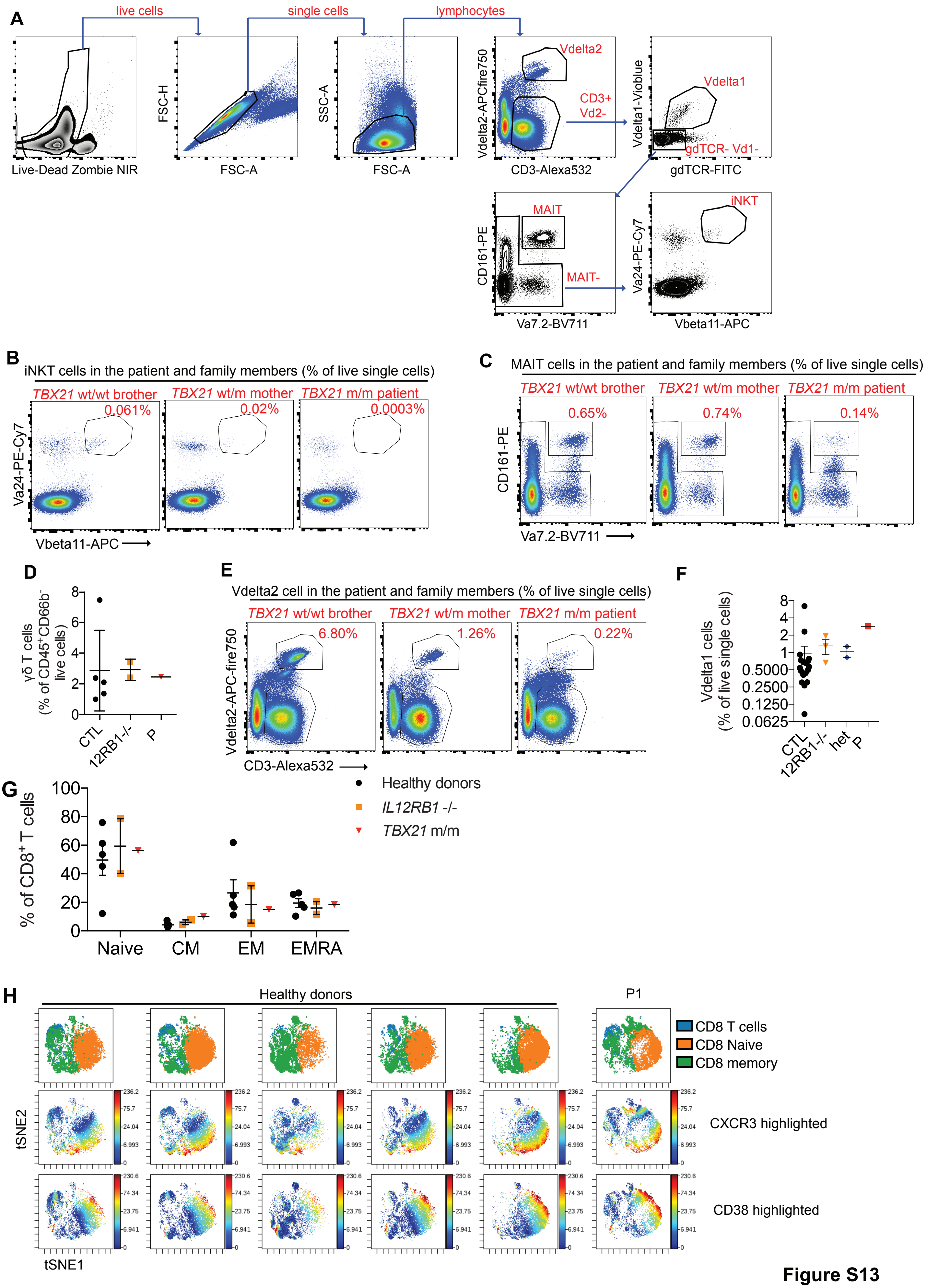
A
live cells
single cells
lymphocytes
Live-Dead Zombie NIR
FSC-H
FSC-A
SSC-A
FSC-A
Vdelta2-APCfire750
Vdelta2
CD3+
Vd2-
CD3-Alexa532
Vdelta1-Vioblue
Vdelta1
gdTCR- Vd1-
gdTCR-FITC
CD161-PE
MAIT
MAIT-
Va7.2-BV711
Va24-PE-Cy7
iNKT
Vbeta11-APC
B
iNKT cells in the patient and family members (% of live single cells)
TBX21 wt/wt brother
TBX21 wt/m mother
TBX21 m/m patient
0.061%
0.02%
0.0003%
Va24-PE-Cy7
Vbeta11-APC
C
MAIT cells in the patient and family members (% of live single cells)
TBX21 wt/wt brother
TBX21 wt/m mother
TBX21 m/m patient
0.65%
0.74%
0.14%
CD161-PE
Va7.2-BV711
D
γδ T cells (% of CD45+CD66b- live cells)
CTL
12RB1-/-
P
E
Vdelta2 cell in the patient and family members (% of live single cells)
TBX21 wt/wt brother
TBX21 wt/m mother
TBX21 m/m patient
6.80%
1.26%
0.22%
Vdelta2-APC-fire750
CD3-Alexa532
F
Vdelta1 cells (% of live single cells)
CTL
12RB1-/-
het
P
G
% of CD8+ T cells
Naive
CM
EM
EMRA
Healthy donors
IL12RB1 -/-
TBX21 m/m
H
Healthy donors
P1
CD8 T cells
CD8 Naive
CD8 memory
tSNE2
tSNE1
CXCR3 highlighted
CD38 highlighted

**Figure S13**

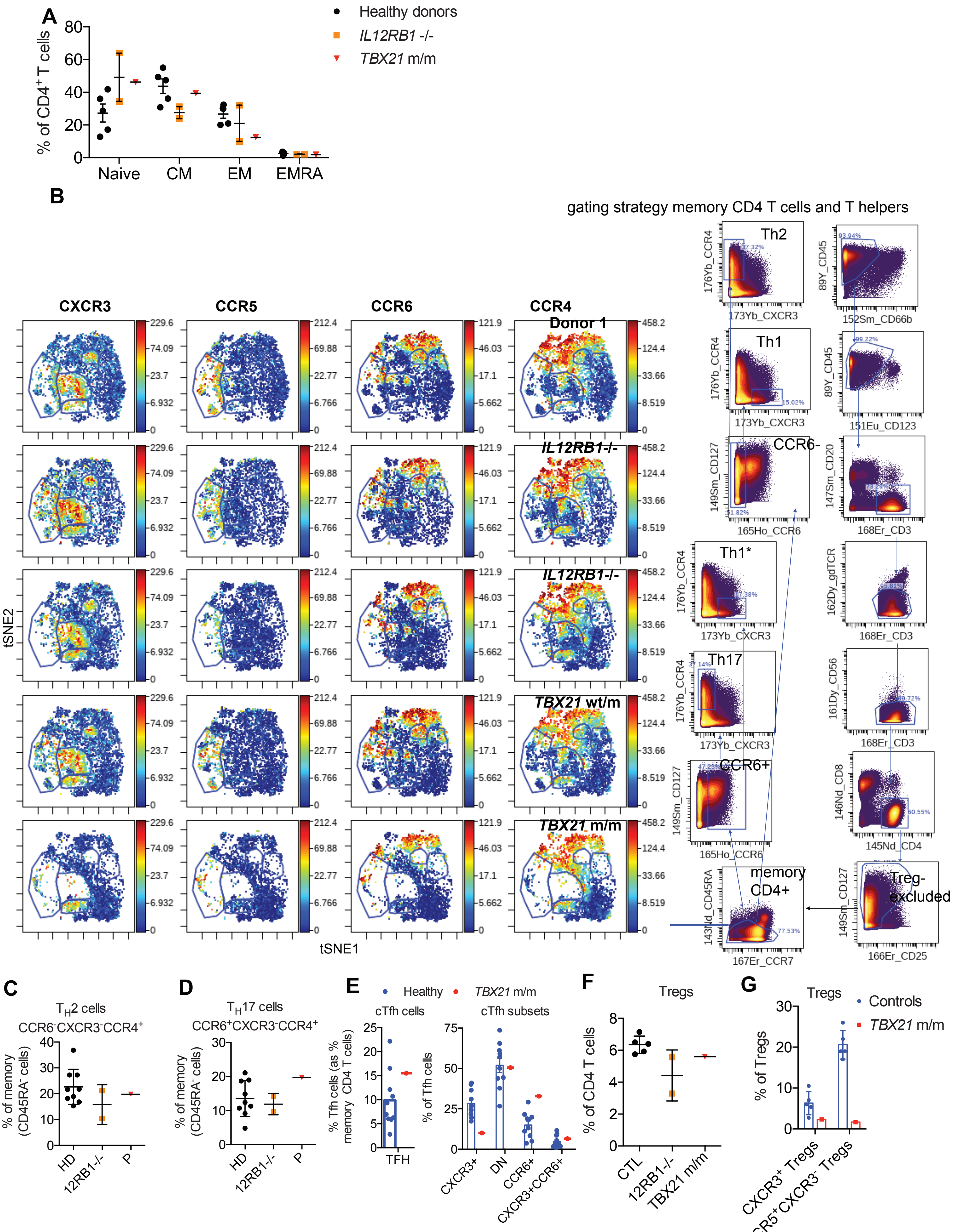

A
Healthy donors
IL12RB1 -/-
TBX21 m/m
% of CD4+ T cells
Naive
CM
EM
EMRA
B
gating strategy memory CD4 T cells and T helpers
CXCR3
CCR5
CCR6
CCR4
Donor 1
IL12RB1-/-
IL12RB1-/-
TBX21 wt/m
TBX21 m/m
tSNE2
tSNE1
Th2
Th1
CCR6-
Th1*
Th17
CCR6+
memory CD4+
Treg-excluded
C
TH2 cells
CCR6-CXCR3-CCR4+
% of memory (CD45RA- cells)
HD
12RB1-/-
P
D
TH17 cells
CCR6+CXCR3-CCR4+
% of memory (CD45RA- cells)
HD
12RB1-/-
P
E
Healthy
TBX21 m/m
cTfh cells
cTfh subsets
% Tfh cells (as % memory CD4 T cells)
TFH
% of Tfh cells
CXCR3+
DN
CCR6+
CXCR3+CCR6+
F
Tregs
% of CD4 T cells
CTL
12RB1-/-
TBX21 m/m
G
Tregs
Controls
TBX21 m/m
% of Tregs
CXCR3+ Tregs
CCR5+CXCR3- Tregs


**Figure S14**

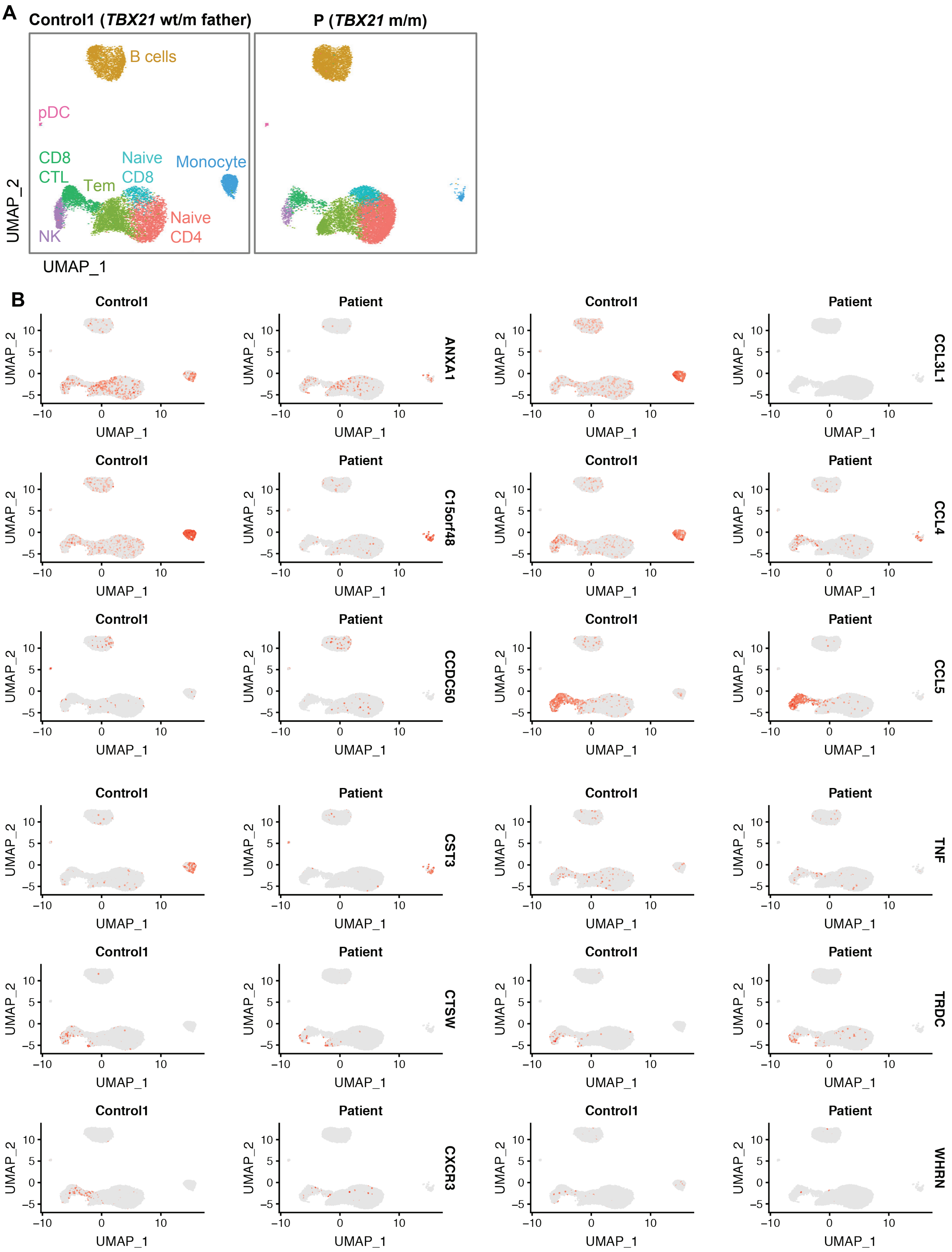
A
Control1 (TBX21 wt/m father)
P (TBX21 m/m)
B cells
pDC
CD8
CTL
Tem
Naive
CD8
Monocyte
Naive
CD4
NK
UMAP_2
UMAP_1
B
Control1
Patient
UMAP_2
UMAP_1
ANXA1
CCL3L1
C15orf48
CCL4
CCDC50
CCL5
CST3
TNF
CTSW
TRDC
CXCR3
WHRN

Figure S15

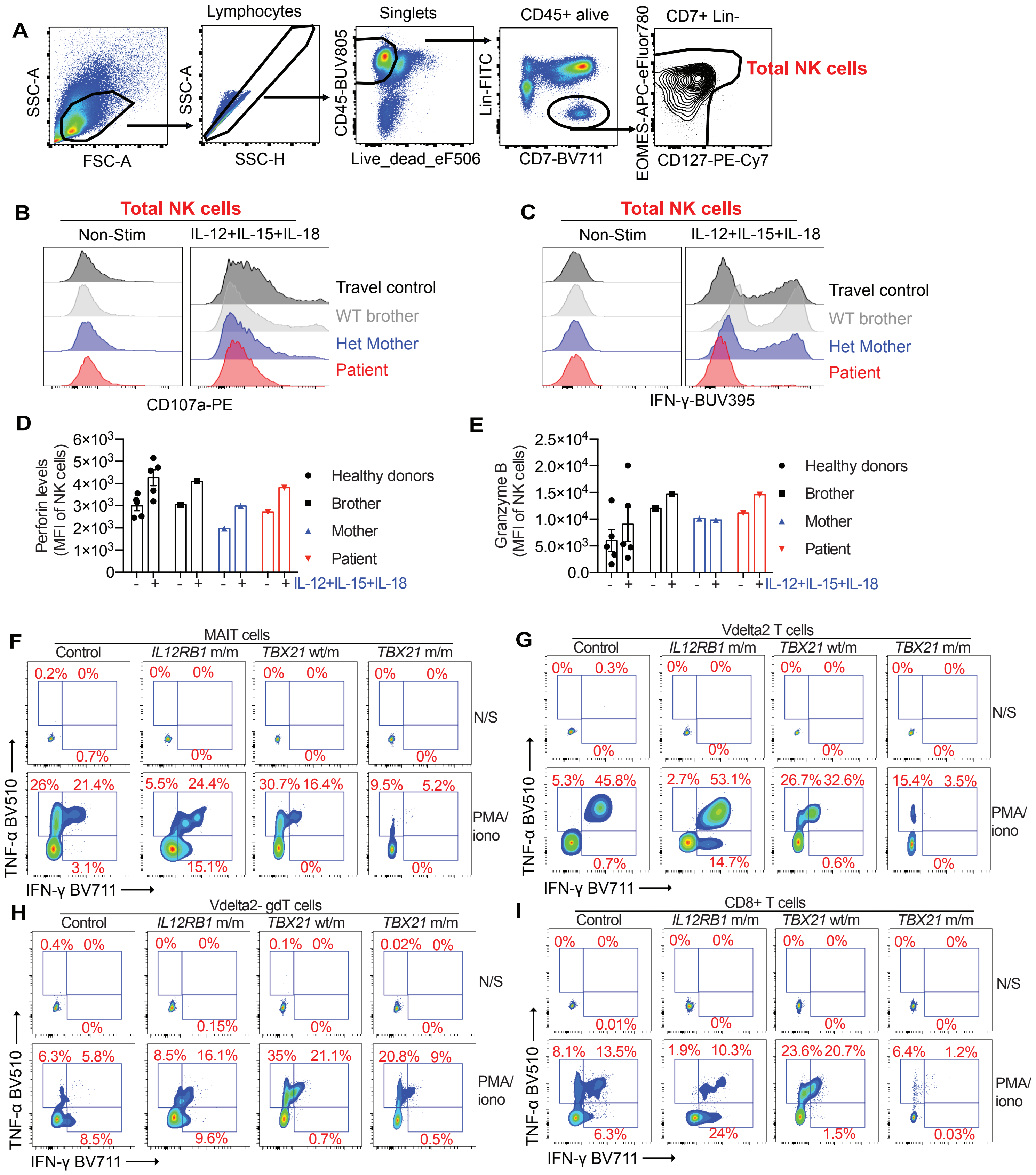

A
Lymphocytes
Singlets
CD45+ alive
CD7+ Lin-
Total NK cells
SSC-A
FSC-A
SSC-H
CD45-BUV805
Live_dead_eF506
Lin-FITC
CD7-BV711
EOMES-APC-eFluor780
CD127-PE-Cy7
B
Total NK cells
Non-Stim
IL-12+IL-15+IL-18
Travel control
WT brother
Het Mother
Patient
CD107a-PE
C
IFN-γ-BUV395
D
Perforin levels (MFI of NK cells)
Healthy donors
Brother
Mother
Patient
IL-12+IL-15+IL-18
E
Granzyme B (MFI of NK cells)
F
MAIT cells
Control
IL12RB1 m/m
TBX21 wt/m
TBX21 m/m
N/S
PMA/ iono
TNF-α BV510
IFN-γ BV711
G
Vdelta2 T cells
H
Vdelta2- gdT cells
I
CD8+ T cells


**Figure S16**

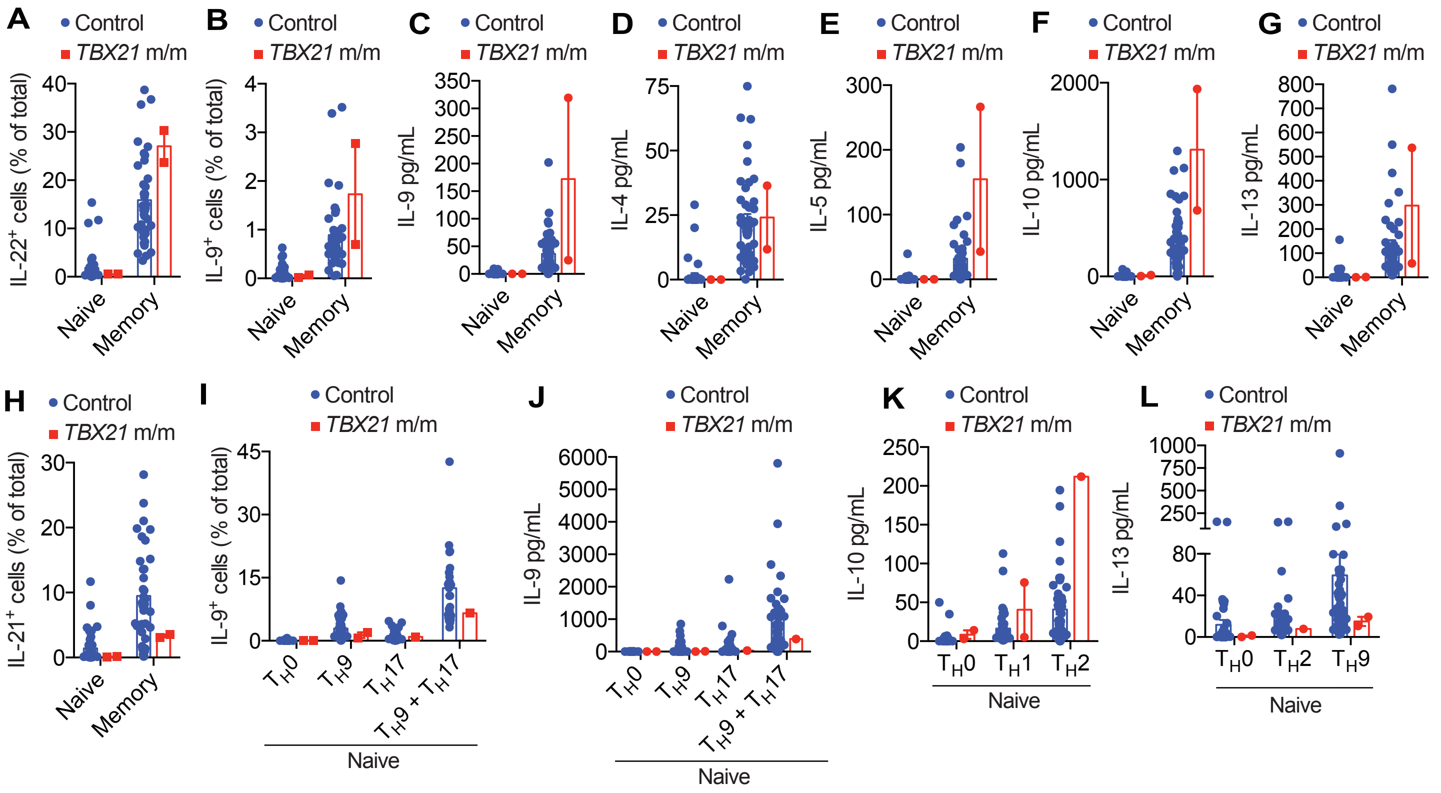
A
B
C
D
E
F
G
H
I
J
K
L
Control
TBX21 m/m
IL-22+ cells (% of total)
IL-9+ cells (% of total)
IL-9 pg/mL
IL-4 pg/mL
IL-5 pg/mL
IL-10 pg/mL
IL-13 pg/mL
IL-21+ cells (% of total)
Naive
Memory
TH0
TH9
TH17
TH9 + TH17
TH1
TH2

**Figure S17**

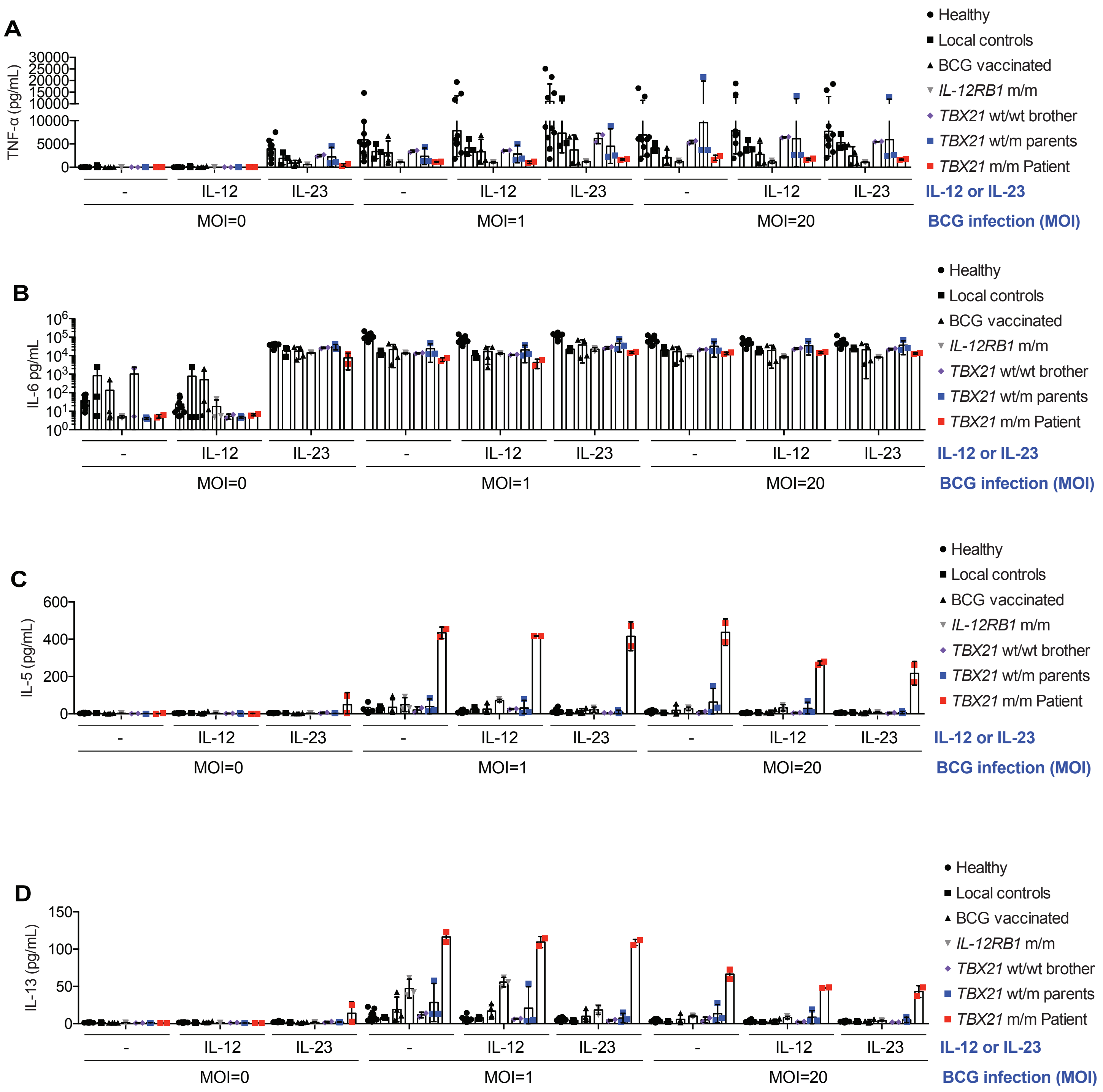
A
TNF-α (pg/mL)
30000
25000
20000
15000
10000
10000
5000
0
B
IL-6 pg/mL
$10^6$
$10^5$
$10^4$
$10^3$
$10^2$
$10^1$
$10^0$
C
IL-5 (pg/mL)
600
400
200
0
D
IL-13 (pg/mL)
150
100
50
0
-
IL-12
IL-23
MOI=0
MOI=1
MOI=20
IL-12 or IL-23
BCG infection (MOI)
Healthy
Local controls
BCG vaccinated
*IL-12RB1* m/m
*TBX21* wt/wt brother
*TBX21* wt/m parents
*TBX21* m/m Patient

**Figure S18**

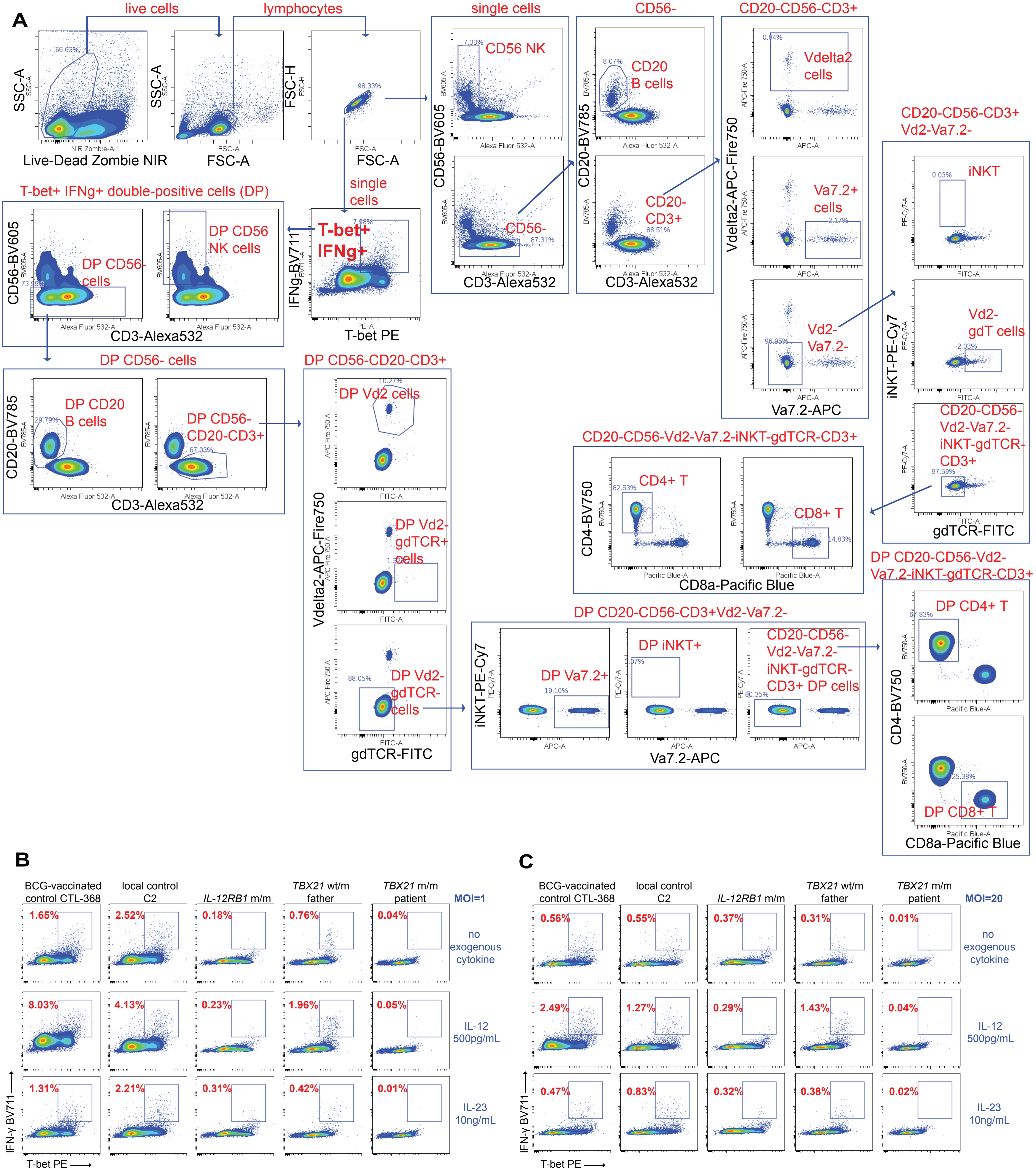
A
live cells
lymphocytes
single cells
CD56-
CD20-CD56-CD3+
66.63%
SSC-A
Live-Dead Zombie NIR
FSC-A
98.33%
FSC-H
FSC-A
7.33%
CD56 NK
CD56-BV605
8.07%
CD20
B cells
CD20-BV785
0.84%
Vdelta2
cells
CD56-
87.31%
CD3-Alexa532
CD20-
CD3+
88.51%
CD3-Alexa532
Va7.2+
cells
2.17%
Vdelta2-APC-Fire750
96.95%
Vd2-
Va7.2-
Va7.2-APC
CD20-CD56-CD3+
Vd2-Va7.2-
0.03%
iNKT
Vd2-
gdT cells
2.03%
CD20-CD56-
Vd2-Va7.2-
iNKT-gdTCR-
CD3+
97.59%
iNKT-PE-Cy7
gdTCR-FITC
T-bet+ IFNg+ double-positive cells (DP)
single
cells
7.98%
T-bet+
IFNg+
IFNg-BV711
T-bet PE
DP CD56
NK cells
DP CD56-
cells
73.99%
CD56-BV605
CD3-Alexa532
DP CD56- cells
DP CD20
B cells
29.79%
DP CD56-
CD20-CD3+
67.03%
CD20-BV785
CD3-Alexa532
DP CD56-CD20-CD3+
10.27%
DP Vd2 cells
DP Vd2-
gdTCR+
cells
88.05%
DP Vd2-
gdTCR-
cells
Vdelta2-APC-Fire750
gdTCR-FITC
CD20-CD56-Vd2-Va7.2-iNKT-gdTCR-CD3+
82.53%
CD4+ T
CD8+ T
14.83%
CD4-BV750
CD8a-Pacific Blue
DP CD20-CD56-CD3+Vd2-Va7.2-
DP Va7.2+
19.10%
DP iNKT+
0.07%
CD20-CD56-
Vd2-Va7.2-
iNKT-gdTCR-
CD3+ DP cells
80.35%
iNKT-PE-Cy7
Va7.2-APC
DP CD20-CD56-Vd2-
Va7.2-iNKT-gdTCR-CD3+
67.83%
DP CD4+ T
25.38%
DP CD8+ T
CD4-BV750
CD8a-Pacific Blue
B
BCG-vaccinated control CTL-368
local control C2
IL-12RB1 m/m
TBX21 wt/m father
TBX21 m/m patient
MOI=1
1.65%
2.52%
0.18%
0.76%
0.04%
no exogenous cytokine
8.03%
4.13%
0.23%
1.96%
0.05%
IL-12 500pg/mL
1.31%
2.21%
0.31%
0.42%
0.01%
IL-23 10ng/mL
IFN-γ BV711
T-bet PE
C
BCG-vaccinated control CTL-368
local control C2
IL-12RB1 m/m
TBX21 wt/m father
TBX21 m/m patient
MOI=20
0.56%
0.55%
0.37%
0.31%
0.01%
no exogenous cytokine
2.49%
1.27%
0.29%
1.43%
0.04%
IL-12 500pg/mL
0.47%
0.83%
0.32%
0.38%
0.02%
IL-23 10ng/mL
IFN-γ BV711
T-bet PE

**Figure S19**

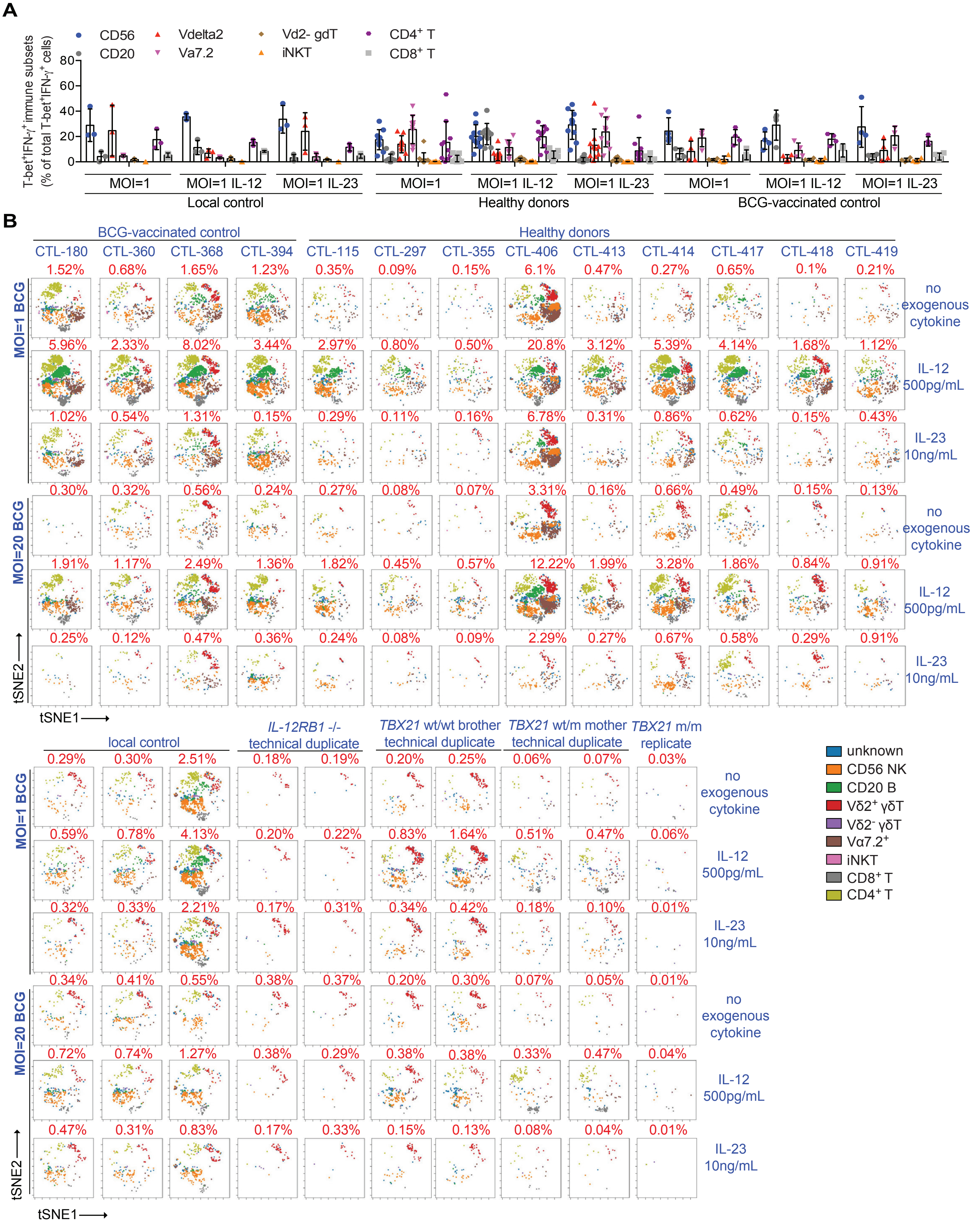

A
CD56
CD20
Vdelta2
Va7.2
Vd2- gdT
iNKT
CD4⁺ T
CD8⁺ T
T-bet⁺IFN-γ⁺immune subsets (% of total T-bet⁺IFN-γ⁺ cells)
0 20 40 60 80
MOI=1 MOI=1 IL-12 MOI=1 IL-23
Local control
MOI=1 MOI=1 IL-12 MOI=1 IL-23
Healthy donors
MOI=1 MOI=1 IL-12 MOI=1 IL-23
BCG-vaccinated control
B
BCG-vaccinated control
Healthy donors
CTL-180 CTL-360 CTL-368 CTL-394 CTL-115 CTL-297 CTL-355 CTL-406 CTL-413 CTL-414 CTL-417 CTL-418 CTL-419
MOI=1 BCG
1.52% 0.68% 1.65% 1.23% 0.35% 0.09% 0.15% 6.1% 0.47% 0.27% 0.65% 0.1% 0.21%
no exogenous cytokine
5.96% 2.33% 8.02% 3.44% 2.97% 0.80% 0.50% 20.8% 3.12% 5.39% 4.14% 1.68% 1.12%
IL-12 500pg/mL
1.02% 0.54% 1.31% 0.15% 0.29% 0.11% 0.16% 6.78% 0.31% 0.86% 0.62% 0.15% 0.43%
IL-23 10ng/mL
MOI=20 BCG
0.30% 0.32% 0.56% 0.24% 0.27% 0.08% 0.07% 3.31% 0.16% 0.66% 0.49% 0.15% 0.13%
no exogenous cytokine
1.91% 1.17% 2.49% 1.36% 1.82% 0.45% 0.57% 12.22% 1.99% 3.28% 1.86% 0.84% 0.91%
IL-12 500pg/mL
0.25% 0.12% 0.47% 0.36% 0.24% 0.08% 0.09% 2.29% 0.27% 0.67% 0.58% 0.29% 0.91%
IL-23 10ng/mL
tSNE2
tSNE1
local control
IL-12RB1 -/- technical duplicate
TBX21 wt/wt brother technical duplicate
TBX21 wt/m mother technical duplicate
TBX21 m/m replicate
MOI=1 BCG
0.29% 0.30% 2.51% 0.18% 0.19% 0.20% 0.25% 0.06% 0.07% 0.03%
no exogenous cytokine
0.59% 0.78% 4.13% 0.20% 0.22% 0.83% 1.64% 0.51% 0.47% 0.06%
IL-12 500pg/mL
0.32% 0.33% 2.21% 0.17% 0.31% 0.34% 0.42% 0.18% 0.10% 0.01%
IL-23 10ng/mL
MOI=20 BCG
0.34% 0.41% 0.55% 0.38% 0.37% 0.20% 0.30% 0.07% 0.05% 0.01%
no exogenous cytokine
0.72% 0.74% 1.27% 0.38% 0.29% 0.38% 0.38% 0.33% 0.47% 0.04%
IL-12 500pg/mL
0.47% 0.31% 0.83% 0.17% 0.33% 0.15% 0.13% 0.08% 0.04% 0.01%
IL-23 10ng/mL
tSNE2
tSNE1
unknown
CD56 NK
CD20 B
Vδ2⁺ γδT
Vδ2⁻ γδT
Vα7.2⁺
iNKT
CD8⁺ T
CD4⁺ T


**Figure S20**

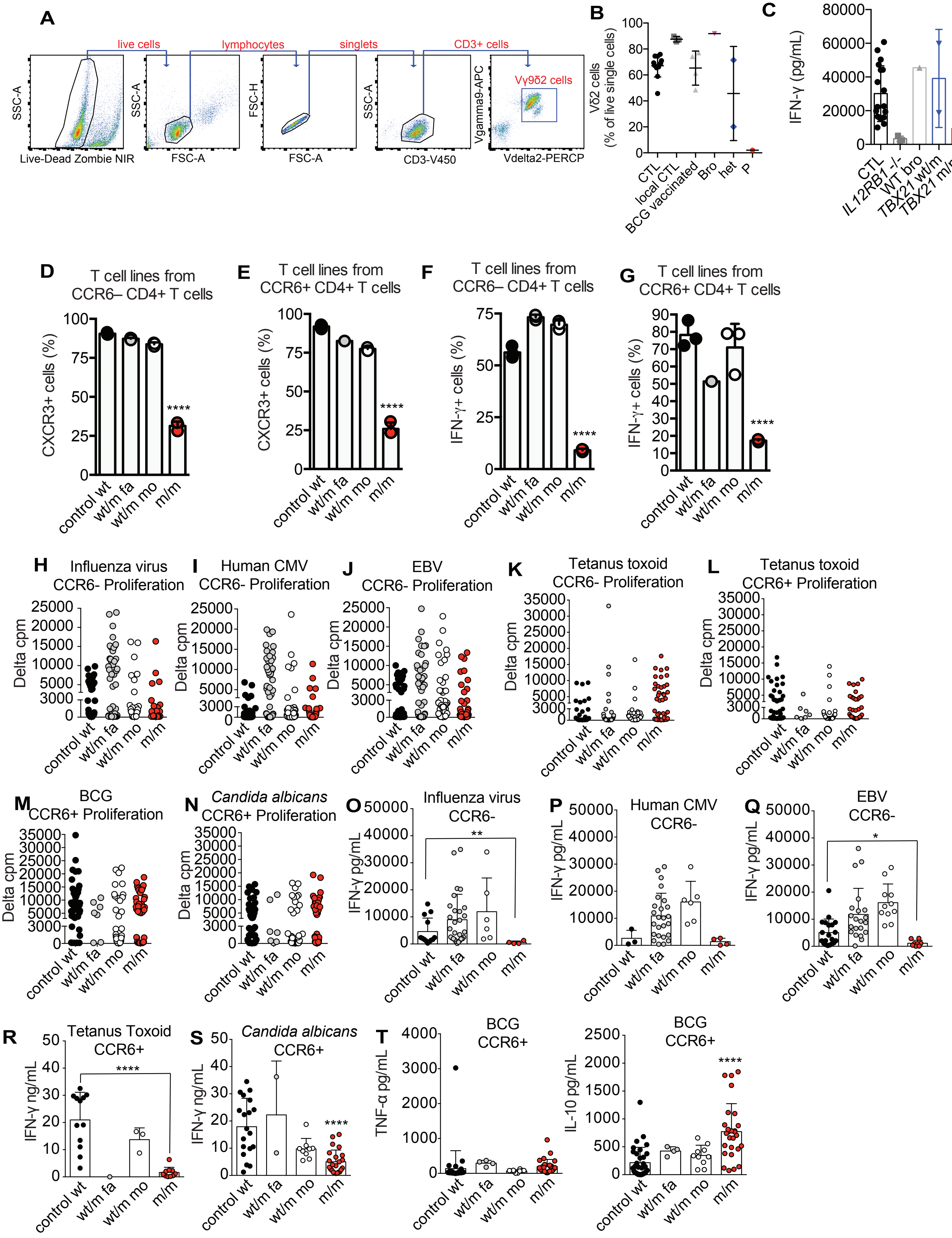

A
live cells
lymphocytes
singlets
CD3+ cells
Vγ9δ2 cells
SSC-A
Live-Dead Zombie NIR
FSC-A
FSC-H
CD3-V450
Vgamma9-APC
Vdelta2-PERCP
B
Vδ2 cells (% of live single cells)
CTL
local CTL
BCG vaccinated
Bro
het
P
C
IFN-γ (pg/mL)
CTL
IL12RB1 -/-
WT bro
TBX21 wt/m
TBX21 m/m
D
T cell lines from CCR6– CD4+ T cells
CXCR3+ cells (%)
E
T cell lines from CCR6+ CD4+ T cells
CXCR3+ cells (%)
F
T cell lines from CCR6– CD4+ T cells
IFN-γ+ cells (%)
G
T cell lines from CCR6+ CD4+ T cells
IFN-γ+ cells (%)
control wt
wt/m fa
wt/m mo
m/m
****
H
Influenza virus CCR6- Proliferation
I
Human CMV CCR6- Proliferation
J
EBV CCR6- Proliferation
K
Tetanus toxoid CCR6- Proliferation
L
Tetanus toxoid CCR6+ Proliferation
M
BCG CCR6+ Proliferation
N
Candida albicans CCR6+ Proliferation
Delta cpm
O
Influenza virus CCR6-
P
Human CMV CCR6-
Q
EBV CCR6-
IFN-γ pg/mL
R
Tetanus Toxoid CCR6+
S
Candida albicans CCR6+
IFN-γ ng/mL
T
BCG CCR6+
TNF-α pg/mL
BCG CCR6+
IL-10 pg/mL


**Figure S21**

**Figure legends for Supplementary Figures**

**Figure S1. Clinical and genetic features of a patient with MSMD, Related to Figure 1.** (**A**) Schematic view of the 15 MSMD-causing genes within myeloid and lymphoid cells in the context of host antimycobacterial immunity. **(B)** Serum IgG against various viral peptides or peptides derived from bacteria or other pathogens was tested by phage immunoprecipitation-sequencing. Serum samples from an unrelated healthy three-year-old boy (HC1), two healthy adults (HC2 and HC3), a *TBX21* m/m patient P, IgG-depleted serum and intravenous immunoglobulin (IVIg) were assayed. The heatmap (left) shows the counts of peptides significantly enriched for a given species with less than a continuous seven-residue subsequence, the estimated size of a linear epitope, in common. The species for which the patient had a score greater than our set threshold (>3) are shown. Count values above the set threshold are shown in on a color scale from green (⩾4) to red (⩾40) and dark-red (⩾80). **(C)** Whole-genome linkage (WGL) analysis of the kindred. Genomic regions with LOD scores (LOD score >1.3 and size >500 kb) are shown for each chromosome. The linked region of chromosome 17 containing *TBX21* is indicated. **(D)** Homozygous genetic variants in P were filtered based on: 1) their presence in P but not in siblings or parents, 2) predicted changes in the amino-acid composition of the encoded protein, 3) Rarity in the general population and major ethnic groups (MAF < 0.003, http://gnomad.broadinstitute.org and 1000 Genomes), 4) CADD score > MSC. **(E)** genomic DNA from P and his parents was subjected to Sanger sequencing to confirm the *TBX21* variant.

**Figure S2. The *TBX21* variant led to the substitution of two amino acids, Related to Figure 1. (A)** Comparison between the WT and Mut sequence of the region in which the variant is located in exon 1 of *TBX21*. An alignment of the WT and Mut sequences revealed a juxtaposed region with a sequence identical to that of the variant sequence. **(B** and **C)** The cDNAs generated from EBV-B cells derived from three healthy donors, *TBX21* wt/m heterozygous individuals (heterozygous) and the *TBX21* m/m patient (homozygous) were used as templates for the amplification of a region spanning the mutant site. Three sets of primers were used for this purpose (F forward primer and R4/R5/R6 reverse primers), as indicated. Agarose gel electrophoresis was performed to characterize these PCR products (B). PCR products were subjected to Sanger sequencing for genotyping (C). **(D)** RNA was extracted from PBMCs from two healthy donors and P and used for cDNA synthesis with random hexamers. The region spanning exons 1 to 4 was amplified from the cDNA by PCR. The PCR products were barcoded and an adaptor was added for Nano MiSeq sequencing. Sequencing reads passing quality controls were aligned with the reference genome, with RNA-STAR aligner. The sequence alignment for the junction between exons 1 and 2 of *TBX21* is shown with Integrative Genome Viewer.

**Figure S3. Overexpression of the mutant *TBX21* cDNA, Related to Figure 1. (A)** Alignment of peptide fragments from human T-box domain-containing transcription factors. **(B)** pCMV6 plasmids containing EV, WT T-bet (WT), the K314R variant of T-bet (K314R), T-bet with a deletion of the T-box domain (T-del), and mutant T-bet (Mut) were used to transfect HEK293T cells. Cells were harvested and lysed, and the lysates were subjected to immunoblotting. T-bet expression was characterized with anti-T-bet 4B10 monoclonal Ab and anti-Flag Ab. Anti-

GAPDH antibody was used as an endogenous control. **(C)** Mutations of individual residues in the T-bet cDNA (E156S, M157L, E156A, and E157A) were generated in pCMV6 plasmids by site-directed mutagenesis. The plasmids were used to transfect HEK293T cells for overexpression. T-bet levels were measured by immunoblotting with anti-T-bet 4B10 Ab and anti-Flag Ab. Anti-GAPDH Ab was used as endogenous control. **(D)** The plasmids described in (C) were used to transfect HEK293T cells, together with the WT-TBRE firefly luciferase and *Renilla* luciferase pRL-SV40 plasmids. After 3 days, the luciferase signal was detected in a Dual-Glo assay. The bars represent the mean and the standard deviation. Dots represent individual samples or technical replicates.

**Figure S4. Overexpression of the mutant *TBX21* cDNA does not induce IFN-γ production, Related to Figure 1. (A)** Schematic diagram of the CRISPR/Cas9 design for disrupting the exon-intron junctions of *TBX21*. gRNA9 was designed to disrupt the end of exon 1 and gRNA15 was designed to disrupt the end of exon 3. **(B)** Lentiviruses generated from empty vector, gRNA9 or gRNA15-containing pLenti-Crispr-V2 plasmids were used to transduce NK-92 cells. Stably transduced cells were genotyped with a three-primer PCR protocol. **(C** and **D)** Three EV-transduced, gRNA9- or gRNA15-transduced single-cell clones with a stable KO were selected based on the genotyping described in (B). We determined the levels of *TBX21* mRNA (C) by reverse transcription-quantitative PCR (RT-qPCR). mRNA levels were normalized by the $2^{-\Delta\Delta CT}$ method. T-bet protein levels were determined by immunoblotting (D). **(E)** Single-cell CRISPR/Cas9-edited NK-92 clones were stimulated with 50 pg/mL IL-12 and 10 ng/mL IL-18 or were left untreated (NS). Intracellular IFN-γ production was determined by ICS flow cytometry. **(F)** T-bet-deficient or EV-transduced NK-92 single-cell clones were transduced with retroviruses generated from pLZRS-ires-ΔNGFR plasmids containing no insert (EV), or the WT, K314R or Mut *TBX21* cDNA with a C-terminal Flag-tag as previously described. T-bet proteins levels were determined by immunoblotting. GAPDH levels were determined as an endogenous control. **(G)** NK-92 cells transduced as described in (F) were stimulated with 50 pg/mL IL-12 and 10 ng/mL IL-18 or were left untreated. Intracellular IFN-γ production was determined by ICS flow cytometry. **(H)** Naïve $CD4^+$ T cells were expanded under $T_H0$ conditions, and were then transduced with retroviruses generated from pLZRS-ires-ΔNGFR plasmids containing no insert (EV), or the WT, or Mut *TBX21* cDNA with a C-terminal Flag-tag. Transduced cells were positive for the bicistronic marker CD271. Cells were surface-stained for CD271 and intracellularly stained for IFN-γ and TNF-α and subjected to flow cytometry analysis. Bars represent the mean and the standard deviation. Dots represent individual samples or technical replicates.

**Figure S5. Variants of *TBX21* common in the general population and all variants from in-house cohorts were functionally neutral, Related to Figure 1. (A)** Schematic representation of all variants found in the homozygous state at least once in the gnomAD database (http://gnomad.broadinstitute.org). **(B)** Overexpression of seven variants of *TBX21* (shown in A) for which at least one homozygous individual has been identified in the general population. The variants indicated were overexpressed by transfecting HEK293T cells with pCMV6 plasmids containing the variants concerned. Protein levels were assessed by western blotting. **(C)** All variants (including the only homozygous variant and 28 heterozygous variants) from the HGID in-house database, excluding the H33Q allele already tested in (B), are represented on the CADD-MAF graph. The minor allele frequency (MAF) in the general population is shown on the *x*-axis. The CADD score is shown on the *y*-axis. **(D)** A schematic representation of all variants from our in-house cohorts is shown in (C). **(E)** All in-house *TBX21* variants were tested for transcriptional activity in the WT-TBRE luciferase reporter assay. pCMV6 plasmids containing variants were used to transfect cells, together with the TBRE luciferase reporter plasmid, and the pRL-SV40 *Renilla* plasmid for luciferase activity measurement. Bars represent the mean and standard deviation. Dots represent technical replicates.

**Figure S6. The T-bet variant in patient-derived cells is loss-of-function, Related to Figure 2.** **(A)** $CD4^+$ T cells from healthy donors (HD), the heterozygous father (Fa) or the patient (P) were expanded with anti-CD3/CD28 antibody-coated beads. They were subjected to RT-qPCR for *TBX21* with two different probes. Data are displayed as $2^{-\Delta\Delta Ct}$ after normalization relative to *GUS* (endogenous control) expression, using the control mean as a calibrator. **(B)** *TBX21* expression was assessed in immortalized T cells from healthy donors (HD), the heterozygous father (Fa) or the patient (P) by RT-qPCR. **(C)** P-derived HVS-T cells (*TBX21* m/m) were retrovirally transduced with EV or WT *TBX21* containing a bicistronic CD271 surface marker. Along with HD and heterozygous *TBX21* wt/m HVS-T cells, they were analyzed for intracellular T-bet and IFN-γ production by flow cytometry ICS in the presence or absence of P/I stimulation. **(D)** Isolated $CD4^+$ T cells from a healthy donor, a *TBX21* wt/m parent and the *TBX21* homozygous m/m P were expanded with anti-CD3/CD28 antibody-coated beads under $T_H0$ or $T_H1$ conditions. After 7d, EV or WT T-bet plasmids were used to transduce P's cells in the presence of a bicistronic CD271 surface reporter. Cells were subjected to ICS for IFN-γ and TNF-α in response to P/I restimulation. The gating strategy for the experiment is shown. **(E and F)** Cells transduced as in (D) were isolated with anti-CD271 antibody-coated beads by MACS.

Together with non-transduced cells, they were restimulated with anti-CD3/CD28 antibody-coated beads, and their culture supernatants were subjected to ELISA for IFN-γ (E) and TNF-α (F). Bars represent the mean and the standard deviation. Dots represent individual samples or technical replicates.

**Figure S7. The production of T helper effectors in T-bet deficiency is T-bet-dependent, Related to Figure 2. (A)** Expanded CD4$^{+}$ $T_H0$ cells from healthy donors, a T-bet heterozygous parent (TBET_HET), the T-bet homozygous P (TBET_HOM), P's cells complemented with empty vector (TBET_HOM_EV) or P's cells complemented with WT T-bet (TBET_HOM_WT) were restimulated with anti-CD3/CD28 antibody-coated beads for 16 h. RNA was extracted from these cells and subjected to RNA-seq analysis. Genome-wide transcriptome profile of all genes differentially expressed relative to the mean value for of healthy donors, represented as a heat map. **(B)** Transcriptomic analysis, as in (A), revealed target genes up- or downregulated in conditions of T-bet deficiency (WT/WT-control > or < M/M- homozygous deficiency), this differential regulation being abolished by WT T-bet complementation (M/M+EV > or < M/M+WT). The numbers of differentially regulated targets with T-bet-dependent expression are shown. **(C)** Volcano plots of transcriptomic profile comparing T-bet_HOM with CTL, and T-bet_HOM+WT with T-bet_HOM+EV. **(D and E)** Transcriptomic profile of immune genes displaying differential expression relative to the control (healthy donors), shown as a heat map for fold-change in expression (D) and Z-score (E). **(F)** Volcano plots of the transcriptomic profiles of immune genes comparing T-bet_HOM with CTL, and T-bet_HOM+WT with T-bet_HOM+EV.

**Figure S8. Changes in chromatin accessibility in CD4$^+$ T cells in conditions of T-bet deficiency, Related to Figure 3. (A)** Expanded CD4 $T_H0$ cells from healthy donors (control), IL-12Rβ1-deficient MSMD patients (IL12_rb1), a T-bet-heterozygous parent (Tbet_WTm), the T-bet-homozygous P (Tbet_mm), P's cells complemented with empty vector (Tbet_mm_EV) or P's cells complemented with WT T-bet (Tbet_mm_WT) in the absence of stimulation were used to prepare an omni-ATAC-seq library. The library was then subjected to omni-ATAC-seq. Heat map of the genome-wide omni-ATAC-seq peaks called. **(B)** A pie chart showing the genomic annotation of ATAC-seq peaks with respect to known RefSeq genes. **(C)** Number of chromatin peak accessibility changes, as determined by comparisons of the indicated groups of samples. **(D - H)** Regions of immune genes in which chromatin accessibility was downregulated in conditions of T-bet deficiency, including regions within *IRF1* (D), *HDAC9* (E), *CYLD* (F), *PSEN1* (G) and *HERC5* (H).

**Figure S9. Regions of immune genes in which chromatin accessibility was downregulated in conditions of T-bet deficiency, Related to Figure 3. (A - I)** Chromatin accessibilities of regions within *IL23R* (A), *IRF8* (B), *GZMB* (C), *KLRD1* (D), *RUNX3* (E), *SLAMF7* (F), *CCL5* (G), *IRF4* (H) and *RBPJ* (I).

**Figure S10. T-bet deficiency results in a normal dendritic cell distribution, Related to Figure 4. (A)** PBMCs were immunophenotyped with CyTOF. The gating strategy for different immune subsets is indicated. **(B)** Gating strategy for dendritic cell subsets, including plasmacytoid dendritic cells (pDC), conventional dendritic cells of type 1 and type 2 (cDC1 and cDC2), based on CyTOF data. **(C - E)** Percentages of pDCs (C), cDC1 cells (D) and cDC2 (E) cells. Bars represent the mean and the standard deviation. Dots represent individual samples.

**Figure S11. T-bet deficiency impairs NK cell maturation, Related to Figure 4. (A)** CD117$^+$ ILCs, consisting principally of ILC precursors (ILCP) and type 2 ILC (ILC2) cells, were gated manually on conventional flow cytometry. PBMCs from P or a healthy donor control were subjected to FACS immunophenotyping of innate lymphoid cells (ILCs) and NK cells. The gating strategy is shown. **(B)** Total NK cells (CD16$^+$ or CD94$^+$Lin$^-$CD7$^+$ cells) were plotted with CD56 against CD127 for manual gating on different NK cell subsets.

**Figure S12. T-bet deficiency does not affect ILC homeostasis in human PBMCs, Related to Figure 4. (A)** CD117$^+$ ILCs, consisting primarily of ILC precursors (ILCP) and type 2 ILC (ILC2) cells, were gated manually on CyTOF data. **(B** and **C)** Percentage of CD117$^+$ ILCs (B) and ILC2 cells (C) in healthy donors (CTL), IL-12Rβ1-deficient MSMD patients (12RB1$^{-/-}$) and the T-bet-deficient patient (P). **(D)** Immunophenotyping of ILC2 and ILCP cells, analyzed by fluorescence flow cytometry, as in Fig. S11. Plot of ILC2 and ILCP, and the percentage of ILC2 and ILCP (within live CD45$^+$ cells) in a healthy donor, the T-bet deficient patient and his healthy brother. Bars represent the mean and the standard deviation. Dots represent individual samples.

**Figure S13. T-bet deficiency results in the impaired development of iNKT, MAIT and Vδ2$^+$ cells, but normal Vδ1$^+$ γδT cell development, Related to Figure 4. (A)** PBMCs from healthy donors, IL-12Rβ1 deficient MSMD patients, the *TBX21* wt/m heterozygous parents, and the *TBX21* m/m patient were stained with surface markers for lymphocytes involved innate immunity (MAIT and iNKT cells) or both adaptive and innate immunity (γδT cells, in particular Vδ1 and Vδ2 cells). The gating strategy for flow cytometry is shown. **(B and C)** iNKT (B) and

MAIT (C) cells are shown for the T-bet-deficient patient and his family members. **(D)** Percentages of total γδT cells, gated on CyTOF data. **(E)** V$\delta 2^+$ γδT cells in the T-bet deficient patient and his family members. **(F)** Percentages of Vδ1 γδT cells in healthy donors (CTL), IL-12Rβ1-deficient MSMD patients (12RB1$^{-/-}$), the T-bet-deficient patient (P), and P's healthy heterozygous parents (het). **(G)** Naïve (CD45RA$^+$CCR7$^+$), central memory (CM, CD45RA$^-$CCR7$^+$), effector memory (EM, CD45RA$^-$CCR7$^-$) and effector memory T cells ($T_{EMRA}$, CD45RA$^+$CCR7$^-$) were manually gated from CD8$^+$ T cells on CyTOF data. The percentages of these cells are shown. **(H)** Total CD8$^+$ T cells were clustered by viSNE, based on a selection of surface markers. Manual gating of naïve and memory subsets overlaid on clusters of CD8$^+$ T cells. CXCR3 and CD38 expression are indicated on this plot. Bars represent the mean and the standard deviation (A and G) or the mean and the standard error of the mean (D). Dots represent individual samples.

**Figure S14. Preserved $T_H$2, $T_H$17 and circulating $T_{FH}$ memory CD4$^+$ T cells and normal Treg cells in T-bet deficiency, Related to Figure 4. (A)** Naïve (CD45RA$^+$CCR7$^+$), central memory (CM, CD45RA$^-$CCR7$^+$), effector memory (EM, CD45RA$^-$CCR7$^-$) and effector memory T cells ($T_{EMRA}$, CD45RA$^+$CCR7$^-$) were manually gated from CD4$^+$ T cells on the basis of CyTOF data. The percentages of these cells are shown. **(B)** Memory CD4$^+$ T cells (CD45RA$^-$) were gated out manually, and further subjected to viSNE deconvolution with T-cell chemokine markers for 5,000 randomly selected events. Expression of CXCR3, CCR5, CCR6 and CCR4. The gating strategy for memory CD4$^+$ T cells and $T_H$1, $T_H$2, $T_H$17 and $T_H$1* cells is shown. **(C** and **D)** Percentages of $T_H$2 (CCR6$^-$CXCR3$^-$CCR4$^+$) (C) and $T_H$17 cells (CCR6$^+$CXCR3$^-$CCR4$^+$) (D) among memory CD4$^+$ T cells. **(E)** Frequencies of the circulating follicular T helper ($cT_{FH}$)

cells and their subsets ($CXCR3^+$, double-negative as DN, $CCR6^+$, or $CXCR3^+CCR6^+$ $T_{FH}$) from healthy donors and the T-bet-deficient patient (*TBX21* m/m). **(F)** Percentage of Tregs. **(G)** Percentages of $CXCR3^+$ and $CCR5^+CXCR3^-$ Tregs. Bars represent the mean and the standard deviation (A, C, D, and F) or the mean and the standard error of the mean (E and G). Dots represent individual samples.

**Figure S15. Single-cell RNA-seq on human T-bet-deficient cells, Related to Figure 4. (A)** Single-cell RNA-seq UMAP clustering of PBMC from the index P (*TBX21* m/m) and his father (*TBX21* wt/m – Control 1). Major cell lineage clusters are labeled. **(B)** Cells expressing genes of interest are highlighted in red in UMAP clusters. *ANXA1*, *C15orf48*, *CCDC50*, *CST3*, *CTSW*, *CXCR3*, *CCL3L1*, *CCL4*, *CCL5*, *TNF*, *TRDC* and *WHRN* were selected as examples.

**Figure S16. *Ex vivo* stimulation of the residual lymphocytes from the patient with T-bet deficiency, Related to Figure 5. (A)** PBMCs from the patient or controls (healthy donors, brother-*TBX21* wt/wt, mother-*TBX21* wt/m) were stimulated with IL-12, IL-15 and IL-18. The gating strategy for total NK cells is shown. **(B** and **C)** Intracellular staining of IFN-γ (B) and TNF-α (C) in total NK cells from a travel control, P, P's brother and P's heterozygous mother, as shown in (A), in the presence and absence of stimulation with IL-12, IL-15 and IL-18. **(D** and **E)** Intracellular staining of perforin and granzyme B in total NK cells from a travel control, P, P's brother and P's heterozygous mother, as shown in (A). Mean fluorescence intensity (MFI) for perforin (D) and granzyme B (E) for the various samples. **(F - I)** MAIT cells ($CD56^-CD20^-CD3^+V\delta2^-V\alpha7.2^+$), Vδ2 cells ($CD56^-CD20^-CD3^+V\alpha7.2^-V\delta2^+$), Vδ2-negative γδT ($CD56^-CD20^-CD3^+V\alpha7.2^-V\delta2^-\gamma\delta TCR^+$) and $CD8^+$ T cells ($CD56^-CD20^-CD3^+V\alpha7.2^-V\delta2^-\gamma\delta TCR^+CD4^-CD8^+$)

were gated from PBMCs prepared as in (A). Intracellular staining for IFN-γ and TNF-α in these MAIT cells (Vα7.2$^+$) (F), Vδ2$^+$ cells (Vδ2$^+$) (G), Vδ2$^-$γδTCR$^+$ T cells (H) and CD8$^+$ T cells (I). Dots represent individual samples. Bars represent the mean and the standard error of the mean.

**Figure S17. *Ex vivo* and *in vitro* stimulation of CD4$^+$ T cells in conditions of T-bet deficiency, Related to Figure 5. (A - H)** Naïve and memory CD4$^+$ T cells were sorted by FACS and activated/expanded. Intracellular staining of IL-22 (A), IL-9 (B) and IL-21 (H). Production of IL-9 (C), IL-4 (D), IL-5 (E), IL-10 (F) and IL-13 (G), as measured in culture supernatants. **(I and J)** Naïve CD4$^+$ T cells were isolated by FACS and subjected to activation under $T_H0$, $T_H9$, $T_H17$ or $T_H9$ conditions combined with $T_H17$ conditions. The production of IL-9 was assessed intracellularly (I) and in culture supernatants (J). **(K)** Naïve CD4$^+$ T cells were isolated by FACS and subjected to activation under $T_H0$, $T_H1$ or $T_H2$ conditions IL-10 production, measured on culture supernatants. **(L)** Naïve CD4$^+$ T cells were isolated by FACS isolated and subjected to activation under $T_H0$, $T_H2$ or $T_H9$ conditions. IL-13 production, as measured on culture supernatants. Dots represent individual samples. Bars represent the mean and the standard error of the mean.

**Figure S18. T-bet deficiency leads to changes in cytokine production in response to *M.bovis* BCG stimulation, Related to Figure 6. (A - D)** PBMCs from healthy donors, local controls, BCG-vaccinated controls, IL-12Rβ1-deficient MSMD patients (*IL12RB1* m/m), the T-bet-deficient patient (*TBX21* m/m), and his family members were stimulated with or without live *M. bovis* BCG in the presence and absence of IL-12 and IL-23. The production of TNF-α (A), IL-6

(B), IL-5 (C) and IL-13 (D) was assessed with Legendplex cytometric bead arrays. Dots represent individual samples. Bars represent the mean and the standard deviation.

**Figure S19. T-bet deficiency leads to smaller numbers of T-bet$^+$ IFN-$\gamma^+$ double-positive cells following stimulation with live *M. bovis* BCG, Related to Figure 6. (A)** PBMCs from healthy donors, local controls, BCG-vaccinated controls, IL-12Rβ1-deficient MSMD patients (*IL12RB1* m/m), the T-bet-deficient patient (*TBX21* m/m), and his family members were stimulated with live *M. bovis* BCG in the presence and absence of IL-12 and IL-23. The gating strategies for total CD56$^+$ NK cells, CD20$^+$ B cells, Vδ2$^+$ γδT cells, Vα7.2$^+$ MAIT cells, Vδ2$^-$ γδT cells, iNKT cells, CD4$^+$ T cells and CD8$^+$ T cells are shown. The gating strategies for T-bet$^+$ IFN-$\gamma^+$ double-positive CD56$^+$ NK cells, CD20$^+$ B cells, Vδ2$^+$ γδT cells, Vα7.2$^+$ MAIT cells, Vδ2$^-$ γδT cells, iNKT cells, CD4$^+$ T cells and CD8$^+$ T cells are also shown. **(B** and **C)** Plots and percentages of T-bet$^+$ IFN-$\gamma^+$ double-positive cells from the individuals indicated in response to live *M. bovis* BCG infection at a MOI=1 (B) and a MOI=20 (C) in the presence and absence of IL-12 or IL-23.

**Figure S20. T-bet deficiency leads to defective Vδ2$^+$ γδT and MAIT cells, resulting in susceptibility to mycobacteria, Related to Figure 6. (A)** PBMCs from healthy donors, local controls, BCG-vaccinated controls, IL-12RB1-deficient MSMD patients (IL-12Rβ1 m/m), the T-bet-deficient patient (T-bet m/m), and his family members were stimulated with live *M. bovis* BCG in the presence and absence of IL-12 and IL-23. The percentages of CD56$^+$ NK cells, CD20$^+$ B cells, Vδ2$^+$ γδT cells, Vα7.2$^+$ MAIT cells, Vδ2$^-$ γδT cells, iNKT cells, CD4$^+$ T cells and CD8$^+$ T cells among T-bet$^+$ IFN-$\gamma^+$ double-positive cells are shown for healthy donors, local controls and BCG-vaccinated controls. **(B)** T-bet$^+$ IFN-$\gamma^+$ double-positive cells among PBMCs from healthy donors, local controls, BCG-vaccinated controls, an IL-12Rβ1-deficient MSMD patient (*IL12RB1*$^{-/-}$), the *TBX21* m/m patient and his family members in response to *M. bovis* BCG stimulation, deconvoluted into automated clusters by viSNE analysis. Clusters representing CD56$^{+}$ NK cells, CD20$^{+}$ B cells, Vδ2$^{+}$ γδT cells, Vα7.2$^{+}$ MAIT cells, Vδ2$^{-}$ γδT cells, iNKT cells, CD4$^{+}$ T cell and CD8$^{+}$ T cells are shown. Dots represent individual samples. Bars represent the mean and the standard deviation.

**Figure S21. Defective prolonged anti-BCG immunity in Vδ2 γδ T but not memory CD4$^{+}$ T cells in T-bet deficiency, Related to Figure 6. (A)** PBMCs from healthy controls, IL-12Rβ1-deficient patients, the T-bet wt/m parents (mother and father), the T-bet wt/wt brother, and the patient (T-bet m/m) were stimulated with lysates of *M. bovis* BCG for 14 days. The gating strategy for the analysis of Vγ9δ2 T cells, also known as Vδ2 cells, is shown. (**B**) Percentages of Vγ9δ2 T cells among live single cells in culture, as in (A). **(C)** IFN-γ production by the end of the 2two-week culture period, as in (A), was measured in Legendplex assays. **(D - G)** T-cell libraries from a healthy BCG-vaccinated donor (control wt), the T-bet wt/m parents (wt/m), and the T-bet m/m patient (m/m) were generated by the polyclonal stimulation of sorted memory CD4$^{+}$ T cells (either CCR6$^{-}$ or CCR6$^{+}$). The proportions of CXCR3$^{+}$ (D and E) and IFN-γ$^{+}$ (F and G) cells were determined by surface staining and ICS flow cytometry. **(H - N)** Libraries from CCR6$^{-}$ (H - K) or CCR6$^{+}$ (L - N) memory CD4$^{+}$ T cells from a control (wt control), the T-bet wt/m parents (wt/m), and the T-bet m/m patient (m/m) were screened for antigen reactivity by culturing T cells with autologous antigen-presenting cells, with and without peptide pools derived from influenza virus (H), human CMV (I), EBV (J), tetanus toxoid (K and L), *M. bovis* BCG (M) and *C. albicans* (N). Proliferation was assessed by evaluating [$^{3}$H]-thymidine incorporation, and is expressed in delta cpm values. **(O - Q)** CCR6$^-$ memory CD4$^+$ T cell clones responsive to peptide pools from influenza virus (O), human CMV (P) or EBV (Q) were subjected to Luminex assays to assess the production of IFN-γ. **(R and S)** CCR6$^+$ memory CD4$^+$ T-cell clones responsive to peptide pools from tetanus toxoid (R) and *C. albicans* (S) were subjected to Luminex assays to assess the production of IFN-γ. **(T)** BCG-responsive T-cell clones, as in (M), were subjected to Luminex assays to assess the production of TNF-α and IL-10. Dots represent individual samples or T-cell clones. Bars represent the mean and the standard deviation. One-way ANOVA was used to analyze statistical significance for D, E, F, G, S and T. A non-parametric *t*-test comparing WT to m/m was used to analyze statistical significance for (O - R).

**Table S1: Overview of diseases underlying isolated or syndromic Mendelian susceptibility to mycobacterial disease (MSMD)**

| Gene | Inheritance | Defect | Protein |
|---|---|---|---|
| *IL12RB1* | AR | Complete | E- |
| | AR | Complete | E+ |
| *IL12B* | AR | Complete | E- |
| *IL23R* | AR | Complete | E+ |
| *IL12RB2* | AR | Complete | E- |
| *SPPL2A* | AR | Complete | E- or E+ |
| *IRF8* | AD | Partial | E+ |
| | AR | Complete | E- or E+ |
| *IFNGR1* | AR | Complete | E+ |
| | AR | Complete | E- |
| | AD | Partial | E+++ |
| | AR | Partial* | E+ |
| | AR | Partial | E+ |
| *IFNGR2* | AR | Complete | E+ |
| | AR | Complete | E- |
| | AR | Partial | E+ of mutant protein |
| | AR | Partial | E+ of WT protein |
| | AD | Partial | E+ |
| *STAT1* | AD | Partial | E+P- |
| | AD | Partial | E+B- |
| | AD | Partial | E+P-B- |
| | AR | Complete | E- |
| | AR | Partial | E+ |

| | | | |
|---|---|---|---|
| *NEMO* (*IKBKG*) | XR | Partial | E+ |
| *CYBB* | XR | Partial* | E+ |
| *TYK2* P1104A | AR | Partial | E+ |
| *TYK2* | AR | Complete | E- |
| *JAK1* | AR | Partial | E- |
| *RORC* | AR | Complete | E- |
| *ISG15* | AR | Complete | E- |

MSMD genetic etiologies may have autosomal recessive (AR), autosomal dominant (AD), or X-linked recessive (XR) inheritance. Defects can be complete (C) or partial (P). Protein expression (E) of the mutant can be abolished (E-), normal or weaker than normal (E+), or stronger than normal (E+++). The mutant protein may be able or unable to undergo phosphorylation (P-) or to bind to DNA (B-). * Impaired function in specific cell types.

| | No. of patients diagnosed | αβ $CD4^+$ T | αβ $CD8^+$ T | MAIT | B cell/Ig | DC | NK | iNKT | Mycobacterial infection |
|---|---|---|---|---|---|---|---|---|---|
| CD8 deficiency (*CD8A*) | 3 | Nl | A | | Nl | | | | No (OMIM#186910) (Dumontet et al., 2015; de la Calle-Martin et al., 2001; Mancebo et al., 2008). |
| MHC class I deficiency (*TAP1*, *TAP2*, *TAPBP*, *B2M*) | ~30 | Nl | ↧↧↧↧ | | Nl | | | | No (OMIM#170260, OMIM#170261, OMIM#601962, OMIM#109700) (Ardeniz et al., 2015; Chen et al., 1996; Colonna et al., 1992; Furukawa et al., 1999; Gao et al., 2016; Hanalioglu et al., 2017; Hanna and Etzioni, 2014; De La Salle et al., 1994; Law-Ping-Man et al., 2018; Maeda et al., 1985; Moins-Teisserenc et al., 1999; Sophie et al., 2012; Wani et al., 2006; Yabe et al., 2002). |
| TCRα deficiency | 1 | A | A | | Nl | | | | No (OMIM#615387) (Morgan et al., 2011). |
| MHC class II deficiency (*CIITA*, *RFXANK*, *RFX5*, *RFXAP*) | >200 | ↧↧↧↧ | Nl | | ↧↧ | | | | Generally no (OMIM#209920) (Bontron et al., 1997; Durand et al., 1997; Dziembowska et al., 2002; Ghaderi et al., 2006; Hanna and Etzioni, 2014; Lennon-Duménil |

| | | | | | | | | | |
|---|---|---|---|---|---|---|---|---|---|
| | | | | | | | | | et al., 2001; Lisowska-Grospierre et al., 1994; Mach et al., 1998; Nagarajan et al., 2000; Nekrep et al., 2002; Ouederni et al., 2011; Peijnenburg et al., 1999; Steimle et al., 1993, 1995; Swanberg et al., 2005; Villard et al., 1997a, 1997b, 2001, 2002; Wiszniewski et al., 2000, 2003). One case of *M. avium* infection reported (Dimitrova et al., 2014). |
| MCM4 deficiency (*MCM4*) | ~24 | Nl | Nl | | Nl | | ↧↧↧↧ | | No (OMIM#602638) (Casey et al., 2012; Eidenschenk et al., 2006; Gineau et al., 2012; Hughes et al., 2012). |
| GINS1 deficiency (*GINS1*) | 5 | Nl | Nl | | Nl | | ↧↧↧↧ | | No (OMIM#610608) (Cottineau et al., 2017). |
| RORγ/RORγT deficiency | 7 | ↧ | Nl | A | | | | A | Yes (OMIM#602943) (Okada et al., 2015). |
| AD IRF8 deficiency | 4 | | | | | ↧↧ to A | ↧↧ | | Yes (OMIM#601565) (Hambleton et al., 2011; Salem et al., 2014). |
| SPPL2A deficiency (*SPPL2A*) | 2 | | | | | ↧↧ | | | Yes (Kong et al., 2018). |

| | | | | | | | | | |
|---|---|---|---|---|---|---|---|---|---|
| SAP deficiency | >300 | | | | | | | ↧↧↧↧ | No (OMIM#308240) reviewed in (Tangye et al., 2017). |
| WASP deficiency | Many | | | | | | | A | Generally No (OMIM#300392) reviewed in (Locci et al., 2009; Massaad et al., 2013). Two cases of mycobacterial infection reported (Pacharn et al., 2017; Yasutomi et al., 2015). |

**Abbreviations:** A absent**,** ↧ reduced, ↥ elevated, Nl normal

**Table S2.** Mycobacterial diseases in primary immunodeficiency disorders in humans leading to the selective depletion of one or more populations of αβ T cells, NK cells, MAIT cells or dendritic cell subsets.

| Gene | Chr | Position | Ref | Alt | Consequence | Function category | Comments |
|---|---|---|---|---|---|---|---|
| *TSEN15* | 1 | 184041317 | G | A | p.Arg127Gln | Nucleic acid binding | MAF=0.00336 in Middle Eastern population (Joseph Gleeson cohort). MAF<0.003 in all other ethnic groups in public databases. |
| *EPHA1* | 7 | 143098636 | C | G | p.Met71Ile | Angiogenesis, GTPase activation, ephrin receptor signaling pathway | MAF<0.003 in all ethnic groups in public databases. |
| *AKR1D1* | 7 | 137776608 | T | C | p.Ile119Thr | Androgen metabolism, bile acid synthesis | MAF<0.003 in all ethnic groups in public databases. |
| *ABCA2* | 9 | 139908757 | TGGGTCAGCTCCGAGCACC | T | p.Arg1362_Thr1367del | Cholesterol homeostasis, lipid metabolism | MAF<0.003 in all ethnic groups in public databases. |
| *PDE2A* | 11 | 72297209 | C | T | p.Glu363Lys | GPCR signaling, cAMP-regulation | MAF<0.003 in all ethnic groups in public databases. |
| *PSMD9* | 12 | 122326874 | A | C | p.Lys38Gln | Insulin secretion, bHLH transcription factor binding | MAF<0.003 in all ethnic groups in public databases. |
| *GCN1L1* | 12 | 120568996 | G | A | p.Ala2519Val | Amino acid starvation, cadherin | MAF<0.003 in all ethnic groups in |

| | | | | | | | |
|---|---|---|---|---|---|---|---|
| | | | | | | binding, regulation of translation | public databases. |
| *ATP6V0A2* | 12 | 124228405 | G | A | p.Arg371His | Iron homeostasis, ATP hydrolysis coupled proton transport | MAF<0.003 in all ethnic groups in public databases. |
| *CIB2* | 15 | 78416081 | C | T | p.Asp18Asn | Calcium homeostasis | MAF<0.003 in all ethnic groups in public databases. |
| *ERN1* | 17 | 62121392 | C | T | p.Glu964Lys | JUN kinase activation, cell cycle arrest, insulin metabolism | Another patient with a different phenotype in laboratory cohort is homozygouse for the same variant. |
| *TBX21* | 17 | 45811289 | ATG | A | p.Glu156_Met 157SerLeu | T-helper regulation, T-cell differentiation | Private variant, and highly related to interferon gamma immunity |
| *AOC2* | 17 | 40997583 | G | T | p.Gly314Cys | Amine metabolism, catecholamine metabolism, visual perception | >2% Ashkenazi Jews carry a LOF premature stop variant in *AOC2* (Variant: 17-40996783-G-A). |
| *AKAP1* | 17 | 55182923 | T | G | p.Val33Gly | Blood coagulation, protein kinase A regulation | >6.4% of Africans carry a LOF premature stop variant |

| | | | | | | | |
|---|---|---|---|---|---|---|---|
| | | | | | | | of *AKAP1* (Variant: 17-55198645-C-CT) |
| *ARHGAP27* | 17 | 43506940 | G | A | p.Arg236Cys | GTPase regulation, receptor-mediated endocytosis | Another patient with a different phenotype in laboratory cohort is homozygous for the same variant. |
| *RHBDD3* | 22 | 29656330 | G | C | p.Ser323Cys | Liver development, MAPK cascade, serine-type endopeptidase activity | >0.79% of the general population (all gnomAD) carry a LOF premature stop variant of *RHBDD3* (Variant: 22-29656431-C-T) |

**Table S3.** Homozygous candidate variants that might cause MSMD and reactive airway disease in the patient.

| Target Gene | Distance | Rank | *P*-Value (percentile) | Sphere | Degrees of separation |
|---|---|---|---|---|---|
| *TBX21* | 1.11111 | 7 | 0.00042 | 0 | 1 |
| *EPHA1* | 4.44444 | 602 | 0.03597 | 2 | 2 |
| *PSMD9* | 4.44444 | 1040 | 0.06214 | 3 | 2 |
| *ERN1* | 4.72222 | 2132 | 0.12738 | 4 | 2 |
| *ABCA2* | 10.41667 | 7835 | 0.46812 | 5 | 3 |
| *GCN1L1* | 10.41667 | 7193 | 0.42977 | 5 | 3 |
| *ARHGAP27* | 11.98582 | 10149 | 0.60638 | 6 | 3 |
| *PDE2A* | 18.11722 | 14155 | 0.84573 | 7 | 4 |
| *CIB2* | 19.29372 | 14797 | 0.88409 | 7 | 3 |
| *ATP6V0A2* | 19.68254 | 15112 | 0.90291 | 7 | 4 |
| *TSEN15* | 19.88576 | 15163 | 0.90596 | 7 | 4 |
| *AKR1D1* | 27.3569 | 16308 | 0.97437 | 7 | 4 |

**Table S4.** Connectome analysis of the distance of *TBX21*, *EPHA1*, *PSMD9*, *ABCA2*, *GCN1L1*, *ERN1*, *ARHGAP27*, *PDE2A*, *CIB2*, *ATP6V0A2*, *TSEN15*, and *AKR1D1* to *IFNG* as the anchor gene.

| | **In human T-bet deficiency** | **In mouse T-bet deficiency** |
|---|---|---|
| **CD56$^{bright}$ NK cells** | ↓ 25-fold in frequency | ↓60-fold in frequency in periphery (Townsend et al., 2004) |
| **CD16$^{+}$CD56$^{dim}$ NK cells** | ↓ 15-fold in frequency | |
| **Invariant NKT cells** | ↓ 200-fold in frequency | ↓18-fold in frequency in liver (Townsend et al., 2004) |
| **MAIT cells** | ↓ 14-fold in frequency<br>Low levels of IFN-γ production in the remaining cells** | Not known |
| **$T_{H}$1 CD4$^{+}$ T cells** | ↓ 9-fold in frequency<br>Low levels of IFN-γ production in the remaining cells** | ↓ in frequency (Szabo et al., 2002) |
| **$T_{H}$1* CD4$^{+}$ T cells** | Normal in frequency | Not known |
| **Vδ2 γδT cells** | ↓ 8-fold in frequency<br>Low levels of IFN-γ production in the remaining cells** | Not present in mice |
| **Vδ1 γδT cells** | Normal in frequency<br>Normal IFN-γ production in Vδ1-containing γδT cells (Vδ2$^{-}$ γδT cells) ** | Not known |
| **$T_{H}$2 CD4$^{+}$ T cells** | Normal in frequency | ↑ $T_{H}$2 differentiation and cytokine production during experimental inoculation (Finotto et al., 2002) |
| **$T_{H}$17 CD4$^{+}$ T cells** | Normal in frequency | ↑ $T_{H}$17 differentiation and cytokine production during experimental inoculation (Lazarevic et al., 2011) |
| **CD8$^{+}$ T cells** | Normal in frequency<br>Low levels of IFN-γ production** | Normal in frequency (Tang et al.)<br>Low levels of IFN-γ production (Bettelli et al., 2004; Intlekofer et al., 2007; Pearce et al., 2003; Sullivan et al., 2003) |
| **ILC2** | Normal in frequency | ↑ ILC2 and ↑ activity of ILC2 in the spleen, lymph nodes and gut mucosa (Garrido-Mesa et al., 2019) |
| **ILCP** | Normal in frequency | Not known |
| **ILC1** | Cannot be studied as present at too low a frequency in peripheral blood | ↓ in frequency (Daussy et al., 2014) |
| **ILC3** | Cannot be studied as present at too low a frequency in peripheral blood | ↓ in frequency (Rankin et al., 2013) |

| | | |
|---|---|---|
| **mDC and pDC subsets** | Normal in frequency | Normal in frequency (Lugo-Villarino et al., 2003) |

**In response to PMA-ionomycin stimulation

**Table S8.** Comparison of the development and function of various immune lineages of human and murine T-bet-deficient cells.

| | Mice or Rats (T-bet KO or cKO or silencing) | Human T-bet deficiency (P) |
|---|---|---|
| Susceptibility to infections | ➢ ⥣ *Leishmania major* and *L. donovani* (Rosas et al., 2006; Szabo et al., 2002)<br>➢ ⥣ *Toxoplasma gondii* colonization at secondary sites of infection (Harms Pritchard et al., 2015)<br>➢ ⥣ Attenuated rabies virus CNS infection (Lebrun et al., 2015)<br>➢ ⥣ *Staphylococcus aureus* sepsis and arthritis (Hultgren et al., 2004)<br>➢ ⥣ Genital HSV-2 (Svensson et al., 2005)<br>➢ ⥣ *Mycobacterium tuberculosis* (Sullivan et al., 2005)<br>➢ ⥣ *M. avium complex* (MAC) (Matsuyama et al., 2014)<br>➢ ⥣ *Salmonella* infection (Ravindran et al., 2005)<br>➢ ⥣ *M. pulmonis* colonization (Bakshi et al., 2006)<br>➢ ⥣ Chronic LCMV infection (Kao et al., 2011)<br>➢ ⥣ *Plasmodium berghei* parasitemia (Oakley et al., 2013)<br>➢ ⥥ *Plasmodium berghei* experimental cerebral malaria (Oakley et al., 2013)<br>➢ ⥥ *Plasmodium yoelii* 17XNL infection<br>➢ ⥥ *T. spiralis* infection in intestines (Oakley et al., 2014)<br>➢ ⥥ *T. spiralis* infection in intestines (Alcaide et al., 2007) | ➢ ⥣ BCG infection<br>➢ Mild chronic CMV viremia (this publication) |

**Table S9.** Comparison of the infectious phenotypes of T-bet deficiency between humans and rodents.

**Legends to the supplemental tables**

**Table S1.** Overview of diseases underlying isolated or syndromic Mendelian susceptibility to mycobacterial disease (MSMD).

**Table S2.** Mycobacterial diseases in primary immunodeficiency disorders in humans leading to the selective depletion of one or more populations of αβ T cells, NK cells, MAIT cells or dendritic cell subsets.

**Table S3.** Homozygous candidate variants that might cause MSMD and reactive airway disease in P.

**Table S4.** Connectome analysis of the distances of *TBX21*, *EPHA1*, *PSMD9*, *ABCA2*, *GCN1L1*, *ERN1*, *ARHGAP27*, *PDE2A*, *CIB2*, *ATP6V0A2*, *TSEN15*, and *AKR1D1* to *IFNG,* the anchor gene.

**Table S5.** RNA-seq analysis of expanded CD4 $T_H0$ cells from healthy donors, a T-bet heterozygous parent (TBET_HET), the T-bet homozygous P (TBET_HOM), P complemented with empty vector (TBET_HOM_EV) or P complemented with WT T-bet (TBET_HOM_WT) were restimulated with anti-CD3/CD28 Ab-coated beads for 16 h. Lists of DEG_Status, DEG_FoldChange and DEG_ZScore identifying differentially regulated target genes genome-wide. Slim_DEG identifies differentially regulated target immune genes.

**Table S6.** Chromatin accessibility of 1649 loci opening in a T-bet-dependent manner and 666 loci closing in a T-bet-dependent manner.

**Table S7.** Beta-value for the differential methylation of CpG sites between T-bet m/m (Hom) and controls (CTL) abolished by WT T-bet complementation.

**Table S8.** Comparison of the development and function of various immune lineages of human and murine T-bet-deficient cells.

**Table S9.** Comparison of the infectious phenotypes of human T-bet deficiency in humans and rodents.
